# Cellular Aging Signatures in the Plasma Proteome Record Human Health and Disease

**DOI:** 10.64898/2026.02.10.704909

**Authors:** Daisy Yi Ding, Veronica Augustina Bot, Kenneth L Chen, James Groves, Róbert Pálovics, Daisuke Masuda, Amelia Farinas, Hamilton Se-Hwee Oh, Viktoria Wagner, Nannan Lu, The Global Neurodegeneration Proteomics Consortium (GNPC), Carlos Cruchaga, Alina Isakova, Jonathan M Schott, Tony Wyss-Coray

**Author notes:** These authors contributed equally: Daisy Yi Ding, Veronica Augustina Bot, Kenneth L Chen.

## Abstract

Aging is asynchronous across cells and organs, but whether plasma proteins can capture cell type-specific aging and predict disease and mortality remains unknown. We developed machine learning models to estimate the biological age of more than 40 distinct cell types—spanning neuronal, immune, glial, endocrine, epithelial, and musculoskeletal origins—using over 7,000 plasma proteins measured in 60,000 individuals across three cohorts, comprising the largest human plasma proteomics aging study to date. Individuals showed heterogeneous aging profiles, with 20-25% exhibiting accelerated aging in a single cell type and 1-3% across ten or more cell types. APOE genotype showed antagonistic aging effects in different cell types: APOE4 carriers exhibited older astrocytes but younger macrophages, while APOE2 carriers showed the inverse. Cellular aging signatures were uniquely associated with disease status and predicted incident disease and mortality over 15 years of follow-up. Amyotrophic lateral sclerosis (ALS) showed the strongest association with skeletal myocyte aging (hazard ratio = 12.7 for extreme accelerated versus youthful aging). In Alzheimer’s disease (AD), prevalent cases showed accelerated aging across multiple neural and peripheral cell types, with extreme astrocyte aging conferring AD risk comparable to APOE4 carrier status. Moreover, extreme astrocyte aging increased AD risk in APOE4/4 carriers threefold, while youthful astrocytes strikingly reduced risk. Beyond neurodegeneration, respiratory cell aging identified smokers at 58% higher lung cancer risk, and myeloid aging identified normoglycemic individuals at higher diabetes risk. Both specific cellular vulnerabilities and cumulative aging burden influenced survival, wherein youthful immune or neuronal profiles were protective. A polycellular aging risk score provided robust mortality risk stratification across platforms and cohorts. These findings establish a framework for quantifying biological aging at the cellular resolution using plasma proteomics, revealing heterogeneity in aging trajectories and their impact on disease susceptibility and resilience.

## MAIN

Over fifty percent of the global disease burden can be attributed to aging^1^. One’s risk of neurodegenerative disease, including Alzheimer’s disease (AD) and Parkinson’s disease (PD), increases twofold for each elapsed five calendar years, with the timing of onset being largely heterogeneous among adults 65 years and older^2^. Similar age-dependent patterns are observed for cancer and chronic diseases such as chronic obstructive pulmonary disease (COPD) and type 2 diabetes, whose incidence markedly increases per decade of life^3,4^. Despite being a central driver of disease susceptibility, the aging process itself remains poorly understood. As a result, identifying a precise and quantifiable biology of aging has emerged as a major scientific priority, with the promise of improving diagnosis, guiding preventive strategies, and enabling new therapeutic approaches for prevalent age-related conditions.

Aesthetic manifestations of aging such as facial aging patterns have long been known to vary between individuals^5^. Recent studies utilizing machine learning models known as aging clocks demonstrate that the internal aging process (i.e., biological aging) involves similar inter- and intra-individual variation. Estimators of organ-specific biological age including epigenetic^6^, proteomic^7–9^, transcriptomic^10^, multi-omic^11^, and MRI image-based^12^ aging clocks demonstrate that molecular shifts occur asynchronously in different organs over the lifespan, correlating with the incidence of clinical disease at population scale. In other words, organs and tissues can age at different rates within a single individual.

Utilizing the measurement of thousands of proteins collected from the blood, plasma proteomic aging clocks provide a non-invasive window into the aging state of particular organs. These models demonstrate that individuals with biologically older brains are associated with an elevated risk of cognitive decline, a finding which has been reproduced in multiple cohorts and confirmed with orthogonal measures of brain health^13–16^. Beyond risk, related results point to youthful organ aging as a linkage to longevity and disease resilience that can be shaped by genetic, lifestyle, and environmental factors^16^. These discoveries, among others, suggest that rather than biological aging being a uniform, linear driver of disease susceptibility, it represents a heterogeneous and dynamic shaping influence, which is important to quantify, since it likely affects disease onset and trajectory.

Beyond organ aging, more recent advances in the field have led to the development of transcriptomic^10,17–19^ and epigenetic^20^ aging clocks that pinpoint the molecular hallmarks of aging at the resolution of the cell. These studies reveal that aging manifests differently across cell types, with biologically old and young cells uniquely contributing to cognitive decline in murine studies of neurodegeneration^19^, as well as immune^17^ and brain^18^ function in human populations. Intriguingly, biologically old cells that are associated with disease have been shown to be responsive to rejuvenating interventions such as exercise and partial reprogramming, suggesting cellular aging clocks encode actionable molecular targets^19^. While promising, existing studies of biological aging at the resolution of the cell are limited to transcriptomic or epigenetic modalities that rely on non-human animal models or surgically acquired tissues, which restricts scalability and raises questions about translational relevance to human populations.

In this study, we perform a comprehensive analysis of cellular aging using plasma proteomic measurements derived from more than 7,000 proteins and three independent cohorts encompassing over 60,000 individuals combined (**Fig. 1a**). We first identify proteins circulating in the blood likely to originate from cell types of neuronal, glial, immune, epithelial, myeloid, lymphoid, endothelial, among other putative sources. Leveraging these cell-to-protein mappings, we describe temporal patterns of cellular aging and pinpoint dynamic physiological events that occur during human lifespan. To explore biological age variation within human subjects, we then construct plasma proteomic aging clocks that measure the biological age of a broad spectrum of more than 40 different cell types. Finally, to determine the clinical relevance of cell-type biological age variation, we quantify age acceleration in the form of age gaps and evaluate associations with clinical and biological markers of disease and mortality both cross-sectionally and over extended follow-up.

**Figure 1.**
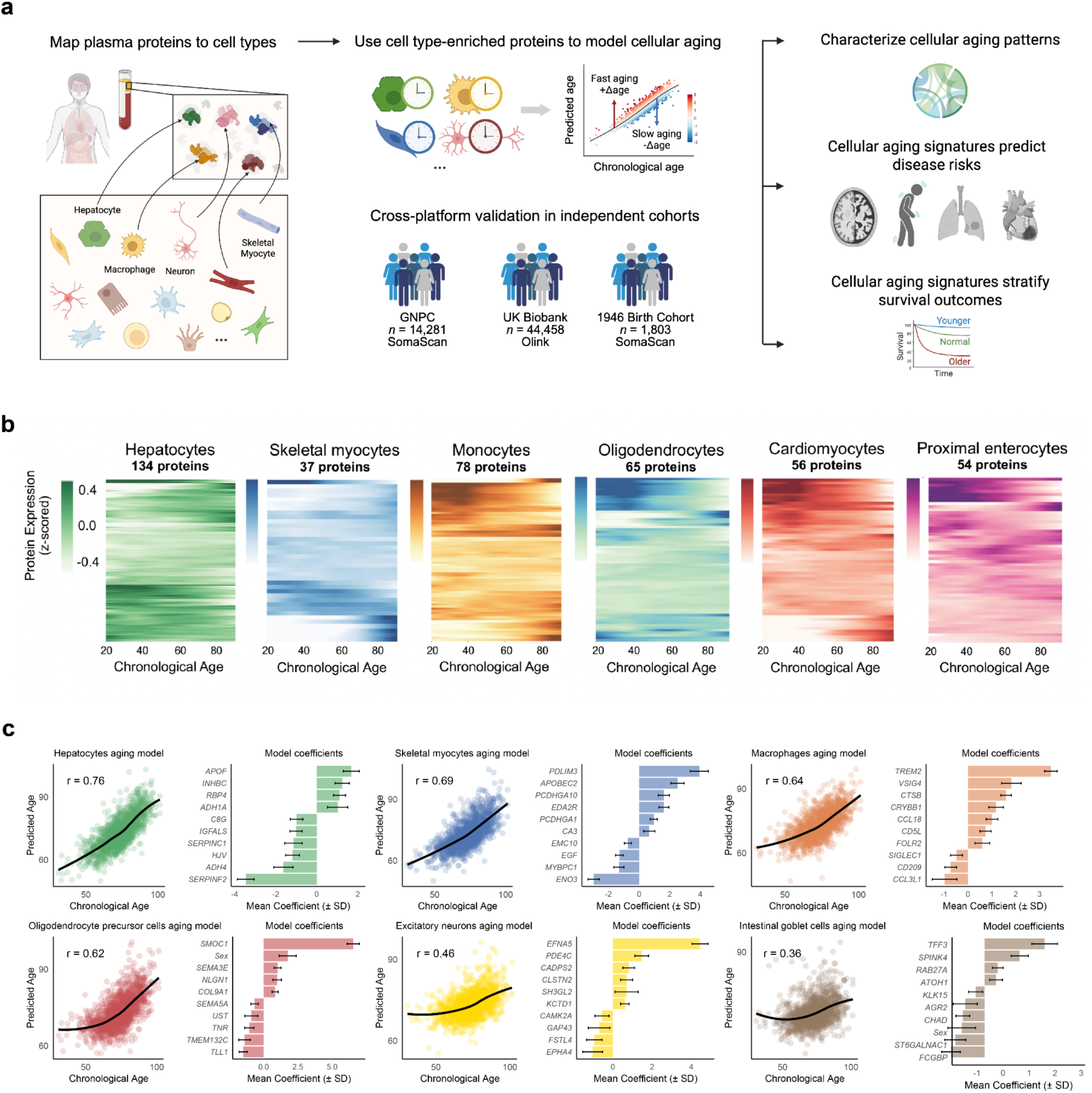
A framework for modeling cellular aging with plasma proteomics. **a,** Study design for evaluating plasma proteomic signatures of cellular aging across platforms and cohorts. Plasma proteins (7,289 on SomaScan, 2,923 on Olink) were mapped to cell types using the Human Protein Atlas (left panel). Cell type-enriched proteins were used to train machine learning models predicting chronological age, generating cell type-specific aging clocks. Cell type-specific age gaps—the disparity between predicted biological age and expected age for healthy individuals at the same chronological age—were calculated and z-scored to enable cross-cell-type comparisons. Cross-platform validation was performed across three independent cohorts: the Global Neurodegeneration Proteomics Consortium (GNPC, n=14,281, SomaScan), the UK Biobank (UKB, n=44,458, Olink), and the National Survey of Health and Development (NSHD, n=1,803, SomaScan). Cellular aging signatures were applied to characterize heterogeneous aging, predict disease risk, and stratify survival (right panel). **b,** Plasma protein expression trajectories for six illustrative cell types in healthy GNPC individuals (n=7,074). Trajectories were modeled using Locally Weighted Scatterplot Smoothing (LOWESS) regression spanning chronological ages 20 to 90 years. Each panel shows proteins assigned to the respective cell type. **c,** Illustrative cellular aging models trained on Knight-ADRC healthy individuals, the largest well-characterized healthy cohort in the GNPC. Scatter plots show estimated biological age versus chronological age with correlation coefficients (r). Bar plots display mean coefficients (± SD) of the top proteins by absolute magnitude in each cellular aging model.

### A framework for modeling cellular aging with plasma proteomics

Plasma protein expression is known to vary with age in an organ-specific manner, but whether similar patterns exist at the level of individual cell types remains unknown. To address this question, we constructed a blood-based framework that captures both chronological and biological cell-type aging patterns across the lifespan. Leveraging single-cell transcriptomic data in the Human Protein Atlas (**Methods, Extended Data Fig. 1**), we linked 60 human cell types to their corresponding plasma proteins. Guided by prior studies^16,21,22^, we classified genes as cell-type specific if they were expressed at least twofold higher in one cell type compared to any other cell type. Using this criterion, we found 16.5% of measurable plasma proteins in the SomaScan assay (1,202/7,289) and 24.2% of proteins in the Olink assay (708/2,923) mapped to specific cell types (**Supplementary Tables 1, 2**). Although the twofold enrichment threshold may seem permissive, many proteins mapped to cell types have fold changes markedly exceeding this criterion, supporting the premise that cell types possess strong individualized signatures that are detectable in plasma (**Methods, Extended Data Fig. 2**). Consistent with these signatures, we observed distinct age-related trajectories of plasma protein expression among cell types, illustrated for 7,074 healthy participants in the Global Neurodegeneration Proteomics Consortium (GNPC), a large-scale international multi-cohort neurodegenerative disease proteomics resource^23^(**Fig. 1b; Supplementary Table 3**).

Beyond individual cell-type patterns, we performed unsupervised hierarchical clustering of plasma protein trajectories to identify groups of proteins whose age-related changes occur in concert. Altogether, we identified 33 clusters ranging in size from 6 to 1,011 proteins that increased, decreased, or remained stable over time (**Extended Data Fig. 3**). Clusters exhibited significant and interpretable enrichment for cell types and biological pathways, spanning hepatocytes and regulation of hemostasis (Cluster 10, adjusted *p*-value=4.20×10^-5^, **Supplementary Table 4**), to B cells, erythroid cells, oligodendrocyte precursor cells, and nervous system development (Cluster 12, adjusted *p*-value=2.49 ×10^-10^, **Supplementary Table 4)**. Intriguingly, we found that cells of a shared lineage frequently co-occur within clusters (e.g. endoderm-derived ciliated cells, glandular and luminal cells or mesoderm-derived mesothelial cells and skeletal myocytes), a discovery that may reflect developmental shifts during lifespan and coordinated gene expression programs^24^. The stability of clusters was evaluated by performing a sensitivity analysis of trajectory clustering with bootstrap resampling (**Extended Data Fig. 4**), revealing our results to be robust. We speculate that co-clustered shifts between proteins from ontologically distant cell types could serve as a fruitful area of future study into mechanistic interactions of these tissues during aging.

These observations support our central hypothesis that cell types age at different rates, and their distinct aging trajectories can be captured through variations in plasma protein abundance. To test this, we built population-based models of biological aging for different cell types (i.e., “aging clocks”) by training machine learning models to predict chronological age based on the plasma abundance of cell type-specific proteins (see **Methods**). To demonstrate robustness and generalizability, we developed cellular aging models on two distinct proteomics platforms — SomaScan (measuring 7,289 proteins) and Olink (measuring 2,923 proteins) — and applied them to three independent cohorts: the Global Neurodegeneration Proteomics Consortium (GNPC, n=14,281, SomaScan), the National Survey of Health and Development (NSHD 1946 British Birth Cohort, n=1,803, SomaScan), and the UK Biobank (UKB, n=44,458, Olink).

For the SomaScan platform, we trained models using healthy individuals from the Knight Alzheimer’s Disease Research Center (KADRC), the largest well-characterized healthy cohort in the GNPC (**Fig. 1c** and **Extended Data Fig. 5**). These SomaScan-based models were then applied to the broader GNPC cohort for disease association analyses and to an independent cohort (NSHD) for external validation. We applied this approach to 60 cell types reflective of systemic aging, including hepatocytes, skeletal myocytes, cardiomyocytes, macrophages, T cells, B cells, NK cells, excitatory neurons, inhibitory neurons, oligodendrocytes, pancreatic endocrine cells, fibroblasts, among other distinct cell types (**Supplementary Table 1**)^25^. After applying performance quality control criteria (see **Methods)**, 43 cellular aging models were retained for downstream analyses (**Extended Data Fig. 5)**.

Using a similar approach, Olink-based cellular aging models were built using a training subset of the UK Biobank (n=21,983) and subsequently applied to a held-out UK Biobank test set (n=22,475). Given that the Olink platform captures fewer protein markers and thus fewer cell type-specific signatures, we additionally incorporated 14 “parental” lineage-level cell type models that aggregate ontologically related cell types (e.g., lymphoid lineage combining B cells, T cells, NK cells, and plasma cells; neurons combining excitatory neurons and inhibitory neurons) to capture robust aging signatures across platforms and to gain further insight into cellular aging at varying levels of granularity (**Supplementary Table 5, Extended Data Fig. 6**, **Methods**). After quality control, 48 cellular aging models were retained for Olink-based analyses.

We calculated age gaps for each cell type as the residual between an individual’s predicted cell-type biological age and the model-predicted biological age of an average healthy individual at the same chronological age. These age gaps offer a quantitative measure of relative biological age – or physiological state – allowing us to identify individuals whose cellular age deviates from chronological age-matched peers. Individuals with a positive age gap represent accelerated agers, whereas a negative age gap represents youthful agers. Age gaps were *z*-scored per aging model to facilitate comparison across cell types in downstream analyses. Extreme agers were identified as individuals with age gaps at least two standard deviations from the mean for that cell type^21^. This plasma proteomic framework enables population-scale measurement of cellular aging across the lifespan and investigation of how it influences disease susceptibility and resilience.

### Cellular aging signatures reveal individual heterogeneity

Cellular aging patterns exhibited substantial heterogeneity across individuals. To characterize age-dependent patterns of this heterogeneity, we applied cellular aging models to 7,074 healthy individuals in the GNPC cohort and visualized the distribution of extreme agers of different cell types across five chronological age windows (Fig. 2a). Notably, neuronal and glial cell types—including Schwann cells, inhibitory and excitatory neurons—showed elevated age gaps predominantly in later life, with 7.1%, 6.6%, and 6.4% of individuals classified as extreme agers in the over 85-year-old age group, respectively. In contrast, intestinal goblet cells and ciliated cells exhibited early onset of accelerated aging, affecting 4.9% and 4.2% of individuals under 60 years old. We hypothesize that the prevalence of cell-specific biological extreme agers in unique chronological age windows is reflective of cellular vulnerabilities and the timing of disease onset. While accelerated aging of neurons and Schwann cells with advanced age may be related to cognitive impairment and loss of sensory perception with age, accelerated aging of goblet and ciliated cells in younger individuals may point to increased gut leakiness and inflammation and reduced cilia function and ependymal cilia barrier integrity during midlife^26–29^. These observations reveal differential biological aging trajectories among various cell populations, suggesting cell type-specific windows of vulnerability to cellular aging across lifespan.

**Figure 2:**
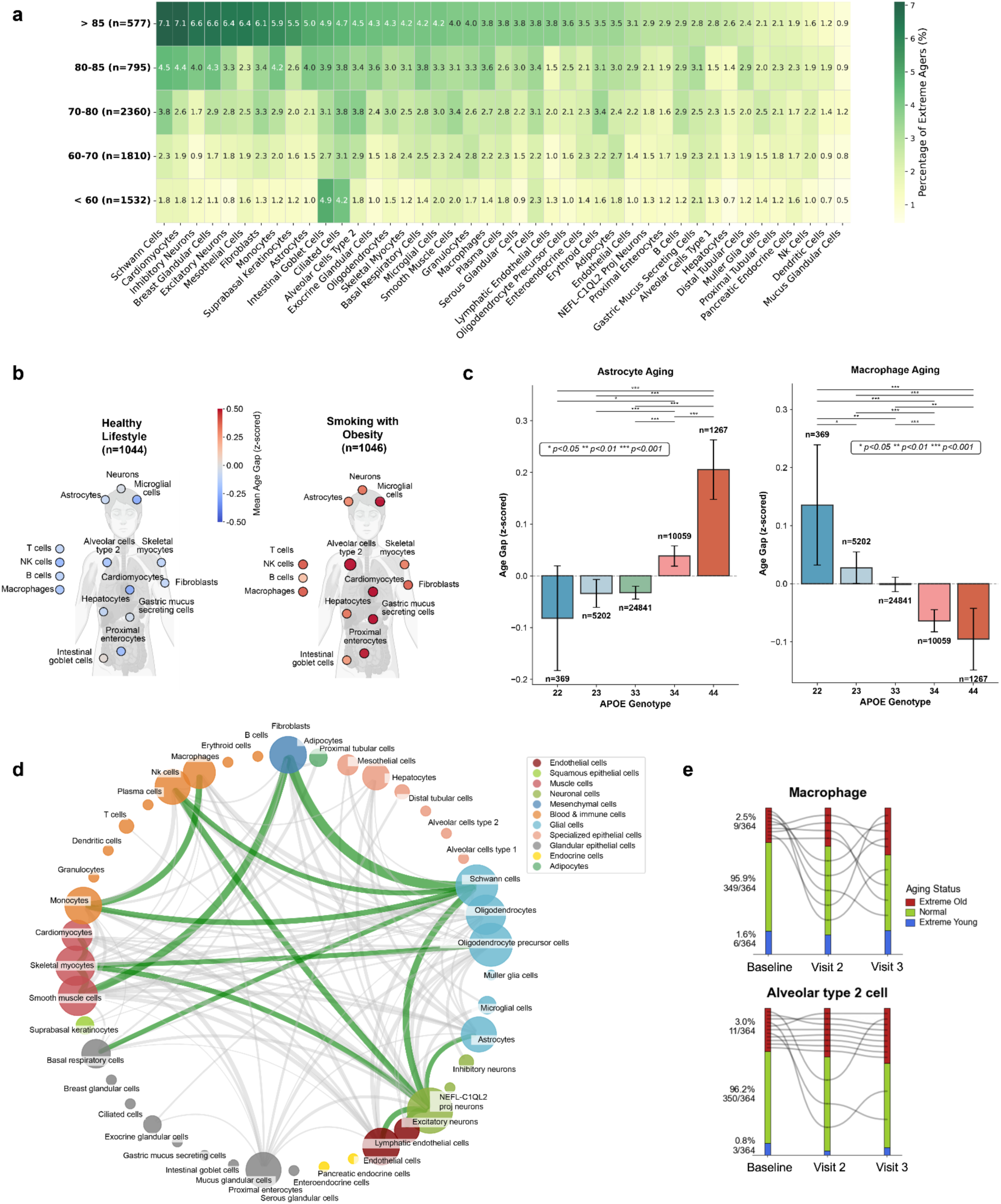
Cellular aging signatures reveal individual heterogeneity. **a,** Age-dependent patterns of extreme cellular aging in healthy individuals of the GNPC cohort (n=7,074). Heatmap shows the percentage of extreme agers across 43 cell types (columns) and five age windows (rows), with extreme agers defined as individuals with age gaps at least two standard deviations from the mean. **b,** Cellular aging and modifiable risk factors in the UKB cohort. Smoking and obesity are associated with accelerated aging across multiple cell types (right, n=1,046), while “healthy lifestyle”—defined as never smoking, no regular alcohol consumption, at least 5 days per week of 10+ minutes of moderate or vigorous physical activity, BMI <25, and waist circumference <90cm for men or <84cm for women—is associated with younger biological ages (left, n=1,044). Color intensity corresponds to the mean z-scored age gap (red: older, blue: younger). **c,** APOE genotype shows antagonistic patterns on cellular aging. Bar plots show mean z-scored age gaps for astrocytes (left) and macrophages (right) stratified by APOE genotype (22: n=369, 23: n=5,202, 33: n=24,841, 34: n=10,059, 44: n=1,267) in the UKB cohort. Error bars represent standard error of the mean. APOE2 carriers exhibited younger astrocyte profiles but older macrophages, while APOE4 carriers showed the inverse pattern. Statistical comparisons between genotype groups are shown above bars (*p<0.05, **p<0.01, ***p<0.001). **d,** Correlation network showing cellular aging patterns across cell types in GNPC healthy individuals (n=7,074). Each node represents a specific cell type, with two nodes connected if the correlation in age gaps between the two cell types is above a threshold of 0.35. Edge width corresponds to correlation strength, with green edges highlighting the top 15 correlations. **e,** Stability of extreme cellular aging over a 10-year period for macrophage and AT2 agers in the NSHD cohort (Baseline: 63.2±1.1 years; Visit 2: 70.7±0.7 years; Visit 3: 72.9±0.6 years; n=364 across all visits). Bars show the proportion of extreme-old (red), extreme-young (blue), and normal (green) agers at each visit, with extreme groups proportionally scaled for visualization. Black lines show trajectories of baseline extreme-old agers.

Across all healthy individuals in the GNPC cohort, we found 35.4% had no extreme cellular age gaps and 24.4% had accelerated aging in a single cell type, while 1.5% of the population experienced widespread acceleration across 10 or more cell types. For each cell type studied, we found 0.9–3.8% of the population had extremely old cells, and 0.7–3.2% had extremely young cells (Extended Data Fig. 7a). Similarly, in the UK Biobank cohort, 26.1% showed no extreme cellular age gaps, 22.7% had accelerated aging in a single cell type, and 2.8% exhibited widespread acceleration across 10 or more cell types, with 1.9–4.1% showing extreme acceleration and 0.7–2.5% showing extremely young cells per cell type (Extended Data Fig. 7b).

Cell type-specific age gaps also demonstrated associations with modifiable risk factors. Smoking and obesity were linked to a widespread increase in biological age across multiple cell types, while individuals with healthier lifestyle profiles, including never smoking, no alcohol consumption, BMI lower than 25 without enlarged waist circumference, sufficient sleep (≥7 hours nightly), and regular exercise (≥5 days weekly), showed overall younger cellular ages (Fig. 2b).

Given that extreme aged cells can co-occur within an individual, we sought to characterize these interactions in aggregate. Our examination of age gap profiles across >7,000 individuals in GNPC suggests that cellular aging occurs in a concerted fashion across a small number of cell types (Fig. 2d). These coordinated patterns were particularly pronounced among excitatory neurons, myelinating cells, and endothelial cells, suggesting shared or synchronized pathways. Certain cell populations—such as excitatory neurons, Schwann cells, NK cells, macrophages, skeletal myocytes, and fibroblasts—emerged as potential “aging hubs,” showing correlations with multiple other cell types. In contrast, epithelial cell types tended to exhibit more isolated or weakly correlated age gap profiles.

If extreme aging truly represents a biologically stable phenotype, cells classified as extremely old at baseline should remain old over time. We investigated this hypothesis in the NSHD, the world’s oldest continuously followed population-based birth cohort^30^, wherein we examined the stability of extreme cellular aging over a 10-year period in 364 individuals with near identical chronological ages. In Fig. 2e, we show two representative profiles for baseline extreme old cell type agers: alveolar type 2 cells (AT2, 11/364 individuals) and macrophages (9/364 individuals). Among these groups, there was substantial retention of extreme aged states: 81% of baseline AT2 extreme agers remained old throughout the 10-year follow-up period, while 55% of baseline macrophage extreme agers retained extreme aging status. Evaluating each timepoint separately, we observed a 1.36-fold (AT2) and 1.56-fold (macrophage) expansion of these extreme ager cell type populations over time. We extended this analysis to baseline extreme young cell-type agers in Extended Data Fig. 8. Our observations suggest that individuals in a state of extreme cellular aging tend to retain this state over time. Certain cell types show a tendency towards old or young states, potentially reflecting differential resilience or susceptibility of specific cell types to biological aging.

Analysis of genetic influences revealed specific genotypes could accelerate aging in certain cell types while preserving others, contributing to heterogeneous cellular aging patterns (Fig. 2c). APOE genotype, a major risk factor for neurodegenerative disease, showed dose-dependent, antagonistic effects on immune versus central nervous system cell aging: APOE2 carriers exhibited significantly younger astrocyte profiles but older macrophages, while APOE4 carriers showed the inverse, with older astrocytes but younger macrophages. This inverse pattern suggests antagonistic pleiotropy operating at cellular resolution, with the same allele conferring benefits in certain cell types while imposing costs in others. Such trade-offs align with the evolutionary hypothesis that APOE4’s enhanced immune vigilance conferred survival advantages under historically high pathogen burdens, despite accelerated brain aging that increases Alzheimer’s disease risk in modern extended lifespans^31–34^.

Taken together, these analyses illustrate that plasma proteome-derived, cell-type specific biological age estimates reveal heterogeneous cellular aging profiles, with potential implications for disease susceptibility and overall health trajectories.

### Cellular aging signatures record neurodegenerative diseases

Meaningful biological age estimates should capture underlying physiological states and therefore associate with age-related health trajectories and disease outcomes. To this end, we sought to assess whether cellular age gaps correlate with disease status, focusing on the neurodegenerative and neurological conditions present in the GNPC: Alzheimer’s disease (AD, n=2,761), amyotrophic lateral sclerosis (ALS, n=245), Parkinson’s disease (PD, n=476), frontotemporal dementia (FTD, n=199), and mild cognitive impairment–subjective cognitive impairment (MCI-SCI, n=1,992). The chronological age distributions of patients with these five neurodegenerative conditions in the GNPC aligned with epidemiological studies (Extended Data Fig. 9), with ALS affecting individuals at notably younger ages (mean=58.1 years, s.d.=10.9), followed by FTD (mean=64.0 years, s.d.=10.1), while AD (mean=75.8 years, s.d.=8.8), PD (mean=74.3 years, s.d.=8.8), and MCI-SCI (mean=71.1 years, s.d.=9.1) predominantly manifested at older ages.

We tested the associations between all 43 cellular age gaps and each neurodegenerative disease using Point-Biserial correlation with Benjamini-Hochberg false discovery rate correction (Fig. 3a). The strongest association among all disease-cell type pairs was between ALS and skeletal myocyte aging (r=0.43, adjusted *p*-value=1.36×10^-15^), consistent with known pathophysiological motor neuron degeneration and muscle atrophy in ALS^35–37^.

**Figure 3.**
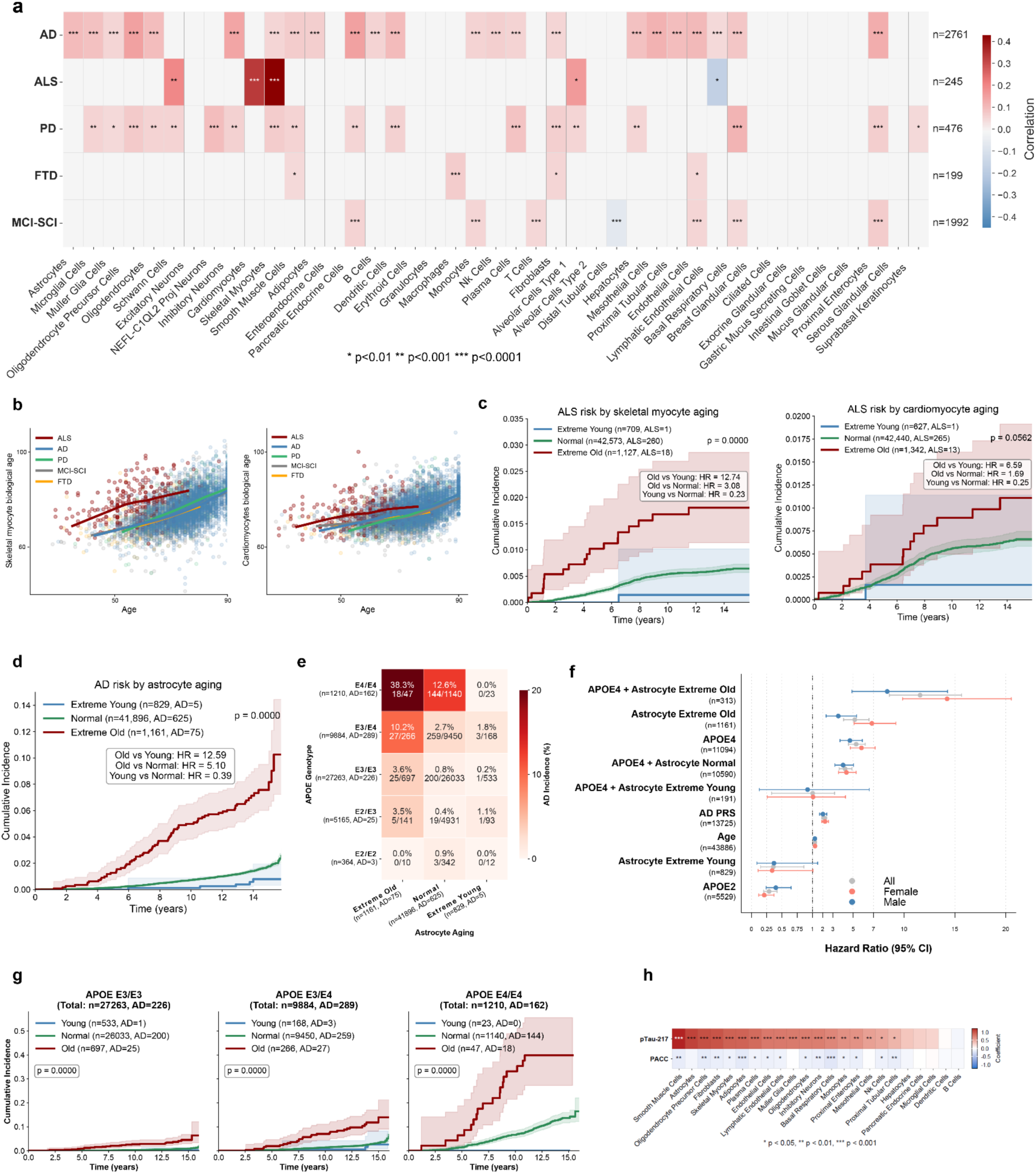
Cell-type specific biological age estimates record neurodegenerative diseases **a,** Correlation between cell type-specific age gaps and neurodegenerative disease status in GNPC. Analysis includes Alzheimer‘s disease (AD, n=2,761), amyotrophic lateral sclerosis (ALS, n=245), Parkinson’s disease (PD, n=476), frontotemporal dementia (FTD, n=199), and mild cognitive impairment/subjective cognitive impairment (MCI-SCI, n=1,992). Associations with adjusted p-value below 0.01 and correlation magnitudes greater than 0.05 are shown in color, with color intensity corresponding to correlation strength and significance levels annotated. P-values adjusted using Benjamini-Hochberg (BH) procedure (*p<0.01, **p<0.001, ***p<0.0001). **b,** Estimated biological age of skeletal myocytes (left) and cardiomyocytes (right) versus chronological age by disease groups in GNPC. Each dot represents an individual, colored by disease diagnosis. LOWESS regression fit estimates the relationship between estimated biological and chronological age for each group. **c,** ALS cumulative incidence by skeletal myocyte (left) and cardiomyocyte (right) aging status over 15 years in UKB: Extreme agers (red), normal agers (green), youthful agers (blue), with sample sizes in legend. Shaded regions represent 95% confidence intervals (CI). Risk stratification assessed by log-rank test, with p-value indicated. **d,** AD cumulative incidence by astrocyte aging status over 15 years in UKB: Extreme agers (red, n=1,161), normal agers (blue, n=41,896), youthful agers (green, n=829). Log-rank tests demonstrated significant risk stratification across aging groups (p<0.0001). **e,** AD cumulative incidence by APOE genotype and astrocyte aging status over 15 years in UKB. Rows: APOE genotype (E2/E2 [n=364], E2/E3 [n=5,165], E3/E3 [n=27,263], E3/E4 [n=9,884], E4/E4 [n=1,210]). Columns: astrocyte aging (extreme-old [n=1,161], normal [n=41,896], youthful [n=829]). Color intensity indicates cumulative incidence percentage. Numbers displayed in each cell indicate the total number of individuals and the number who developed AD during follow-up. **f,** Hazard ratios with 95% CI for incident AD by risk factor in UKB. Overall (gray) and sex-stratified (females, pink; males, blue). **g,** AD cumulative incidence by astrocyte aging status across APOE genotypes in UKB. Extreme (red), normal (green), youthful (blue) agers. Left: APOE3/3 (n=27,263). Middle: APOE3/4 (n=9,884). Right: APOE4/4 (n=1,210). Shaded regions represent 95% CI. Risk stratification by astrocyte aging status was significant for each genotype (log-rank test, p<0.0001). **h,** Cellular age gap associations with plasma pTau-217 burden (pg/mL) and Preclinical Alzheimer Cognitive Composite (PACC) in the NSHD Insight-46 substudy (n=483). Lower PACC corresponds to worse cognition. Multiple comparisons correction was performed using the BH procedure (*p<0.05, **p<0.01, ***p<0.001).

Intriguingly, along with skeletal myocytes, we observed that cardiomyocytes showed accelerated aging in ALS patients (r=0.33, adjusted *p*-value=4.08 x 10^-9^), consistent with emerging evidence of cardiac abnormalities in ALS patients, including rare sudden cardiac arrest in advanced stages of ALS^38–41^ and potentially driven by shared molecular pathways affecting both cardiac and skeletal muscle tissues. Indeed, one study of 35 patients with ALS without a history of cardiac disease reported that myocardial mass and lower left and right ventricular volumes were significantly reduced in ALS^41^. Given the strong association between ALS and both skeletal myocyte and cardiomyocyte aging, we visualized the estimated cellular age for these cell types versus chronological age across disease groups (Fig. 3b). Locally Weighted Scatterplot Smoothing (LOWESS) regression analysis revealed that ALS patients exhibited pronounced acceleration of skeletal myocyte aging across chronological age, in alignment with the central pathophysiology of ALS involving progressive degeneration of motor neurons and consequent muscle atrophy^35–37^. In addition to both skeletal- and cardiomyocytes, Schwann cell aging was significantly linked to ALS, consistent with a growing body of research linking the function of these cells to motor neuron plasticity and survival and to modulation of inflammatory responses^42^.

AD was associated with accelerated aging across a wide range of cell types, most prominently oligodendrocyte precursor cells (r=0.15, adjusted *p*-value=1.86×10^-44^), pancreatic endocrine cells (r=0.15, adjusted *p*-value=3.49×10^-44^), inhibitory neurons (r=0.15, adjusted *p*-value=1.48×10^-40^), proximal enterocytes (r=0.13, adjusted *p*-value=3.85×10^-33^), and astrocytes (r=0.10, adjusted *p*-value=3.25×10^-21^). Accelerated aging of oligodendrocyte precursors in AD aligns with discoveries from unbiased transcriptomic studies linking these cells with brain aging and AD^43,44^. Intriguingly, while aging of inhibitory neurons was linked to AD, aging of excitatory neurons was not (Fig. 3a), potentially supporting a growing body of literature implicating network hyperexcitability related to loss of synaptic inhibition and selective vulnerability of inhibitory interneurons in the pathophysiological process^45,46^. Notably, while we recognized that AD comorbidities such as diabetes may introduce confounding effects, stratified analysis revealed that AD patients exhibited accelerated pancreatic endocrine cell aging also in the absence of type 2 diabetes (T2D), while T2D amplified this effect in a compounding manner (Extended Data Fig. 10). These findings highlight the systemic nature of AD pathophysiology, with particularly strong connections to gut epithelial aging, metabolic dysregulation, and oligodendroglial-neuronal interactions^46–54^.

To further prioritize cell types linked with neurodegenerative diseases, we investigated odds ratios between cell type-specific extreme ager status and disease states (Extended Data Fig. 11). ALS exhibited an exceptionally strong odds ratio with accelerated skeletal myocyte aging (OR=7.85, adjusted p-value=5.50×10^-5^), indicating that individuals with extreme skeletal myocyte aging were > 7 times more likely to have ALS than those without acceleration. In the ALS cohort (n=355; 245 ALS patients, 110 controls), 53 of 57 (93.0%) skeletal myocyte extreme agers had an ALS diagnosis. This pronounced association is consistent with reported muscle pathology in ALS and aligns with emerging evidence that peripheral tissues may contribute to disease progression alongside motor neuron degeneration^35,36,55–59^.

We discovered that a range of systemic aging signatures were associated with cognitive and motor neurodegeneration. Specifically, accelerated aging of basal respiratory cells (OR=2.52, adjusted *p*-value=3.38×10^-4^) and plasma cells were linked to Parkinson’s disease (OR=2.15, adjusted *p*-value=0.02), potentially pointing towards alterations to immune system pathways^60^, or vulnerabilities consistent with increased prevalence of respiratory comorbidity in this patient population^61–63^. FTD showed strong cross-sectional association with the cell type initially labeled as “horizontal cell”, which, to accurately reflect the biological underpinning of this aging signature, we refer to as NEFL-C1QL2 projection neuron aging (Extended Data Fig. 12). NEFL is a widely recognized biomarker of axonal injury, often markedly raised in FTD^64^, while C1QL2 is a synaptic organizer known to be prominent in temporo-limbic structures^65^ vulnerable to fronto-temporal lobar degeneration. Interestingly, more modest cellular aging associations were evident for patients with MCI-SCI, a heterogeneous group characterized by subjective (SCI) or mild (MCI) cognitive symptoms. Predominant homeostatic cellular aging biology is consistent with these more subtle cognitive phenotypes observed, underscoring the correspondence we consistently observed between systemic biology and neurological status. Nevertheless, it is intriguing that accelerated proximal enterocyte and exocrine glandular cell aging emerged as linked to MCI-SCI, potentially reflecting gut-brain axis dysfunction as an early peripheral prodromal signal.

Collectively, these disease-patterned cellular aging signatures not only support several plausible pathophysiological associations, but also may highlight potential novel peripheral cellular targets for mechanistic study and therapeutic intervention.

While the cross-sectional study described above reveals associations between cellular aging and disease status, a critical question is whether these aging signatures can predict future disease onset. We therefore examined the value of cellular aging signatures for stratifying risk of incident neurodegenerative disease using the UKB cohort. For ALS, individuals with extreme skeletal myocyte aging demonstrated markedly elevated risk compared to those with youthful cellular aging (hazard ratio = 12.74), and Kaplan-Meier curves revealed clear risk stratification by aging status (Fig. 3c, log-rank *p*<0.001). Interestingly, the relationship between the plasma signature of accelerated skeletal muscle aging and future ALS diagnosis persists even when only considering cases diagnosed more than three years after blood draw and cellular aging assessment (HR=1.93, 95% CI: 1.02-3.64 for extreme agers compared to the rest of population, Extended Data Fig. 13). Time to diagnosis for ALS is often delayed into the range of 8-15 months, but the multi-year gap seen here suggests pathological mechanisms affecting muscle tissue begin years before symptom onset^66^. Similarly, extreme cardiomyocyte aging showed a hazard ratio of 6.59 compared to youthful agers. While this association raises the interesting question of whether myocardial dysfunction may also presage the diagnosis of ALS, caution is warranted as some cardiomyocyte-assigned proteins including FABP3 are also expressed to a limited degree in skeletal myocytes.

When examining the prognostic value of cellular aging signatures for incident AD risk, astrocyte aging emerged as the strongest predictor (Fig. 3d). Individuals with extreme astrocyte aging demonstrated a 12.59-fold increased risk compared to those with youthful astrocyte aging, while the Kaplan-Meier curves revealed significant risk stratification by aging status (log-rank *p*<0.001). The risk stratification remained robust across independent APOE genotype subgroups (Fig. 3g). Consistent with the known genetic risk gradient for AD, disease incidence increased progressively from APOE3/3 carriers (n=27,263) to APOE3/4 carriers (n=9,884) to APOE4/4 carriers (n=1,210), reflecting the escalating impact of APOE4 allele dosage on AD risk. Notably, within each genotype group, extreme astrocyte aging consistently identified individuals at elevated AD risk compared to those with normal and youthful astrocyte aging, demonstrating that astrocyte aging provides independent risk stratification beyond APOE genotype.

Given our previous finding that astrocytes showed accelerated aging in APOE4 carriers in a dose-dependent manner (Fig. 2c) and that APOE4 is the major genetic risk factor for AD, we examined how APOE genotype and astrocyte aging jointly influence AD incidence. Analysis of cumulative incidence across all combinations of APOE genotype and astrocyte aging status revealed pronounced synergistic effects (Fig. 3e**, Extended Data** Fig. 14). Individuals who were homozygous for APOE4 and had extreme astrocyte aging showed the highest cumulative incidence of 38.3% over 15 years of follow-up, compared to 12.6% for homozygotes with normal astrocyte aging. This risk gradient was consistent across genotypes: APOE3/4 carriers with extreme astrocyte aging showed 10.2% cumulative incidence versus 2.7% with normal astrocyte aging, and APOE3/3 carriers demonstrated a similar pattern (3.6% versus 0.8%, respectively). Most importantly, and of potential therapeutic relevance, none of the 23 APOE4/4 carriers with youthful astrocytes and 1.8% of APOE3/4 carriers developed AD. While only 10 APOE2/2 carriers exhibited extreme astrocyte aging, APOE2/2 carriers maintained low incidence overall, consistent with the known protective effects of the APOE2 allele. Neuroinflammatory factors are considered pivotal to AD pathological propagation^67^. These results potentially implicate central “inflammaging” mechanisms^68^ and astrocyte activation as key to understanding why only some APOE4 carriers fall susceptible to AD, while others remain protected.

To contextualize the prognostic power of astrocyte aging relative to established AD risk factors, we performed a comparative analysis against AD polygenic risk score (PRS), APOE4 carrier status, and chronological age (Fig. 3f). Excess AD risk associated with extreme astrocyte aging (HR=5.16, 95% CI: 4.06-6.56) was comparable to APOE 4 carrier status (HR=5.30, 95% CI: 4.54-6.18) and exceeded both PRS (HR=2.14, 95% CI: 1.92-2.39) and older chronological age (HR=1.24, 95% CI: 1.22-1.27). Individuals who both carried APOE4 and had extreme astrocyte aging were at highest risk (HR=11.58, 95% CI: 8.56-15.66). Notably, sex-stratified analysis of AD risk revealed women were more vulnerable to the harmful associations linked to not only APOE4 (HR=5.82, 95% CI: 4.73-7.16 for females compared to HR=4.68, 95% CI: 3.71-5.90 for males), but also astrocyte extreme aging (HR=6.84, 95% CI: 5.09-9.20 for females compared to HR=3.54, 95% CI: 2.35-5.34 for males). Moreover, possessing both APOE4 and astrocyte extreme aging conferred a markedly greater increase in AD risk for women (HR=14.23, 95% CI: 9.86-20.54) than men, further contextualizing sex-specific patterns of AD pathogenesis. Most excitingly, youthful astrocytes reduced AD risk by over 60%.

Together, these findings highlight astrocyte aging as a potentially powerful biomarker that stratifies AD risk independently of and synergistically with APOE genotype. Maintaining youthful astrocyte function may be a potential therapeutic strategy to mitigate disease burden, especially in genetically predisposed individuals.

Furthermore, given the significant associations between cell type-specific aging and neurodegeneration, we examined the relationship between cell type age gaps and Clinical Dementia Rating (CDR) scores, a composite measure of dementia severity and general cognitive and functional performance^69^. As the GNPC aggregates data from multiple independent neurodegenerative cohorts with varying clinical assessments, we focused our analysis on the eight cohorts that collected CDR data to examine relationships between cellular aging and cognitive function. Among all cell types, the aging of oligodendrocyte precursor cells and inhibitory neurons demonstrated the strongest correlation with CDR (Extended Data Fig. 15). When visualizing age gaps across cohorts and stratifying by CDR score, oligodendrocyte precursor cell aging showed a consistent stepwise increase with worsening cognitive impairment, with particularly pronounced effects in cohorts J, F, and N. Similarly, inhibitory neuron age gaps also increased with higher CDR scores across cohorts. These observations further illustrate the biological relevance of cell type-specific aging in cognitive decline.

To deepen our analysis of cellular aging and AD, we further examined age gap associations in participants from the NSHD Insight-46 neuroimaging substudy (n=483)^70^. Leveraging neurological deep-phenotyping of Insight-46 participants, we performed cross-sectional regression analyses evaluating the association between plasma pTau-217 burden (pg/mL) and Pre-Alzheimer’s Cognitive Composite (PACC) score with cellular age gaps (Fig. 3h). Given plasma pTau-217 reliably identifies brain amyloid-β and tau-pathology with accuracy comparable to cerebrospinal fluid and PET biomarkers, even in preclinical stages, it serves as a robust orthogonal measure for validating associations between cell-type specific aging and AD pathology^71^. Although CDR scores were not collected as part of the study protocol in Insight-46, the cohort reports the Pre-Alzheimer’s Cognitive Composite (PACC), a validated measure sensitive to early cognitive decline that is widely used in preclinical dementia and Alzheimer’s disease research^72^.

In agreement with results obtained in the GNPC cohort, we found significant associations between plasma pTau-217 burden and several neuronal and glial cell-type age gaps, including astrocytes (ꞵ=1.08, SE=0.19, adjusted *p*-value=3.92×10^-7^), oligodendrocyte precursor cells (ꞵ=1.02, SE=0.19, adjusted *p*-value=1.19×10^-6^), Müller glia cells (ꞵ=0.77, SE=0.19, adjusted *p*-value=1.99×10^-4^), oligodendrocytes (ꞵ=0.76, SE=0.19, adjusted *p*-value=2.2×10^-4^), and inhibitory neurons (ꞵ=0.76, SE=0.19, adjusted *p*-value=2.2×10^-4^). Concordant with GNPC, we further identified significant associations between oligodendrocyte precursor cell (ꞵ=-0.16, SE=0.04, adjusted *p*-value=2.2×10^-3^) and inhibitory neuron (ꞵ=-0.13, SE=0.04, adjusted *p*-value=9.3×10^-3^) age gaps with poorer cognitive performance as measured by PACC. Additional prominent associations with PACC were observed for adipocytes (ꞵ= −0.18, SE =0.04, adjusted *p*-value=7.3×10^-4^) and basal respiratory cells (ꞵ=-0.20, SE=0.04, adjusted *p*-value=1.72×10^-4^), likely reflective of systemic aging having downstream effects on cognitive health (Fig. 3h)^73^.

Collectively, these findings demonstrate that cellular aging signatures derived from plasma proteomics are robustly associated with neurodegenerative diseases across multiple cohorts and offer granular insights into the underlying pathophysiology at the cellular level.

### Cellular aging signatures show prognostic value for chronic disease and cancer risk

We next investigated whether cellular aging signatures have prognostic value for cancer, stroke, and additional incident chronic disease by leveraging the longitudinal depth and comprehensive phenotyping of the UKB cohort. We performed Cox proportional hazards analyses to assess disease incidence over 15 years of follow-up. We found that cellular aging signatures from both tissue-archetypal cells and interacting stromal and immune cells strongly prognosticated future disease. For incident COPD, extreme aging in AT2 cells and respiratory epithelia showed the most prognostic power (HR=6.31, 95% CI: 5.57-7.13 and HR=5.45, 95% CI: 4.75-6.25, respectively) (Fig. 4a and 4c). For incident heart failure, extreme aging in muscle cells and fibroblasts was most prognostic (HR=4.65, 95% CI: 4.00-5.41 and HR=4.62, 95% CI: 3.98-5.39, respectively), consistent with known mechanisms of dysfunctional remodeling following myocardial damage or with aging^74,75^. For incident stroke, extreme aging in NEFL-C1QL2 projection neurons demonstrated the strongest prognostic value (HR=3.03, 95% CI: 2.47-3.72), followed by microglia (HR=2.84, 95% CI: 2.33-3.47) (Extended Data Fig. 16). The prominence of NEFL-C1QL2 projection neurons suggests particular vulnerability of neurite structures (where NEFL and C1QL2 are located) to preclinical vascular changes presaging occurrence of clinically significant cerebrovascular events, possibly reflecting the prognostic value of white matter changes for future stroke^64,76^.

**Figure 4.**
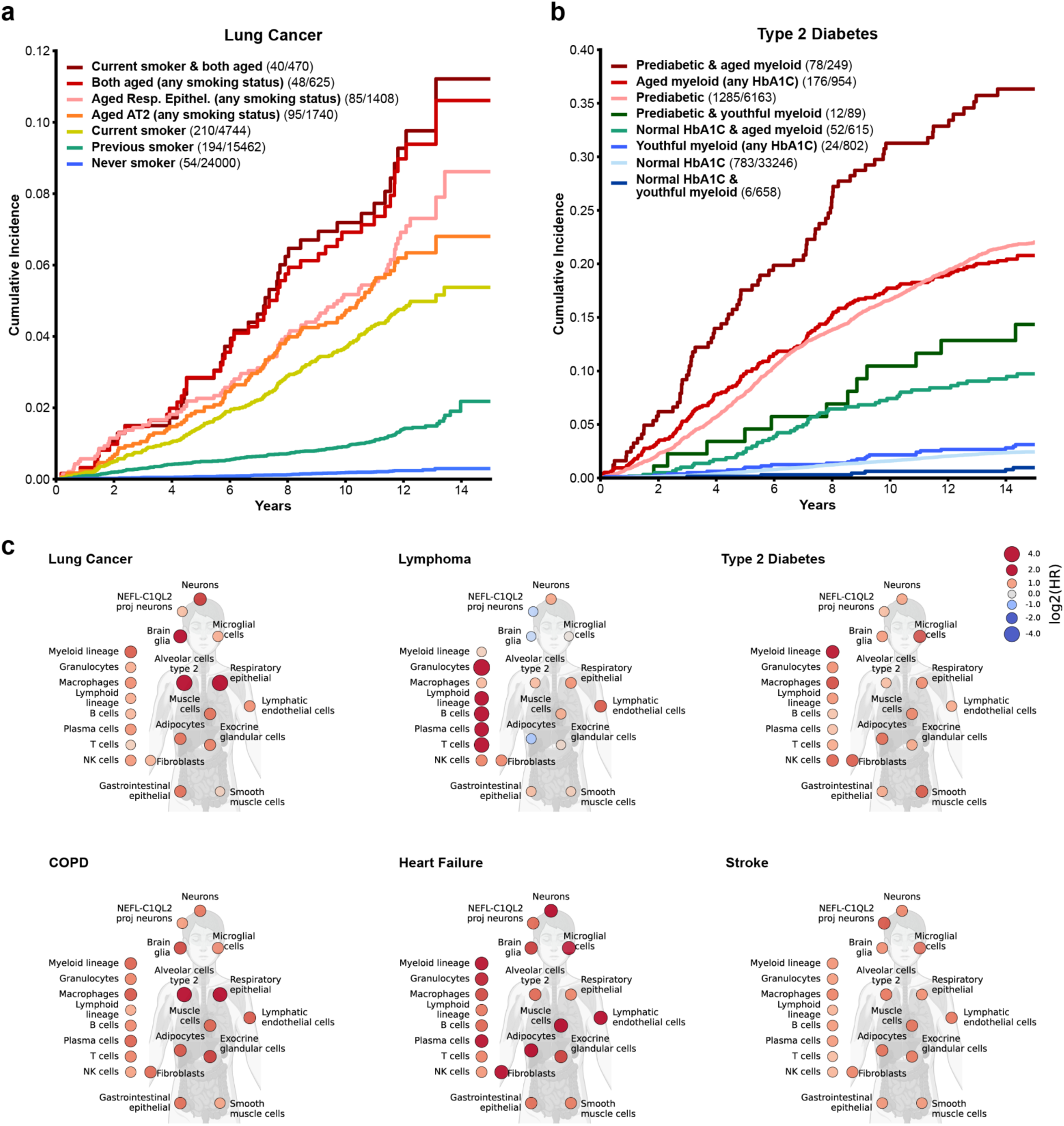
Cellular aging signatures predict cancer and chronic disease risk. **a,** Lung cancer cumulative incidence by smoking status and respiratory cell aging over 15 years in the UKB cohort. Groups (highest to lowest incidence): current smokers with extreme aging in both alveolar type 2 and respiratory epithelial cells (dark red), individuals with both extreme aging (red), respiratory epithelial cell extreme aging (pink), alveolar type 2 cell extreme aging (orange), current smokers (yellow), previous smokers (green), and never smokers (blue). Sample sizes and case counts are indicated in legend. **b,** Type 2 diabetes cumulative incidence by HbA1c status and myeloid lineage aging over 15 years in UKB. Groups (highest to lowest incidence): pre-diabetic individuals with extreme myeloid lineage aging (dark red), individuals with extreme myeloid aging (red), pre-diabetic individuals (pink), pre-diabetic individuals with youthful myeloid aging (dark green), individuals with normal HbA1c and extreme myeloid aging (green), individuals with youthful myeloid aging (blue), individuals with normal HbA1c (light blue), and individuals with normal HbA1c and youthful myeloid aging (navy). Shaded regions represent 95% CI. Sample sizes and case counts in legend. **c,** Body plots show hazard ratios for incident disease by extreme cellular aging, adjusted for age and sex. Diseases: lung cancer, lymphoma, type 2 diabetes, COPD, heart failure, and stroke. Displayed cell types represent the union of the top five prognostic cell types for each disease. Color intensity and size indicate hazard ratio (log scale).

Interestingly, for incident lymphoma, extreme aging in B cells was third most prognostic (HR=6.63, 95% CI: 4.25-10.33), following granulocytes (HR=10.09, 95% CI: 6.86-14.83), and T cells (HR=6.75, 95% CI: 4.33-10.54). In B-cell lymphoma, dysregulation of both T-lymphoid and myeloid precursor-derived cells is known to contribute to malignancy pathogenesis^77,78^. However, this finding raises the question of whether coordinated aging across the hematopoietic niche may facilitate early lymphomagenesis. Notably, while difficult to compare directly, these hazard ratios are higher than those for lymphoid clonal hematopoiesis of indeterminate potential (L-CHIP) and rival those of lymphoid mosaic chromosomal alterations (L-mCA)^79^, and their interaction may represent a promising area of future study.

For lung cancer, extreme aging in AT2 cells and respiratory epithelia was most prognostic (HR=8.39, 95% CI: 6.68-10.52 and HR=8.47, 95% CI: 6.69-10.71, respectively) (Fig. 4c). Notably, these cellular aging signatures enhanced lung cancer risk stratification beyond the known risk factor of smoking status (Fig. 4a). Over 15 years of follow-up, cumulative lung cancer incidence showed distinct trajectories based on cellular aging profiles and smoking history. Current smokers with extreme aging in both AT2 and respiratory epithelial cells exhibited the highest risk (HR=15.33, 95% CI: 11.02-21.31), yielding 58% higher hazard than current smoking alone (HR=9.69, 95% CI: 8.04-11.68), while never smokers showed the lowest hazard ratio (HR=0.12, 95% CI: 0.09-0.16). Notably, in a multivariate Cox model adjusted for age, sex, smoking status and estimated pack-years of smoking, AT2 and respiratory epithelium aging signatures retained independent prognostic value for incident lung cancer risk (HR=2.24, 95% CI: 1.73-2.91 and HR=2.10, 95% CI: 1.61-2.74, respectively; Extended Data Fig. 17). The pronounced prognostic value of AT2 cells, which serve as stem cells for repair and regeneration of the alveolus, is consistent with AT2 cells as the cell of origin for lung adenocarcinoma, the most common form of lung cancer. These findings align with literature suggesting that compromised regenerative capacity in the lung parenchyma may create a permissive environment for malignant transformation^80–84^ and may reflect individual differences in lung cancer susceptibility^85^.

For type 2 diabetes, myeloid lineage extreme aging demonstrated the strongest prognostic value (HR=2.62, 95% CI: 2.05-3.33), consistent with a described role of myeloid-derived cytokines in the initiation of an inflamed pancreatic islet microenvironment and increased susceptibility to type 2 diabetes^86^. Importantly, in a multivariate Cox model adjusted for known risk factors including hemoglobin A1C, BMI, smoking status, pack-year history, and renal function, myeloid lineage aging retained prognostic value (HR=1.79, 95% CI: 1.41-2.27, Extended Data Fig. 18).

Moreover, myeloid lineage aging enhanced type 2 diabetes risk stratification beyond glycemic status and identified individuals with normal HbA1c who were nonetheless at elevated risk for future diabetes (Fig. 4b). Though further study is needed, given the efficacy of diabetes interventions and prevention strategies, these findings suggest potential utility of cellular aging signatures for personalized risk stratification and identification of individuals who may benefit from earlier or more aggressive intervention.

A potential limitation to the interpretation is the dependence of plasma protein abundance on overall renal clearance rate and selectivity of the glomerular filtration barrier. However, we observed only modest correlations (generally Spearman rho < 0.2) between cell aging gaps and estimated glomerular filtration rate (eGFR) for individuals with normal or mildly reduced kidney function not meeting criteria for chronic kidney disease (eGFR ≥ 60 mL/1.73m^2^), which comprise the vast majority of the UKB cohort (98.3% of 23,109 individuals with available laboratory values) (Extended Data Fig. 19). Moreover, Cox proportional hazard analyses adjusting for eGFR and albuminuria yielded similar relationships between the cell aging signatures and disease incidence (**Supplementary Tables 8-10**).

### Cellular aging states inform survival and resilience

Beyond cell type-specific disease vulnerabilities, we investigated whether cellular aging signatures have prognostic value for all-cause mortality and overall survival trajectories (Fig. 5). Using Cox proportional hazards regression across over 15 years of UKB follow-up, we identified distinct cellular aging signatures prognostic of mortality risk. The strongest predictors of all-cause mortality were muscle cells (HR=4.38, 95% CI: 4.00-4.80) and skeletal myocytes (HR=4.18, 95% CI: 3.82-4.57), followed by neurons (HR=3.86, 95% CI: 3.53-4.21), fibroblasts (HR=3.73, 95% CI: 3.38-4.11), AT2 cells (HR=3.52, 95% CI: 3.22-3.85), and myeloid lineage cells (HR=3.48, 95% CI: 3.15-3.83), with modest sex differences observed. These associations persisted in models further adjusted for renal function (**Supplementary Tables 8-10**). The prominence of skeletal myocyte aging in mortality prediction is likely linked in part to the importance of overall physical performance status and absence of frailty for survival^87,88^, with neuronal, fibroblast, AT2, and myeloid signatures implicating cognitive, structural, pulmonary, and immune function in longevity.

**Figure 5.**
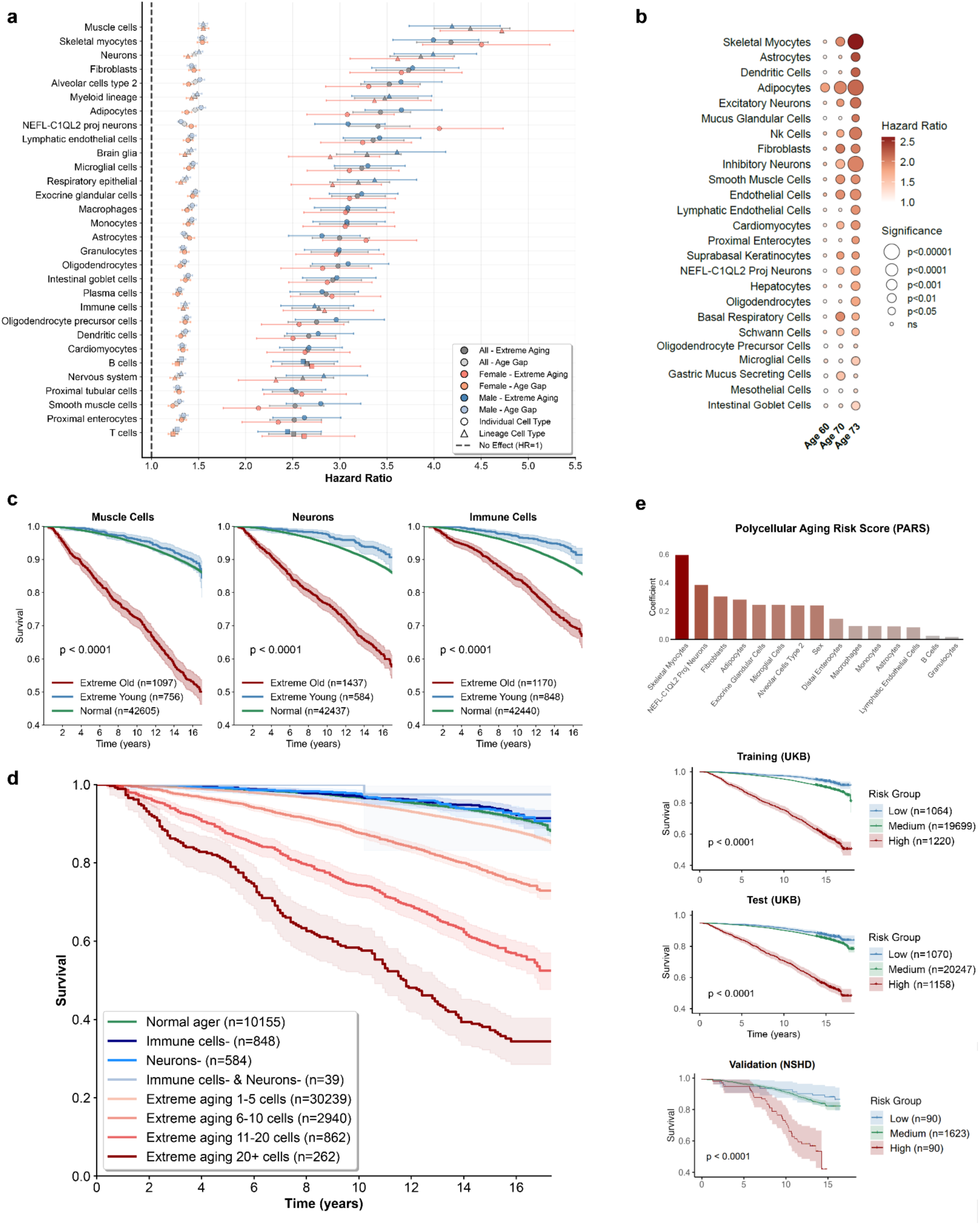
Cellular aging states inform survival and resilience. **a,** Hazard ratios with 95% CI for all-cause mortality by cellular aging over 15 years in the UKB, adjusted for age and sex. Results shown for overall analysis (gray) and sex-stratified analyses (females, red; males, blue). For each cell type, darker markers represent extreme cellular aging and lighter markers represent per-unit increase in age gaps. Vertical dashed line indicates HR=1 (no effect). **b,** Hazard ratios for all-cause mortality by cell type at three timepoints in the NSHD cohort (n=364; ages 63.2±1.1, 70.7±0.7, 72.9±0.6 years; followed to mean age 78.3 years). Color intensity corresponds to hazard ratio magnitude, and dot size corresponds to significance level. **c,** Kaplan-Meier survival curves stratified by cellular aging for muscle cells (left), neurons (middle), and immune cells (right). Extreme agers (red), normal agers (green), youthful agers (blue) in the UKB. Shaded regions represent 95% CI. Sample sizes are indicated in the legend. Shaded regions represent 95% CI. **d,** All-cause mortality by number of extreme aging cell types over 15 years in UKB. Kaplan-Meier curves stratified by extreme aging burden: normal agers (n=10,155, ∼90% survival); 1-5 extreme cell types (n=30,239, ∼85%); 6-10 (n=2,940, ∼73%); 11-20 (n=862, ∼52%); 20+ (n=262, ∼34%). Individuals with youthful immune cells (n=848) or neurons (n=584) regardless of aging in other cell types showed survival similar to or better than normal agers. Shaded regions represent 95% CI. **e,** Polycellular risk score for mortality stratification across plasma proteomics platforms and independent cohorts. Bar plot displays cell type coefficient contributions to the polycellular risk score. Kaplan-Meier survival curves show risk stratification by groups (high, top 5%, red; medium, 90%, yellow; low, bottom 5%, blue) in UKB training cohort (n=21,983), UKB test cohort (n=22,475), and NSHD cohort (n=1,803). Shaded regions represent 95% CI. Sample sizes are indicated in the legend.

We next examined survival stratification by cellular aging status. Kaplan-Meier survival curves demonstrated clear risk stratification, with extreme agers showing markedly reduced survival compared to normal agers (Fig. 5c, Extended Data Fig. 20). Muscle cell extreme aging demonstrated the most pronounced mortality risk, while youthful immune cells and neurons exhibited unique protective effects. These findings indicate that survival depends on maintaining function across multiple physiological systems, with plasma proteomics providing an accessible window into this multidimensional aging process.

Longitudinal tracking of the plasma proteome offers the opportunity to record dynamic molecular events which anticipate physiological decline and death. We assessed the relationship between repeated measures of proteomic cell type aging taken across a 10-year period and subsequent mortality risk in NSHD participants (n=364). We found that the prognostic value of cellular aging signals progressively increased as the follow-up period for mortality drew closer (Fig. 5b). This phenomenon arose prominently for cell types with established connections to motor and cognitive frailty in older age^89,90^, with progressively increasing hazard ratio estimates for skeletal myocytes (Baseline: HR=1.42, 95% CI: 0.97-2.09, Timepoint 2 (T2): HR=1.96, 95% CI: 1.36-2.08, Timepoint 3 (T3): HR=2.79, 95% CI: 1.87-4.16), inhibitory neurons (Baseline: HR=1.64, 95% CI: 1.13-2.37, T2: HR=1.68, 95% CI: 1.24-2.26, T3: HR=2.00, 95% CI: 1.54-2.59) and astrocytes (Baseline: HR=1.14, 95% CI:0.74-1.77, T2: HR=1.50, 95% CI: 0.97-2.33, T3: HR=2.33, 95% CI: 1.45-3.73). This progressive loss of molecular resilience in core motor or cognitive frailty hubs could be a decisive factor in susceptibility to death. At the first timepoint (age ∼ 63), we noted that only adipocyte aging was prognostic of mortality (Baseline: HR=1.83, 95% CI 1.36-2.46), consistent with adiposity as a central determinant of midlife health and a metabolic regulator influencing energy homeostasis, inflammation, and organismal aging^91^. By the third timepoint (age ∼ 73), half of studied cell types (22/43 types, adjusted *p*-value < 0.05) were associated with mortality, with NK cells, endothelial cells, and fibroblasts demonstrating pronounced late-stage amplification of mortality risk. This analysis suggests that biological factors influencing mortality risk change dynamically in a time- and cell-type dependent manner.

We further examined how cumulative burden of cellular aging influences survival and resilience by stratifying individuals by the number of cell types exhibiting extreme aging and visualizing Kaplan-Meier survival curves in the UKB (Fig. 5d). A striking dose-response relationship emerged: individuals with normal aging profiles across all cell types (n=10,155) maintained a survival rate approaching 90%, while those with extreme aging across 20 or more cell types (n=262) experienced dramatically accelerated mortality, with survival declining to approximately 34% after 15 years. The dose-dependent pattern remained across intermediate groups, with extreme aging in 1-5 cell types—the most common aging pattern (n=30,239)—associated with modestly elevated mortality (survival ∼85%), individuals with 6-10 cell types (n=2,940) having further increased risk (survival ∼73%), while those with 11-20 extremely aged cell types (n=862) possessed substantially higher rates of death (survival ∼52%). Notably, individuals with youthful immune cells (n=848) or neurons (n=584) showed survival trajectories similar to, or better than, normal agers, and those youthful in both immune cells and neurons (n=39) exhibited the highest survival, suggesting synergistic protective effects, though the small sample size requires further validation. The prognostic value for mortality of youthful immune cells, youthful neurons, and increasing accumulation of cell types was maintained in multivariate models further adjusted for renal function (Extended Data Fig. 21). The protective effect of maintaining youthful immune and neuronal function suggests these systems may serve key integrative roles in coordinating physiological resilience. Collectively, these findings demonstrate that longevity is shaped by both individual cellular vulnerabilities and the cumulative burden of cellular aging across multiple physiological systems.

The observation that mortality risk depends on both specific cellular vulnerabilities and cumulative aging burden motivated us to develop an integrated “polycellular” aging risk score (PARS). By encoding each individual’s pattern of extreme cellular aging as binary features (extreme aging present or absent for each cell type), we constructed a composite measure of systemic aging burden, using multivariate Cox proportional hazards models. The use of binary extreme aging representation enhances robustness to platform-specific technical variation and facilitates cross-platform generalization. The PARS model was first trained (n=21,983) and tested (n=22,475) in the UKB cohort and further validated in the independent NSHD cohort (n=1,803). Based on the PARS, we stratified individuals into high risk (top 5%), low risk (bottom 5%), and medium risk (all others). Kaplan-Meier survival analysis with log-rank testing revealed clear risk stratification in both the UKB training (*p*<0.0001) and test sets (*p*<0.0001), with high-risk individuals demonstrating markedly reduced survival (Fig. 5e). Critically, when applied to the independent NSHD cohort—which uses the SomaScan platform rather than Olink—the polycellular score maintained significant prognostic value with robust survival curve stratification (*p*<0.0001), despite differences in proteomic technologies and cohort demographics. Consistent with skeletal myocyte aging being the strongest individual cell type predictor of mortality (Fig. 5a), this cell type exhibits the largest coefficient in the PARS, followed by NEFL-C1QL2 projection neurons, fibroblasts, and adipocytes (Fig. 5e). These findings illustrate the prognostic value of the PARS as a platform-agnostic biomarker for mortality and healthspan.

## Discussion

Our study introduces a cell type-specific framework for modeling biological aging patterns with plasma proteomics, leveraging the measurement of over 7,000 plasma proteins across more than 60,000 individuals in three independent cohorts.

Utilizing the putative cell-type proteome, we first described plasma protein expression trajectories that change across lifespan and map to unique cell types. These trajectories formed clusters enriched for cell types and biological pathways that implicate coordinated physiological events. Most notably, we observed a prevalent hepatocyte signature corresponding to age-associated shifts in coagulation and inflammatory processes, consistent with established changes in liver-derived secretory programs with aging^92,93^. Multiple cell-type enrichments also appeared within clusters concurrently, including those of shared developmental lineages, providing insight into the cellular architectures that shape proteomic remodeling during aging. These findings indicate that the plasma proteome captures meaningful signals reflective of physiological events that link to cell types individually and in combination.

When deriving cell type-specific aging models and age gaps, our approach reveals both substantial heterogeneity and coordinated shifts in cellular biological aging patterns, including varied onset of extreme aging that could reflect disease vulnerabilities within specific timeframes across lifespan. Our framework also uncovers significant associations between cellular aging signatures and neurodegenerative diseases, with critical prognostic value for identifying at-risk individuals years before clinical onset. ALS showed the strongest cross-sectional association with skeletal myocyte aging in GNPC; in the UKB cohort with over 15 years of follow-up, extreme skeletal myocyte aging demonstrated pronounced prognostic power for incident ALS, even for cases diagnosed more than three years after baseline assessment, suggesting presymptomatic detection potential.

For AD, GNPC analysis revealed accelerated aging across a broad spectrum of cell types including oligodendrocyte precursor cells, inhibitory neurons, astrocytes, proximal enterocytes, and pancreatic endocrine cells, underscoring the systemic nature of the disease while confirming involvement of key cell types the field has linked to AD^43–45,52–54,94^. Several prominent associations were validated in the NSHD cohort and further supported by analysis of the PACC cognitive score.

Prognostically, astrocyte aging showed the strongest association with incident AD among cellular aging signatures in the UKB cohort, comparable to APOE4 carrier status and exceeding polygenic risk score and chronological age. Intriguingly, astrocyte aging interacted with APOE genotype, with APOE4/4 carriers showing threefold higher incidence with extreme versus youthful astrocyte aging (38.3% versus 12.6% cumulative AD incidence over 15 years). The risk gradient persisted across genotypes, with women more vulnerable than men to the harmful associations of both APOE4 and extreme astrocyte aging. Unexpectedly, youthful astrocytes seem to powerfully neutralize the detrimental effect of carrying one or two APOE4 alleles. Pending independent replication, the biology of this interaction between astrocyte aging – and based on the proteomic signature presumably of a fibrous, matrix rich astrocyte – and APOE might uncover new opportunities to mimic brain resilience.

These findings reciprocally advance aging biomarker science and neurodegenerative research. Neurodegenerative disease incidence is tightly linked to older age^26^, and the strong associations between proteomic cell type signatures and both prevalence and incidence provide face validity that they capture genuine physiology and aging biology. Beyond this, our cellular aging framework sharpens understanding of neurodegenerative processes themselves by contributing a rare human population perspective on their distinct molecular signatures. These conditions, once viewed as relatively isolated from systemic biology, now appear interwoven with gastrointestinal^95^, musculoskeletal^96^, respiratory^97^, and or cardiac^41^ cellular dysregulation^73^. Further exploring these systemic interactions may widen the horizon for therapeutic intervention.

Beyond neurodegenerative diseases, cellular aging signatures have prognostic value for incident cancer and chronic disease. Extreme respiratory cell aging identified smokers at markedly elevated lung cancer risk (58% higher hazard than smoking alone) providing a potential means of targeted surveillance, while myeloid aging identified normoglycemic individuals at an elevated diabetes risk who might benefit from early preventive interventions.

Both specific cellular vulnerabilities and cumulative aging burden influence survival outcomes, with plasma proteomics providing an accessible window into this multidimensional aging process. Skeletal myocyte aging showed the most prognostic value for mortality, followed by neurons, fibroblasts, AT2 cells, and myeloid cells, implicating musculoskeletal, cognitive, structural, pulmonary, and immune function in longevity. A dose-response relationship revealed that individuals with extreme aging across 20+ cell types showed ∼34% survival versus ∼90% for those with normal aging profiles over 15 years. Youthful immune cells or neurons conferred protective effects, suggesting their integrative roles in physiological resilience. The PARS model we introduced maintained robust mortality risk stratification across platforms and cohorts, demonstrating the potential of cellular aging signatures as a platform-agnostic biomarker for healthspan.

Altogether, the non-invasive, blood-based approach introduced in this study enables biological aging characterization at the granularity of the cell, offering a powerful new framework to elucidate aging heterogeneity, underlying disease mechanisms, and potential novel avenues for therapeutic intervention.

In light of these discoveries, we raise several promising directions for future study. In addition to further characterizing biological and chronological aging trajectories across cell types individually and in combination, an important next step is to disentangle causal drivers of cell type-specific aging. Genome-wide association studies hold promise for investigating the genetic architecture of cellular aging signatures and their relationships to disease, potentially uncovering new pathophysiology. Causal inference methods, such as Mendelian randomization, could help pinpoint genetic variants that modulate protein expression and impact cellular aging and disease risk. These insights could identify novel therapeutic targets that alter cellular aging trajectories. Another intriguing direction is exploring the temporal dynamics and sequential progression of cellular aging. While we describe protein expression trajectories for individual cell types, future work should investigate the timing and order of biological events to better contextualize disease progression and onset. Moreover, our analyses were designed to capture normative, population-level aging patterns, inclusive of heterogeneous health states and lifestyle factors that characterize real-world aging. To further disentangle cellular aging signatures from the potential influence of systemic comorbidities, complementary analyses can curate ‘superager’ cohorts—individuals free of chronic disease—to reveal protective cellular trajectories underlying exceptional longevity and resilience. Finally, since the demographic composition of the GNPC, UKB, and NSHD cohorts is predominantly Caucasian and skewed towards older individuals, it will be important to include younger and more diverse populations to capture early-life cellular aging trajectories and extend the generalizability of our models.

We acknowledge the limitations of our study. One such limitation is that the study was restricted to the cell types cataloged in the Human Protein Atlas (HPA) single-cell transcriptomics dataset. While the HPA serves as a resource for a range of cell types and over 20,000 genes likely to capture aging effects, certain disease-relevant cell types, including specialized neuronal subpopulations, pericytes, and ependymal cells^98^, are absent or underrepresented. We also acknowledge that transcript levels do not always correlate directly with protein abundance, and that cell-type specificity thresholds derived from transcriptomic enrichment may benefit from further refinement. Additionally, plasma proteins may arise through diverse processes including secretion, extracellular vesicle traffic, and tissue leakage that are not explicitly disentangled. However, our framework intentionally captures a holistic view of cellular aging encompassing intrinsic aging and damage-related signals, both of which are essential for predicting disease risk and resilience. We view our work as a new framework that leverages plasma proteomics as a non-invasive window into cellular aging, with significant potential for continuous refinement as understanding of cell-type signatures advances and single-cell transcriptomic and proteomic resources expand. Future studies could leverage the rapidly expanding landscape of human single-cell atlases, such as the Allen Human Brain Atlas, while integrating domain-specific knowledge to refine cell type-specific aging models. Likewise, deeper coverage of the plasma proteome, including the quantification of protein isoforms, will increase the number of cell types that can be linked to plasma proteins.

In summary, our study demonstrates that large-scale plasma protein profiling integrated with machine learning techniques enables non-invasive assessment of cell type-specific aging in living human populations. By characterizing cellular aging signatures in the plasma proteome, we achieve granular delineation of aging patterns across distinct cellular populations. Our framework uncovers significant links between cellular aging, disease, and mortality, with implications for understanding the biological underpinnings of longevity, disease mechanisms, and identification of novel therapeutic targets.

## METHODS

### Proteomics Measurements

#### SomaScan Proteomics

The SomaLogic (https://SomaLogic.com/) SomaScan assay^99,100^, which uses slow off-rate modified DNA aptamers (SOMAmers) to bind target proteins with high specificity, was used to quantify the relative concentration of thousands of human proteins in the GNPC, KADRC, and NSHD cohorts. The v4.1 (7,289 proteins) assay was used for all of the mentioned cohorts and samples except the NSHD which utilized the v5.0 assay (11,742 proteins). Standard SomaLogic normalization, calibration, and quality control were performed on all samples, resulting in protein measurements in relative fluorescence units (RFU). Briefly, pooled reference standards and buffer standards are included on each plate to control for batch effects during assay quantification. Samples are normalized within and across plates using median signal intensities in reference standards to control for both within-plate and across-plate technical variation. Samples were further normalized to a pooled reference using an adaptive maximum likelihood procedure. Samples were flagged by SomaLogic if signal intensities deviated significantly from the expected range and were subsequently removed. The resulting values were the provided data from SomaLogic and were considered “raw” data. Downstream preprocessing and quality control were handled by GNPC and its respective cohorts. To facilitate the usage of GNPC plasma proteomic clocks trained on 7,289 (v4.1) SomaScan assay in the NSHD cohort, v5.0→v4.1 scaling factors provided by the SomaLogic package SomaDataIO lift_adat() were applied to raw abundance values before age estimates were generated.

#### Olink Proteomics

The Olink Explore 3072 proximity extension assay (PEA) platform was used to quantify plasma proteins in the UK Biobank. Olink is based on the binding of two polyclonal antibody pools to a target protein and subsequent hybridization and enrichment of two unique single-stranded DNA probes. The assay encompasses 2,941 immunoassays targeting 2,923 proteins. If matching pairs of antibodies bind to the protein, the attached oligonucleotides hybridize, and are then measured using sequencing. Olink measurements were based on NPX values recommended by the manufacturer, which includes normalization. Additional details on Olink proteomics data quality control, acquisition, and handling are well-documented^101^; further details are available at https://biobank.ndph.ox.ac.uk/ukb/ukb/docs/PPP_Phase_1_QC_dataset_companion_doc.pdf.

### Study Population and Clinical Phenotypes

#### Global Neurodegeneration Proteomics Consortium (GNPC)

The GNPC v1.3 dataset (https://www.neuroproteome.org/new-harmonized-data-set) is a large-scale global neurodegenerative disease biomarker discovery effort that hosts the largest collection of SomaScan data – spanning over 40,000 patient samples from over 20 international research groups. Data from the GNPC include proteomic samples from clinically healthy individuals, and those diagnosed with Alzheimer’s disease (AD), Parkinson’s disease (PD), amyotrophic lateral sclerosis (ALS), mild cognitive impairment–subjective cognitive impairment (MCI-SCI), and frontotemporal dementia (FTD). All cohorts and data were anonymized with letter codes by GNPC prior to analysis. A standard sample collection procedure was utilized, with potential site-to-site variation guided by the discretion of individual cohorts. All samples underwent standardized preparation and processing protocols and were stored at −80°C until proteomic profiling was performed. Ethics approval for each cohort was obtained from the respective Institutional Review Boards (IRB), and written informed consent was obtained from all participants or their legally authorized representatives. The study design was approved by each respective participating institution.

Our initial analysis included 21,979 individuals in GNPC and was subsequently reduced to account for reporting completeness and consistency. Specifically, our analysis was applied to version 4.1 (7,289 protein targets) of the SomaScan assay, and only included individuals with complete age and sex information. Due to known variations in sample quality and detection capability for proteins collected in other media including serum and CSF, we limited our analysis to plasma samples collected in ethylenediaminetetraacetic acid (EDTA) by routine venipuncture. As a result of these quality control measures, 14,281 individuals from 14 independent cohorts were prioritized for downstream analysis (**Supplementary Table 3**). Among these individuals, 7,074 were categorically defined as healthy. In agreement with the definition proposed by GNPC, we define the healthy population as individuals with no diagnosis of AD, PD, FTD, MCI, or ALS, and a Clinical Dementia Rating (CDR) score not exceeding zero.

#### Knight Alzheimer’s Disease Research Center (KADRC)

The Knight-ADRC cohort, one of the contributing cohorts in GNPC, is a National Institute on Aging (NIA)-funded longitudinal observational study of clinical dementia subjects and age-matched controls. Research participants at the KADRC undergo longitudinal cognitive, neuropsychological, imaging, and biomarker assessments including CDR testing. The Institutional Review Board of Washington University School of Medicine in St. Louis approved the study, and research was performed in accordance with the approved protocols. The KADRC includes samples of individuals with AD and healthy controls. Blood collection and processing were performed according to a rigorous standardized protocol to minimize variation associated with blood draw and processing. Briefly, ∼10 cc of whole blood was collected in four vacutainer ethylenediaminetetraacetic acid (EDTA) tubes (Becton Dickinson vacutainer EDTA tube) and spun at 1800 x g for 10 minutes to separate plasma, leaving 1 cm of plasma above the buffy coat, with care taken not to disturb the buffy coat and circumvent cell contamination. Plasma was aliquoted into polypropylene tubes and stored at −80°C. Plasma processing times averaged approximately one hour from the time of the blood draw to the time of freezing and storage. All blood draws were performed in the morning to minimize the impact of circadian rhythm on protein concentrations.

#### National Survey of Health and Development (NSHD)

The National Survey of Health and Development (NSHD) is the world’s oldest continuously followed birth cohort, consisting of a geographically diverse sample of UK individuals born in a single week in March 1946. Participants underwent plasma proteomics measurement with the SomaScan 11k (v5.0) proteomics assay at three timepoints corresponding to mean ages 63.2±1.1 years (n=1,803), 70.7±0.7 years (n=483), and 72.9±0.6 years (n=396). 364 participants had proteomics samples collected at all three timepoints, enabling longitudinal profiling. One participant was removed from analysis at the second timepoint and one at the third timepoint due to incomplete participant ID reporting. Two duplicated entries at the third timepoint were excluded. Participant survival data were extracted from the NHS England and the NHS Central Register via NHS Digital in June 2024. Participant survival was tracked to an approximate age of 78. Individuals who withdrew from digital health data collection were censored (n=2/1,803). A subgroup (n=483) participated in the NSHD’s neuroimaging substudy Insight-46 and underwent comprehensive neurological phenotyping. Plasma concentrations of phosphorylated tau at threonine-217 (pTau-217; pg/mL) were quantified using the ALZpath Simoa assay (Quanterix), a high-sensitivity single-molecule immunoassay that specifically detects the pTau-217 epitope associated with Alzheimer’s disease pathology. Cognitive performance was assessed using the Preclinical Alzheimer Cognitive Composite (PACC), a collection of neuropsychological tests sensitive to early cognitive changes in preclinical Alzheimer’s disease.

#### UK Biobank (UKB)

The UK Biobank is a population-based prospective observational cohort with omics and phenotypic data collected from approximately 500,000 participants, aged 40 to 69 years, spanning a recruitment period between 2006 and 2010. Participants underwent baseline assessments including physical measurements, questionnaires, and biological sample collection, with longitudinal follow-up for disease outcomes and mortality extending over 15 years. All participants provided informed consent.

The UKB Pharma Proteomics Project (UKB-PPP) consortium generated Olink Explore 3072 proteomics data from blood plasma samples collected from ∼54,000 UKB participants at baseline. After UKB-PPP quality control, the baseline data included 2,923 proteins measured in ∼53,000 samples. We performed additional quality control steps: samples with more than 1,000 missing protein values (n=8,182) were removed; samples with discordant reported and genetic sex (n=373) were excluded; and proteins with missing values in over 20% of samples (n=7) were removed. This resulted in a post-quality control dataset of 44,458 samples with 2,916 protein measurements. Data were randomly split into train and test sets based on assessment centers (train: n = 21,983; test: n = 22,475), with protein values z-score normalized based on the means and standard deviations of the training set. Missing protein values (2.7% of total measurements) were imputed using k-nearest neighbors imputation. We trained a k-nearest neighbors imputer (k=148, set to the square root of train sample size) using scikit-learn’s KNNImputer function. Imputation performance was validated on a subset of samples with no original missing values by randomly inserting missing values and comparing imputed to true values, yielding a mean absolute error of 0.57, confirming robust imputation performance. UKB data were analyzed under application number 184058.

Disease diagnosis dates were collated from UKB “First Occurrences” (data category 1712), “Cancer Register” (data category 100092), and, where available, “algorithmically defined outcomes” (data category 47; e.g. for AD, PD, ALS, FTD, stroke). If multiple dates were recorded, the earliest date was used. For incident disease analyses, individuals with diagnosis prior to the date of plasma collection were excluded from the corresponding analysis. The following “First Occurrences” or “Cancer Register” ICD-10 (and where available ICD-9) codes were used: heart failure (I42, I43, I50); stroke (I60-64); COPD (J41-44); type 2 diabetes (E11, E14); lymphoma (C81-89, and ICD-9 2014-2016, 2019, 2020); lung cancer (C33, C34, and ICD-9 1623, 1629); AD (F00, G30); PD (G20). For calculation of type 2 diabetes incidence, all individuals with diagnosis of type 1 diabetes (ICD-10 E10) or other uncommon forms of diabetes (“other specified diabetes” ICD-10 E13) were removed from analysis. Type 2 diabetes diagnosis date was derived from the earliest of diagnosis date from the “First Occurrences” category or first date outpatient hemoglobin A1c value was greater or equal to 6.5%. Individuals diagnosed with dementia without specified cause and without evidence of alternative etiology (neurodegenerative disease of the basal ganglia, ALS, multiple sclerosis, syphilis, HIV, B12/folate/thiamine deficiency, frontotemporal dementia, hydrocephalus, alcohol use disorder (either diagnosed or meeting criteria via reported consumption: greater than 28 units/week for women or 35 units/week for men), cerebrovascular pathology, sedative abuse, ICD-10 A81 “atypical virus infections of central nervous system”, or ICD-10 G32 “other degenerative disorders of nervous system”) were assigned a diagnosis of AD given its status as the predominant cause of dementia. Individuals with competing diagnoses (e.g., FTD and AD or progressive supranuclear palsy and PD) were classified as having only the more rare diagnosis given the high clinical index of suspicion needed for formal diagnosis. Individuals with non-physiologic diagnosis dates (i.e., year of birth) were removed from disease incidence analysis of the corresponding disease.

The following variables were extracted from UKB baseline assessment: smoking status (data field 20116_i0), pack-years of smoking (data field 20161_i0), hemoglobin A1c (data field 30750_i0), body mass index (BMI) (data field 21001_i0), waist circumference (data field p48_i0), alcohol consumption (aggregated over data fields 1568_i0, 1578_i0, 1588_i0, 1598_i0, and 1608_i0), sleep duration (data field 1160_i0), physical activity (data field 884_i0 and 904_i0), plasma creatinine (data field 23478_i0), plasma cystatin C (data field 30720_i0), urine creatinine (data field 30510_i0), urine albumin (data field 30600_i0), APOE genotype (data field 2315), and AD polygenic risk score (PRS, data field 26206). Healthy lifestyle was defined as meeting all of the following criteria: never smoking, no regular alcohol consumption, BMI < 25, waist circumference < 90 cm for men or < 84 cm for women, at least 5 days per week of 10+ minutes of moderate or vigorous physical activity, and ≥7 hours of sleep per night. Details on available phenotypes can be found at https://biobank.ndph.ox.ac.uk/showcase/.

#### Identification of cell type-enriched plasma proteins

The Human Protein Atlas single-cell RNA-seq database was used to map the putative cell type-specific plasma proteome. The single-cell transcriptomics dataset utilized in this study encompassed more than 60 cell types and over 20,000 genes (normalized transcripts per million, nTPM; a description of all included cell types is provided in Extended Data Fig. 2) (https://www.proteinatlas.org/humanproteome/, Human Protein Atlas version 24.1). Hofbauer, Langerhans, and Kupffer cells were pooled into a broader macrophage categorization. Sex-specific cell types were excluded from the study. Guided by prior studies, we classified genes as “cell-type-enriched” if they were expressed at least twofold higher in one cell type compared to any other cell type or expressed exclusively in a single cell type. Cell-type-enriched genes were mapped to the 7,289 plasma proteins quantified in the v4.1 SomaScan assay for SomaScan aging clock models, and to the 2,923 plasma proteins quantified in the Olink Explore 3072 assay for Olink aging clock models. Of note, while gene expression in tissues does not guarantee secretion into plasma, studies have shown that a substantial proportion of cell-type-enriched transcripts correspond to detectable proteins in peripheral blood^102^.

#### Estimation of plasma protein trajectories across lifespan

To estimate plasma protein trajectories with respect to chronological age, plasma protein expression was *z*-scored, and LOWESS regression was fitted for each plasma protein with the smoothing parameter set to frac=0.3 (such that 30% of the data were used in each local fit) using the *lowess* function from the *statsmodels* Python package. These data were applied for heatmap construction using the *scipy.cluster.hierarchy* module from *SciPy*. For clustering analysis, similar trajectories were grouped by computing pairwise differences based on the Euclidean distance metric, and hierarchical clustering was performed using Ward’s method. The elbow method was used to determine the optimal number of clusters, resulting in a threshold parameter of 0.05 in the *scipy.cluster.hierarchy* function and 33 distinct clusters.

#### Trajectory clustering with cell-type and pathway enrichment analysis

To identify cell types that were enriched in clusters, we performed hypergeometric enrichment analysis. For each cluster, we assessed the overlap between cluster proteins and predefined sets of proteins mapped to individual cell types. For each cell type, a hypergeometric test was used to compute the probability of observing the actual or greater overlap by chance, given the size of the cluster, the size of the mapped protein set, and the size of the overall background. To account for multiple testing, false discovery rate (FDR) correction was applied with the BH procedure using the *stats.multitest.multipletests* function in the *statsmodels* Python package. Cell types enriched in a cluster were defined as those meeting the significance threshold with adjusted *p*-value < 0.05. Cell types meeting this threshold were plotted in a dot plot, with dot size and opacity representing enrichment significance. Organs were incorporated into this analysis using an identical method. Mapping of the organ-specific proteome was performed utilizing methods from prior studies and publicly available datasets provided by the Genotype Tissue Expression (GTEx) project^21,103^.

To interpret the biological relevance of each cluster, we performed pathway enrichment analysis. For each cluster, we queried *g:Profiler* using the *GProfiler* Python package (*gprofiler-official*) with the human gene namespace (organism=’hsapiens’) and restricted enrichment sources to Gene Ontology (GO), Biological Process (GO:BP), Molecular Function (GO:MF), Cellular Component (GO:CC), as well as Kyoto Encyclopedia of Genes and Genomes (KEGG), and Reactome pathways. To correct for multiple hypothesis testing across all cluster-level enrichments, the BH procedure was applied, with adjusted *p*-values reported.

#### Cellular age estimation and age gap calculation

To estimate biological age using plasma proteomics, we developed cell type-specific aging models using elastic net regression with the *glmnet* R package. Each aging model was trained on a distinct set of cell-type-enriched plasma proteins to predict chronological age. We implemented bootstrap aggregation, generating 100 bootstrap samples through resampling with replacement from our training data. For each bootstrap sample, we trained a model on *z*-scored log-transformed protein abundance values with sex (F=1, M=0) as an additional covariate to predict chronological age. Hyperparameter tuning of the L1 regularization parameter (λ) was performed using 10-fold cross-validation with the *cv.glmnet* function. The final predicted age for each sample was calculated as the mean prediction from the 100 bootstrap models. We applied this approach to 60 cell types, excluding 15 sex-specific cell types (e.g. spermatocytes, oocytes, prostatic glandular cells, and ovarian stromal cells) to ensure representation of both sexes in cell-type biological age estimation. For the SomaScan platform, models were trained on the healthy individuals in the KADRC cohort and subsequently applied to the broader GNPC cohort for disease association analyses and to the independent NSHD cohort for external validation. Aging models were evaluated based on performance and robustness criteria. Models were excluded if they exhibited insufficient predictive performance (correlation coefficient r < 0.25 in the training set or r < 0.15 when applied to the GNPC cohort) or contained fewer than four protein features, which may limit the robustness of downstream inferences due to the inherent technical noise in proteomic assays. After applying these quality control criteria, 43 cell type-specific aging models were retained for downstream analyses.

For the Olink platform, models were trained on 21,983 individuals from the UKB training set and subsequently applied to the UKB test set (n=22,475). Since the Olink dataset included 2,916 proteins after quality control—fewer than the 7,289 proteins measured by SomaScan—and generally yields fewer signatures across cell types, we additionally developed 14 lineage-level cell-type models to capture aging signals at broader ontological categories. To demonstrate the capability of our framework to model cellular aging at multiple levels of the cell-type ontology, we extended our analysis to include 14 parent lineage cell types:

*Immune cells*: Lymphoid lineage (B cells, plasma cells, T cells, NK cells); Myeloid lineage (monocytes, granulocytes, macrophages, dendritic cells); All immune cells (combining all lymphoid and myeloid cell types).

*Nervous system*: Neurons (excitatory neurons, inhibitory neurons); Brain glia (astrocytes, oligodendrocytes, oligodendrocyte precursor cells, microglial cells); Retinal cells (cone photoreceptor cells, rod photoreceptor cells, bipolar cells, horizontal cells, Müller glia cells); All nervous system (all neuronal and glial cell types, including Schwann cells).

*Endothelial cells*: Endothelial cells and lymphatic endothelial cells.

*Glandular cells*: Breast glandular cells, exocrine glandular cells, prostatic glandular cells, pancreatic endocrine cells, mucus glandular cells, serous glandular cells, salivary duct cells, and secretory cells.

*Epithelial cells*: Respiratory epithelial (alveolar type 1 cells, alveolar type 2 cells, basal respiratory cells, ciliated cells, club cells, ionocytes); Skin epithelial (basal keratinocytes, suprabasal keratinocytes, melanocytes, basal squamous epithelial cells, squamous epithelial cells); Gastrointestinal epithelial (distal enterocytes, proximal enterocytes, intestinal goblet cells, Paneth cells, enteroendocrine cells, gastric mucus-secreting cells); Urogenital epithelial (collecting duct cells, distal tubular cells, proximal tubular cells).

*Muscle cells*: Skeletal myocytes, cardiomyocytes, and smooth muscle cells.

For each lineage cell type, we identified protein signatures uniquely enriched within the lineage, defined as genes with average expression at least twofold higher than any cell type outside the lineage. This approach captures shared lineage-level aging signals while maintaining biological interpretability. After applying the same quality control criteria (correlation coefficient r ≥ 0.25 in training, r ≥ 0.15 in test; minimum four protein features), 48 cell type-specific aging models were retained for Olink-based analyses.

To calculate individual age gaps for each cell type-specific aging model, we fit a local regression between predicted and chronological age using the *lowess* function from the *statsmodels* Python package with the fraction parameter set to 2/3 to estimate the population mean. Local regression was used instead of linear regression based on evidence supporting nonlinear changes of the plasma proteome with age. We derived age gaps for each cohort based on the fit of healthy individuals in the corresponding cohorts to account for cohort differences and excluded two cohorts (out of 16) that lacked healthy individuals in GNPC. Individual sample age gaps were calculated as the residual between predicted age and the LOWESS regression estimate of the population mean for the corresponding chronological age. Age gaps were *z*-scored per aging model to account for differences in model variability and facilitate comparison across cell types in downstream analyses. Extreme agers were defined as individuals with an age gap *z*-score absolute value greater than 2 for a given aging model.

#### Cellular aging correlation analyses and network visualization

We examined correlations of age gaps across cell types using Pearson correlation with the *pearsonr* function from the *SciPy* Python package. In the network visualization, each node represents a specific cell type, colored according to its cellular category. Edges connect cell type pairs with age gap correlations exceeding a threshold of 0.35. Edge width corresponds to correlation strength, with green edges highlighting the top 15 correlations.

#### Extreme aging stability analysis

The stability of extreme aging cellular states was evaluated in the NSHD across three timepoints spanning a 10-year period (n=364; mean ages 63.2±1.1 years, 70.7±0.7 years, and 72.9±0.6 years), wherein the retention of extreme agers from baseline to the final timepoint was quantified. Only individuals with proteomic samples at all three timepoints were included in the extreme aging stability analysis.

### Association analysis between age gaps and neurodegenerative disease status

Associations between cell type-specific age gaps and neurodegenerative diseases were evaluated using point-biserial correlation with the *pointbiserialr* function from the *SciPy* Python package. The analysis included the five neurodegenerative and neurological conditions present in the GNPC: AD (n=2,761), ALS (n=245), PD (n=476), FTD (n=199), and MCI-SCI (n=1,992).

Only cohorts with at least 10 disease diagnoses and healthy controls were included in the corresponding analyses. For each disease analysis, healthy individuals from the relevant cohorts containing the disease cases served as controls. P-values were adjusted for multiple hypothesis testing using the BH procedure.

#### Prognostic analysis of incident disease

We assessed the prognostic value of cellular aging signatures for incident disease in the UKB cohort using Cox proportional hazards regression models. Time-to-event data were analyzed with follow-up extending over 15 years from baseline proteomic assessment. Participants were followed until the first occurrence of the disease of interest, death, or last update of clinical data, whichever occurred first. Individuals with disease diagnosis prior to plasma collection were excluded from the corresponding analysis. For each cell type-disease pair, we computed hazard ratios (HR) and 95% confidence intervals (CI) for both continuous age gaps (per unit increase in z-scored age gap) and extreme aging status. All Cox models were adjusted for chronological age and sex as covariates. For specific disease analyses, multivariate models were further adjusted for additional factors (e.g., hemoglobin A1C, BMI, smoking status, pack-years, and renal function for type 2 diabetes). P-values were adjusted for multiple hypothesis testing using the BH procedure with a significance threshold of 0.05.

For AD, we compared the prognostic value of cellular aging signatures against established risk factors. Hazard ratios were computed for APOE4 carrier status, APOE2 carrier status, AD PRS, chronological age, astrocyte aging status, and APOE combined with astrocyte aging status. Sex-stratified analyses were performed by fitting separate Cox models for females and males to assess potential sex differences in AD risk associations. For ALS, a sensitivity analysis was performed considering only cases diagnosed more than three years after blood draw to assess whether the prognostic value was retained.

Cumulative incidence curves were estimated using the Kaplan-Meier method and visualized for key cell type-disease associations stratified by aging status. Differences in survival between groups were assessed using log-rank tests.

#### Association of cellular aging with AD pathology and cognitive performance

We assessed whether cell types linked to AD diagnosis in the GNPC cohort were associated with biological evidence of AD pathology and cognitive performance in the NSHD Insight-46 substudy (n=483). Using AD-relevant measures available in the study, we applied linear regression models adjusted for sex and chronological age to evaluate the association between z-scored cellular age gaps and pTau-217 (pg/mL) burden using the *stats* package in R. Plasma pTau-217 was measured using the ALZpath assay on the Single Molecule Array (Simoa) HD-X platform^70^. For cognition, we applied the same linear regression model to evaluate the association between cellular age gaps and the Preclinical Alzheimer Cognitive Composite composite score sensitive to early cognitive decline in preclinical AD^104^. The analysis was applied to the 22 cell types linked to clinical AD in the GNPC cohort. P-values were adjusted for multiple hypothesis testing using the BH procedure.

#### Extreme ager odds ratio analysis

We assessed the association between extreme aging in specific cell types and disease status by calculating odds ratios using Fisher’s exact test. For each cell type-disease pair, we constructed contingency tables comparing the frequency of extreme agers (*z*-scored age gap greater than 2) versus non-extreme agers among disease cases versus healthy controls. Fisher’s exact test was performed using the *fisher_exact* function from the *SciPy* Python package. The Haldane-Anscombe continuity correction was applied by adding 1/2 to each cell in the contingency tables to mitigate potential bias from zero counts. The 95% confidence intervals for odds ratios were computed using Woolf’s method. FDR for multiple hypothesis testing was conducted using the BH procedure, with a significance threshold of 0.05.

#### All-cause mortality analysis

We evaluated the association between cellular aging signatures and all-cause mortality in the UKB cohort using Cox proportional hazards regression models adjusted for chronological age and sex. For each cell type, we computed hazard ratios (HR) and 95% confidence intervals (CI) for both extreme aging status and continuous age gaps. Sex-stratified analyses were performed by fitting separate Cox models for females and males, adjusted for chronological age. Results were visualized using forest plots, with separate estimates shown for overall, female, and male populations. P-values were adjusted for multiple hypothesis testing using the BH procedure with a significance threshold of 0.05.

Kaplan-Meier survival curves were generated to visualize mortality risk stratified by cellular aging status (extreme old, normal, and extreme young) for key cell types. Differences in survival between groups were assessed using log-rank tests. To examine the cumulative burden of cellular aging on survival, we stratified individuals by the total number of cell types exhibiting extreme aging and generated Kaplan-Meier curves for each stratum.

#### Longitudinal mortality risk analysis in the NSHD

We used the NSHD cohort to compute sequential estimates of mortality risk over a 10-year period using age gap estimates of same chronological aged individuals at three timepoints (n=364; mean ages 63.2±1.1 years, 70.7±0.7 years, and 72.9±0.6 years). Cox proportional hazards models were used to examine associations between cellular age estimates and mortality risk over approximately 15 years of follow-up, with censoring for withdrawal from follow-up (n=2) or death (n=281) and adjustment for sex. For each cell type, we computed hazard ratios (HR) and 95% confidence intervals (CI) for continuous age gaps. Multiple comparisons were corrected using the BH procedure. Results were visualized as a dot plot heatmap, with each timepoint represented as an x-axis label, dot hue and size reflecting HR magnitude and significance level, respectively.

#### Development and cross-platform validation of the polycellular aging risk score (PARS)

To develop an integrated mortality risk prediction model, we constructed a polycellular aging risk score (PARS) using binary representations of extreme aging status across all cell types. The use of binary extreme aging features (present/absent) rather than continuous age gap values enhances robustness to platform-specific technical variation and facilitates cross-platform generalization. Multivariate Cox proportional hazards regression was performed on the UKB training set (n=21,983; Olink) with extreme aging status for each cell type as binary covariates. The resulting model coefficients were used to calculate the PARS for each individual. Based on the score distribution, individuals were stratified into three risk groups: high risk (top 5%), medium risk (middle 90%), and low risk (bottom 5%). To demonstrate generalizability and platform-agnostic utility, the PARS was validated in the UKB test set (n=22,475; Olink) and the independent NSHD 1946 cohort (n=1,803; SomaScan) using the same criteria for mortality risk stratification. Kaplan-Meier survival curves were generated for each risk group, and differences in survival were assessed using log-rank tests.

#### Statistical analyses

Pearson correlation was used to examine the correlation of age gaps across cell types using the *pearsonr* function from the *SciPy* Python package. Pairwise t-tests were performed to compare cellular age gaps between APOE genotype groups using the *ttest_ind* function from *SciPy*. Associations between cell type-specific age gaps and neurodegenerative diseases were evaluated using point-biserial correlation with the *pointbiserialr* function from *SciPy*. FDR correction for multiple hypothesis testing was applied using the BH procedure in all relevant statistical analyses with the *multipletests* function from the *Statsmodels* Python package. The significance threshold for adjusted p-values was set at 0.05.

## DATA AVAILABILITY

GNPC data is available upon request to qualified researchers through a standard protocol (https://www.neuroproteome.org/harmonized-data-set-hds) and will be publicly available at the time of publication. Knight-ADRC proteomics data were generated by the laboratory of principal investigator Carlos Cruchaga and are available upon reasonable request to The National Institute on Aging Genetics of Alzheimer’s Disease Data Storage Site (NIAGADS) https://www.niagads.org/knight-adrc-collection. UK Biobank data are available upon request to qualified researchers through a standard protocol (https://www.ukbiobank.ac.uk/register-apply). Bona fide researchers can apply to access the NSHD data via a standard application procedure 889 (further details available at https://skylark.ucl.ac.uk/NSHD/access/). Mortality data can be 890 requested from the UK Longitudinal Linkage Collaboration (https://ukllc.ac.uk/).

## CODE AVAILABILITY

A GitHub repository entitled *cellage* will become available at the time of publication supporting cell-type proteomic aging clocks and biological age estimation.

## AUTHOR CONTRIBUTIONS

D.Y.D., V.A.B., R.P., and T.W.-C. conceptualized the study. D.Y.D., V.A.B., and K.L.C. led study design and analyses. J.G. and D.M. aided in study design and analyses. A.F., H.S.-H.O., V.W., N.L., and A.I. provided data, aided in analyses, and/or provided insights. C.C. established the Knight-ADRC cohort. D.Y.D., V.A.B., K.L.C., and J.G. produced figures and wrote the manuscript. T.W.-C. edited the manuscript. T.W.-C and J.M.S. supervised the study. All authors critically revised the manuscript for intellectual content. All authors read and approved the final version of the manuscript. T.W-C., D.Y.D, V.A.B, and Stanford University have filed a patent application related to this work.

## ACKNOWLEDGEMENTS

We thank the staff and participants in all studies for their important contributions. We thank A. Poyi-Tsai, E.K. Costa, P. Moran-Losada, K. Ying, M. Reich and other members of the Wyss-Coray laboratory for feedback and support and D. Channappa for laboratory management. We thank A. Keshavan, K. Lu and J. Nicholas and other members of the NSHD study team for feedback and support. We thank Ashvini Keshavan for advice on analysis of dementia-related biomarkers, Jennifer Nicholas for statistical guidance and the Insight 46 team for their roles in data acquisition.

Funding: This work was supported by the Stanford Alzheimer’s Disease Research Center (National Institute on Aging grants P50AG047366 and P30AG066515), the National Institute on Aging (AG072255, T.W.-C), the Milky Way Research Foundation (T.W.-C.), Nan Fung Life Sciences (T.W.-C.), the Knight Initiative for Brain Resilience (T.W.-C.), AHA-Allen Brain Health and Cognitive Impairment Cross-Network Collaborative Grants (23BHCICG1188316, N.L.), MAC3 Impact Philanthropies (MAC3 Dementia and Ageing Fellow, N.L.), Alzheimer’s Research UK (ARUK-PG2014-1946 and ARUK-PG2017-1946 J.M.S.), Alzheimer’s Association (SG-666374-UK BIRTH COHORT J.M.S), British Heart Foundation (J.M.S.) and UK Dementia Research Institute through UK DRI Ltd J.G. is an ARUK clinical research fellow.

---

**Extended Data Figure 1:**
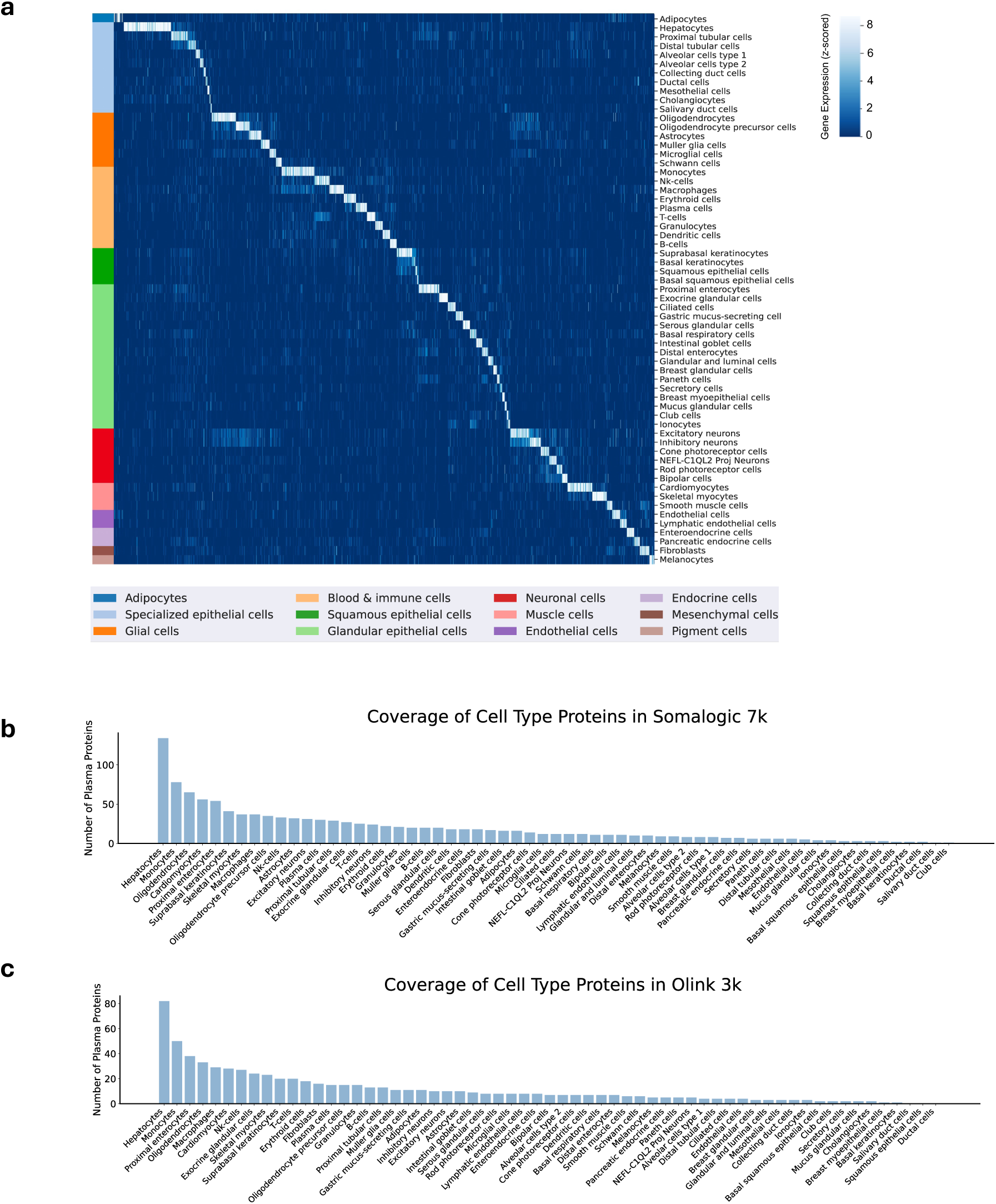
Identification of cell type-specific plasma proteins. **a**, Z-scored gene expression of cell type-enriched plasma protein encoding genes in the Human Protein Atlas (HPA, version 24.1) single-cell transcriptomic dataset, labeled by cell type and grouped according to HPA cell type family categorizations. **b**, Coverage of cell type-specific proteins across the 7,289 proteins measured by the 7k SomaLogic SomaScan assay. **c**, Coverage of cell type-specific proteins across the 2,923 proteins measured by the 3k Olink assay.

**Extended Data Figure 2:**
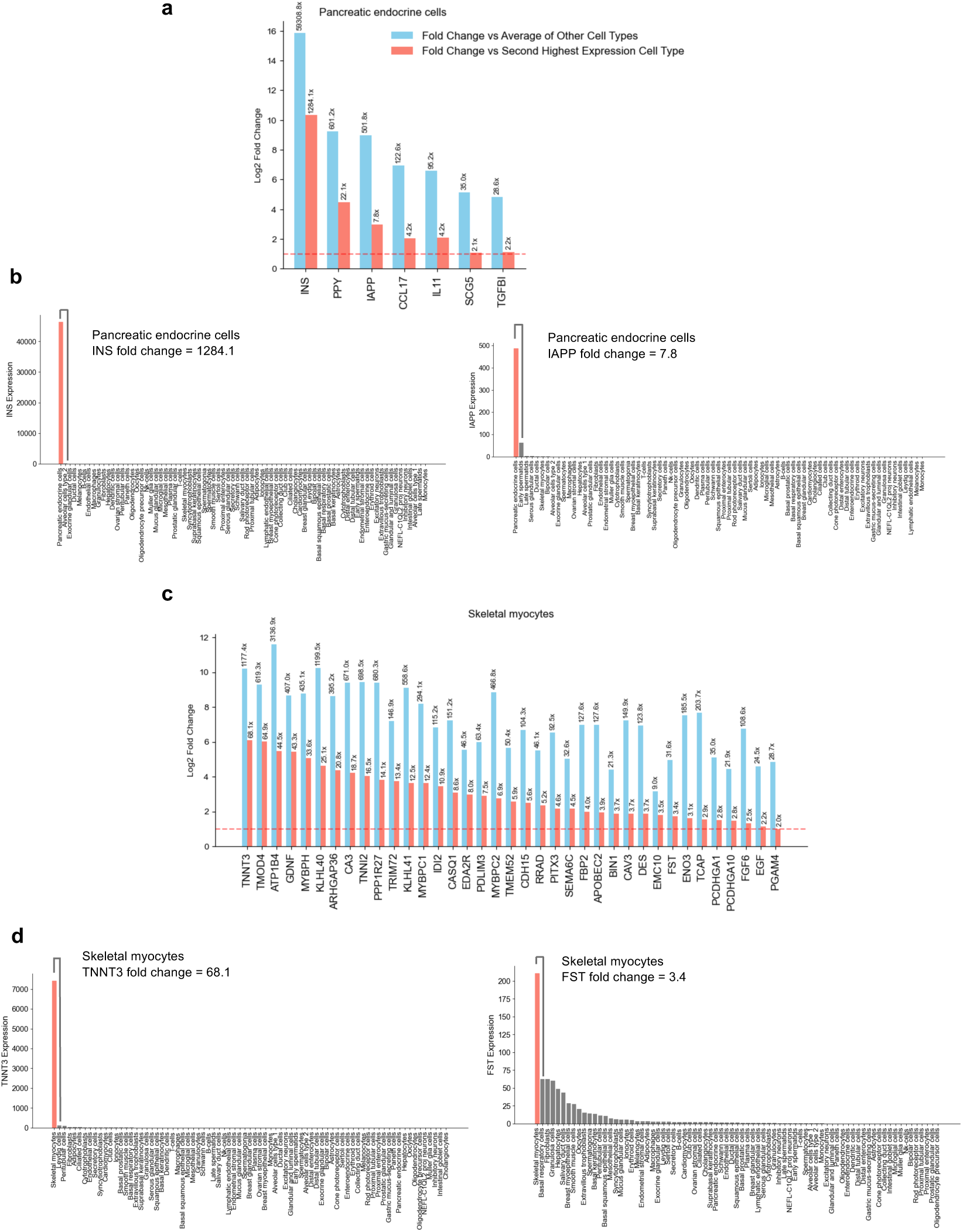
Enrichment profile examples of cell type-specific proteins. **a**, Fold change of enriched proteins for pancreatic endocrine cells based on the Human Protein Atlas single-cell transcriptomic dataset (version 24.1): blue bars represent fold change compared to average expression across all other cell types; red bars show fold change relative to the second-highest expressing cell type. The y-axis shows fold change on the log2 scale. Numbers above bars indicate specific fold change values. The dashed line represents the threshold used for cell type-specific signatures. b, Expression of two example proteins mapped to pancreatic endocrine cells, INS (left) and IAPP (right), across all cell types. c, The corresponding bar plot of fold change for skeletal myocytes. d, Expression of two example proteins mapped to skeletal myocytes, TNNT3 (left) and FST (right), across all cell types.

**Extended Data Figure 3:**
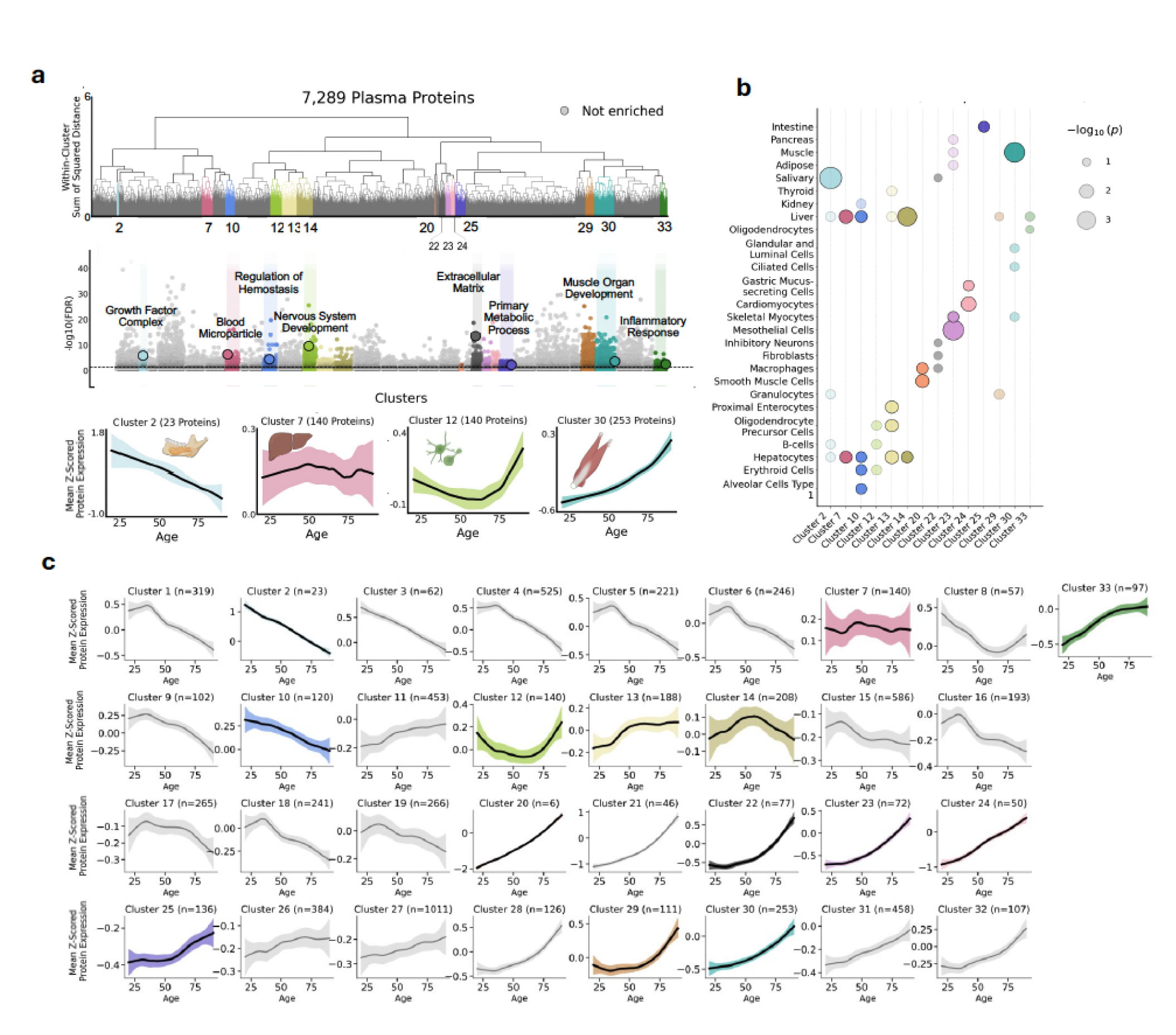
Clustering of plasma protein trajectories. **a**, Expression of 7,289 plasma proteins was measured in 7,074 healthy individuals in GNPC and plotted with age using LOWESS regression. Protein trajectories were clustered using hierarchical clustering, resulting in 33 distinct clusters. Colored clusters indicate significant enrichment for one or more cell types and/or organs. Gray clusters indicate no significant enrichment. b, Trajectory clustering enrichment plots for healthy individuals (n = 7,074). Translucent dots correspond to adjusted p-value < 0.1. Opaque dots correspond to adjusted p-value < 0.05. c, All 33 clusters are shown. The mean LOWESS trajectory of each cluster is plotted as a bolded line with a corresponding shaded 95% confidence interval.

**Extended Data Figure 4:**
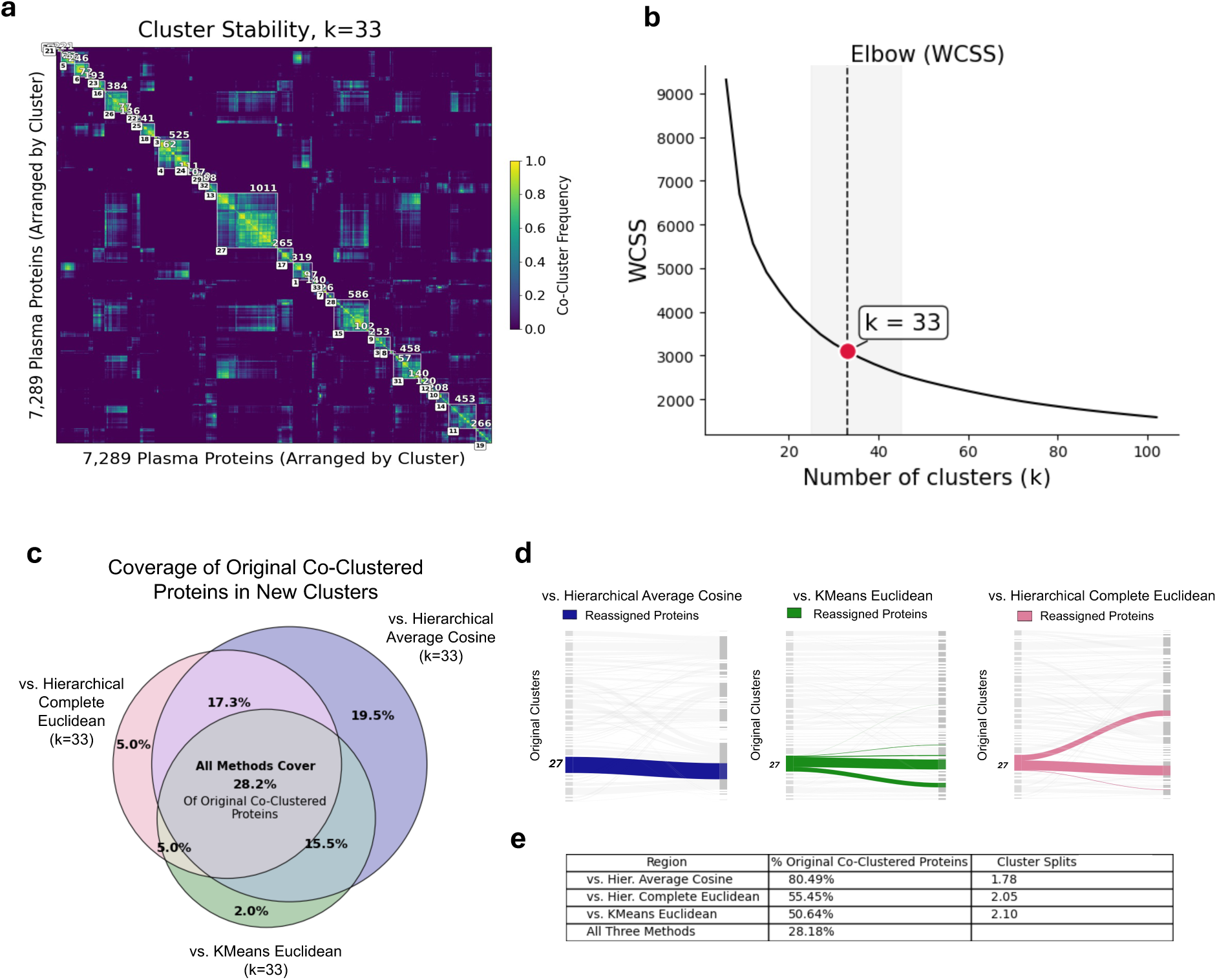
Cluster stability, comparison with other methods, and cluster number rationale. **a**, Cluster stability analysis (k=33) showing within-cluster consensus across 100 bootstrap resamples. Proteins are ordered by original cluster, outlined in white edged blocks, annotated with cluster number (bottom left label) and size (top right label). Higher values indicate more stable cluster assignments. **b**, Elbow plot of within-cluster sum of squares (WCSS) versus number of clusters. The chosen cluster count k = 33 (red) lies in a plateau indicated with a gray band. **c**, Comparison of clustering methods with k=33 clusters. The percentage overlap of co-occurring proteins within clusters is represented for three alternative methods, with Hierarchical Ward Euclidean as the baseline reference. **d**, Distribution of proteins in original clusters within new clusters for each alternative method, with cluster 27 shown as an example. Hierarchical clustering with average linkage method and cosine distance metric shows the highest retention of original co-clustered protein assignments. The mean size-weighted cluster splits for each original cluster were additionally computed.

**Extended Data Figure 5:**
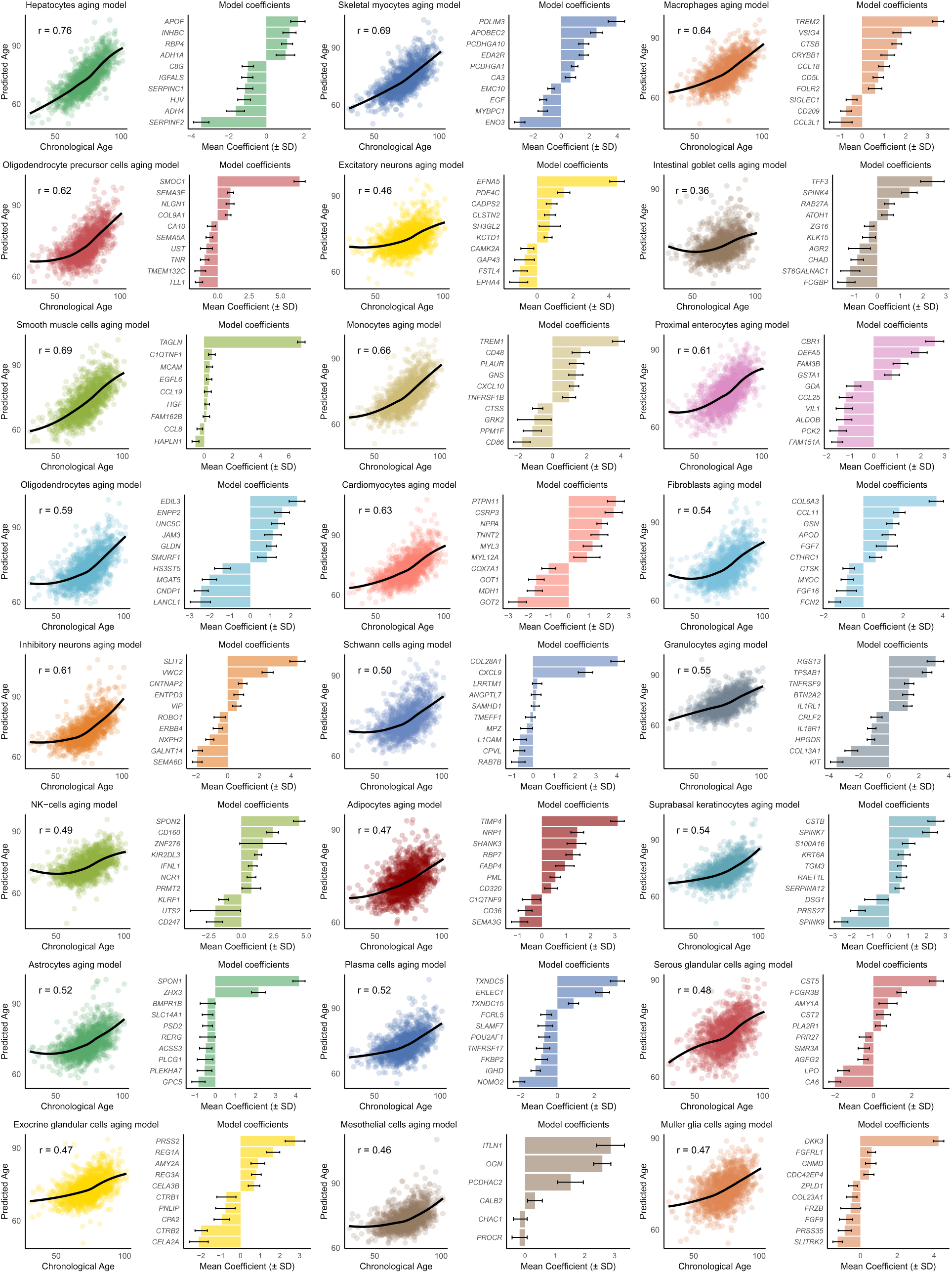
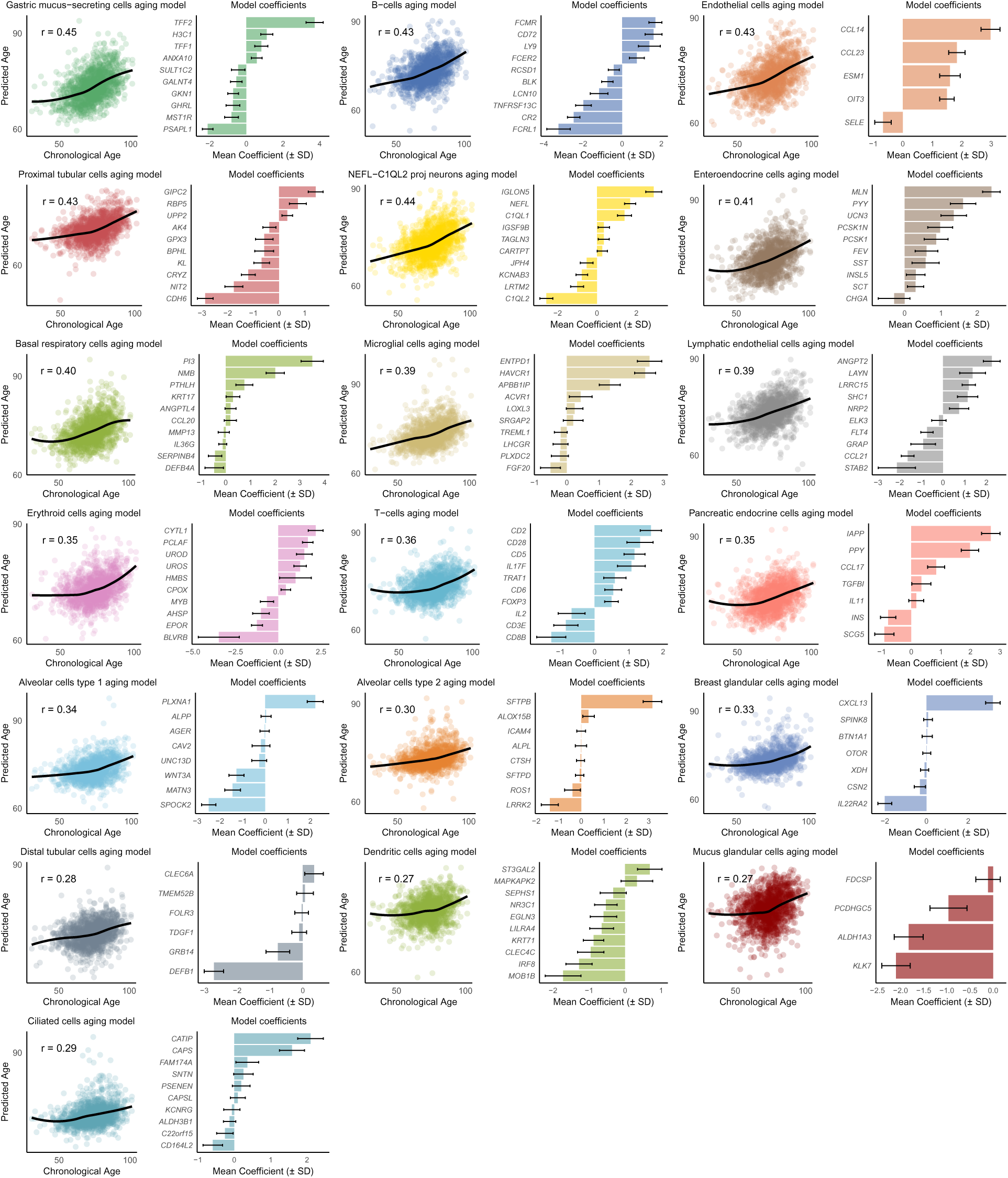
Cell type-specific aging model predictions and core proteins. Scatter plots show cell type-specific aging model predictions versus chronological age with correlation coefficients (r). Bar plots display the mean coefficients (±SD) of the top proteins by absolute coefficients in each cell type-specific aging model. *Cell type-specific aging model predictions and core proteins.* Scatter plots show cell type-specific aging model predictions versus chronological age with correlation coefficients (r). Bar plots display the mean coefficients (±SD) of the top proteins by absolute coefficients in each cell type-specific aging model.

**Extended Data Figure 6:**
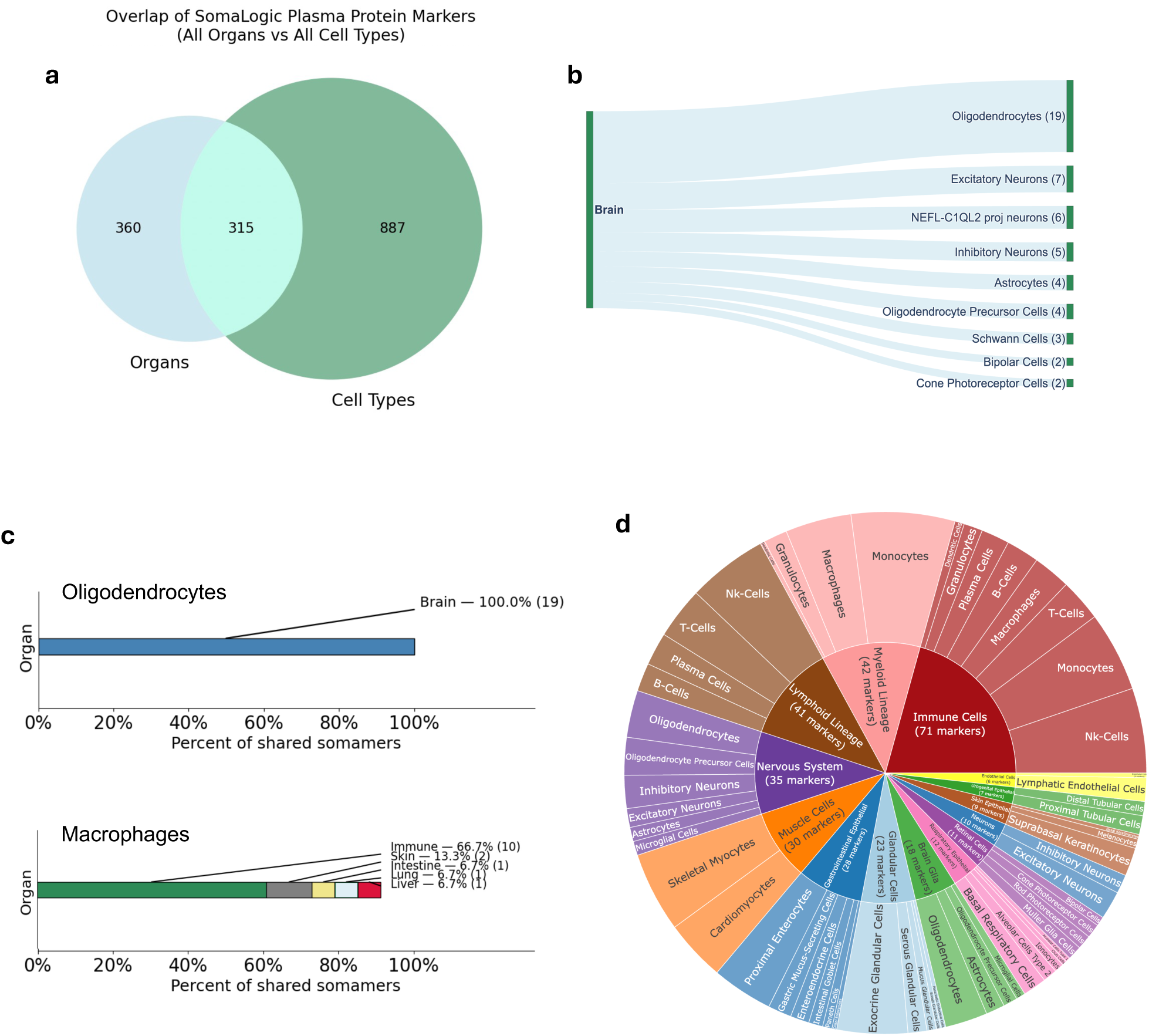
Coverage of putative cell-type plasma proteins measurable with the SomaScan 7k and Olink 3k assays. Cell-type plasma proteins provide granularity beyond organ-level categorizations. **a**, Intersection of putative cellular and organ-enriched plasma proteins measurable with the SomaLogic 7k SomaScan assay. **b**, Example distribution of organ-enriched plasma proteins (n= 18 organs; brain shown) [1] with respect to individual cell types modeled in the study (n=60 cell types). Linkages with two or more shared proteins are shown. **c**, Intersection of cell-type enriched plasma proteins and organ-enriched plasma proteins measurable with the SomaScan assay. Example distribution of oligdendrocyte and macrophage enriched plasma proteins with respect to organs (% Somamers shared) [1]. **d**, Intersection of putative lineage and cell-type plasma proteins measurable with the Olink platform. Only proteins measurable by the Olink assay are shown; a complete list of marker genes and corresponding cell-type lineage members is provided in **Supplementary Table 5** and **Methods**.

**Extended Data Figure 7:**
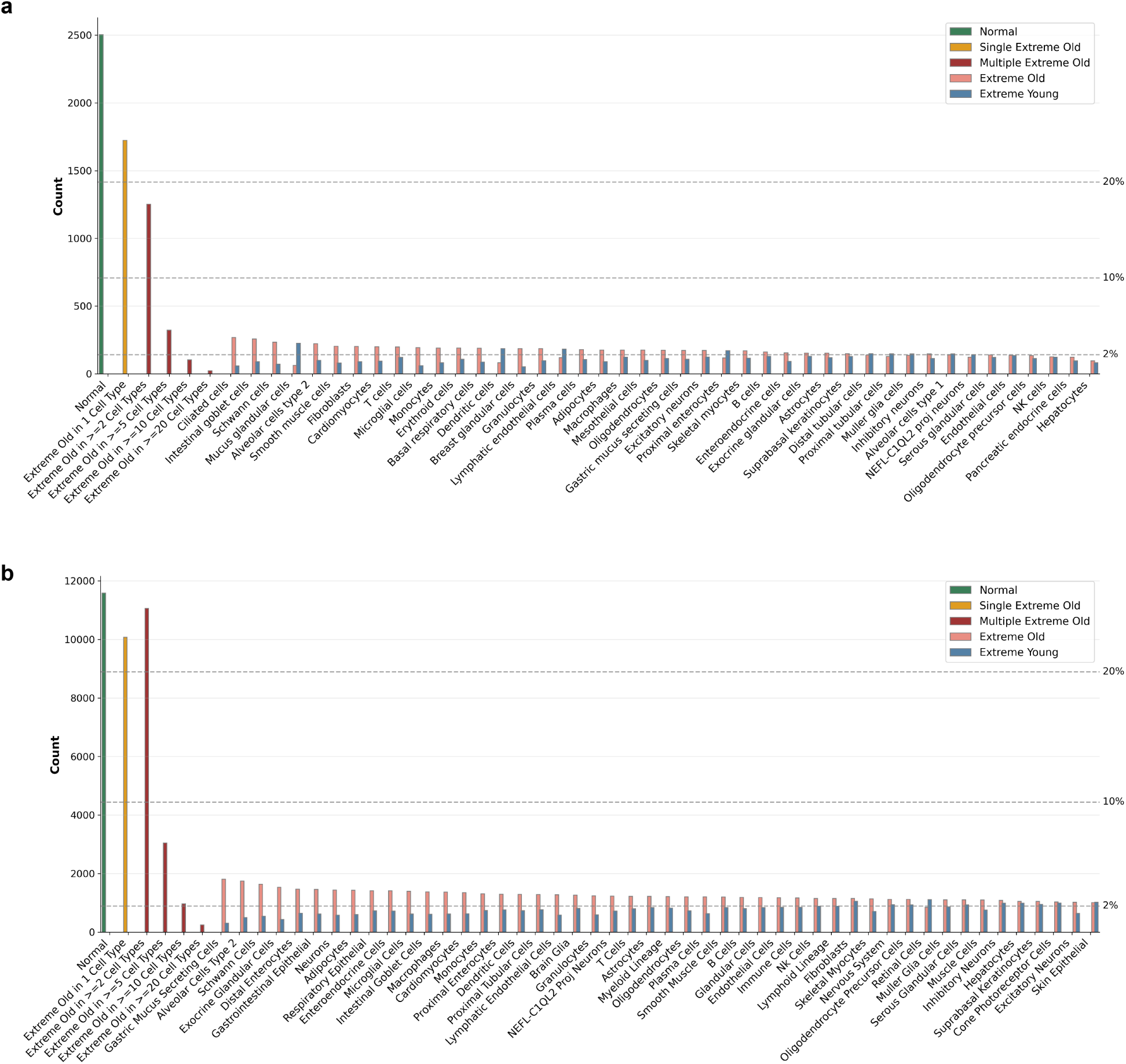
Distribution of extreme aging profiles. **a**, Distribution of extreme aging profiles across cell types among healthy individuals in the GNPC cohort (n=7,074). The bar plot shows the count of individuals categorized by aging status: normal (green, individuals without extreme aging in any cell type), single extreme aging (orange, extreme aging in only one cell type), multiple extreme aging (dark red, extreme aging in multiple cell types), extreme aging by specific cell type (pink, z-scored age gap *>* 2 for the indicated cell type), and decelerated aging by specific cell type (blue, z-scored age gap *< −*2 for the indicated cell type). Horizontal dashed lines indicate reference points at 2%, 10%, and 20% of the cohort. **b**, Distribution of extreme aging profiles across cell types in the UKB cohort (n=44,458). All categorizations, color schemes, and reference lines are as described in (**a**).

**Extended Data Figure 8:**
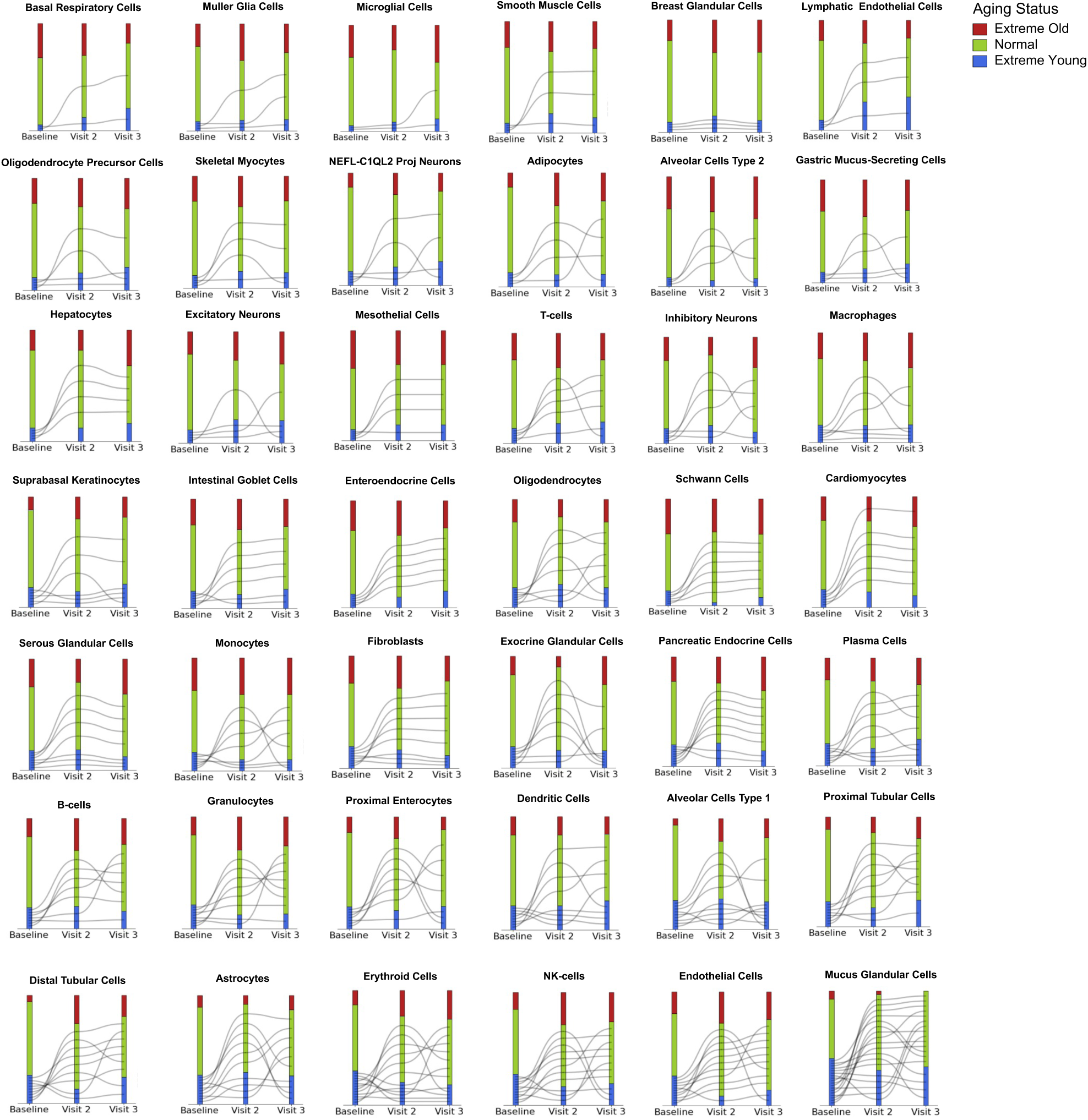
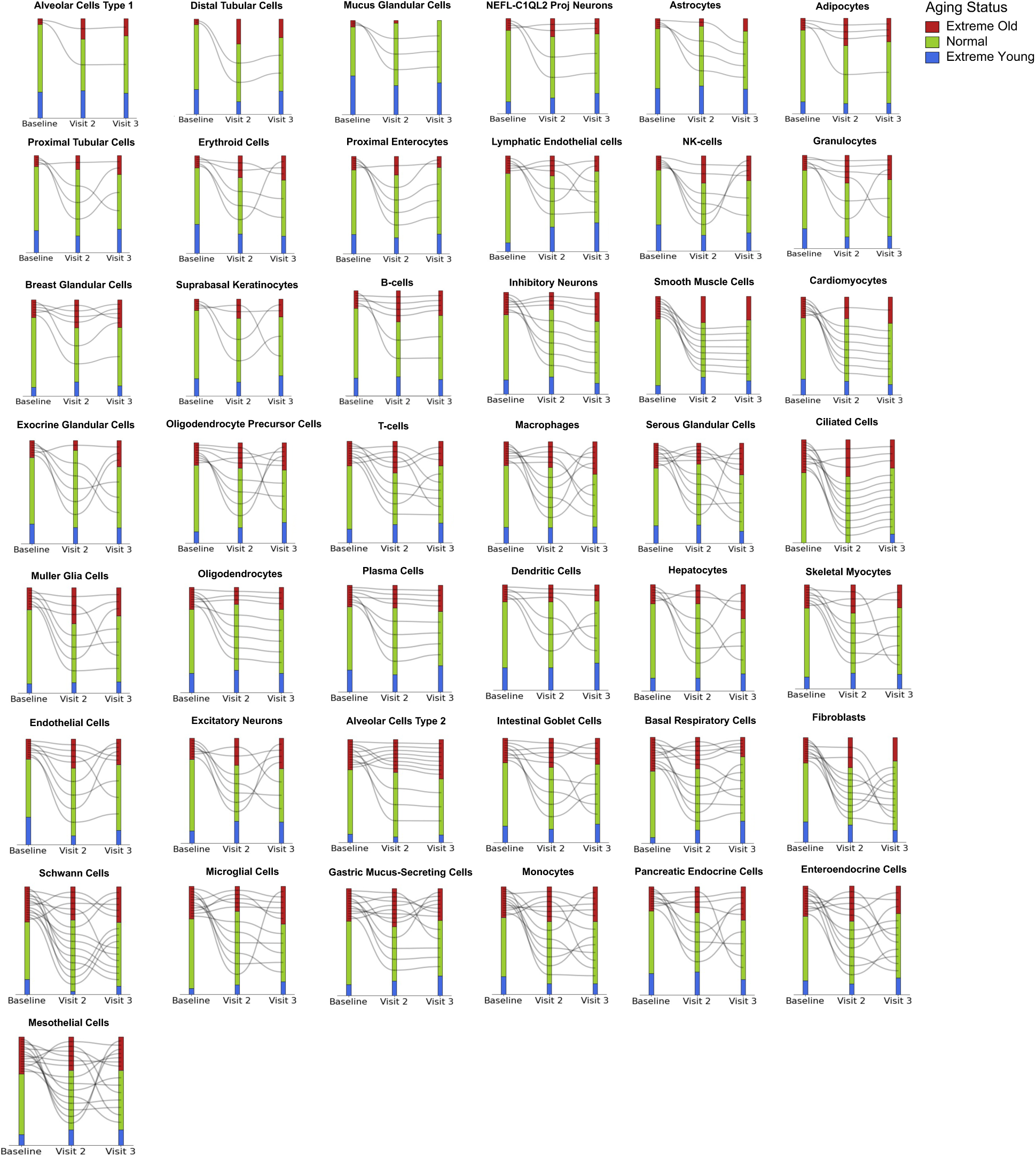
Stability of extreme youth cell-type aging profiles. Cell type aging trajectories for 364 individuals in the NSHD 1946 British Birth Cohort [2] across visits for extreme youthful agers at baseline (z-scored age gap *< −*2). Only cell types with baseline youthful agers are shown. Bin heights for extreme agers (youthful and old) are scaled relative to the cohort-wide totals at each time point to improve visibility, as these groups represent a small percentage compared to normal agers. Cell-types are plotted in order of increasing number of youthful agers at baseline. Timepoints correspond to mean chronological ages in the NSHD cohort: Baseline: 63.2 ± 1.1 years; Visit 2: 70.7 ± 0.7 years; Visit 3: 72.9 ± 0.6 years of age.

**Extended Data Figure 9:**
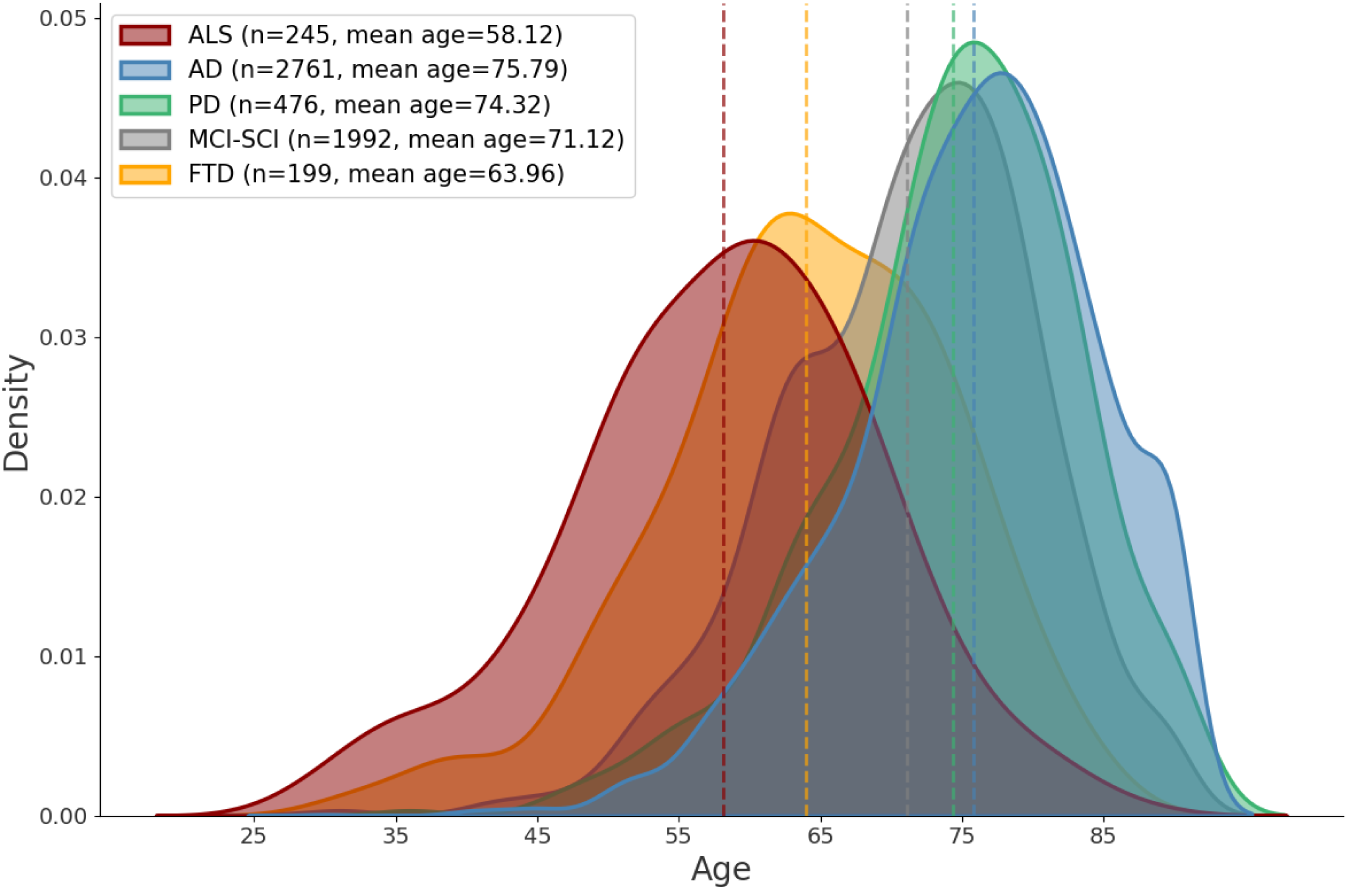
Age distribution across neurodegenerative diseases in GNPC. Density plot showing the chronological age distribution across the five neurodegenerative disease groups in GNPC, with ALS affecting individuals at notably younger ages (mean=58.12 years, s.d.=10.94), followed by FTD (mean=63.96 years, s.d.=10.14), while AD (mean=75.79 years, s.d.=8.83), PD (mean=74.32 years, s.d.=8.84), and MCI-SCI (mean=71.12 years, s.d.=9.11) predominantly manifested at more advanced ages.

**Extended Data Figure 10:**
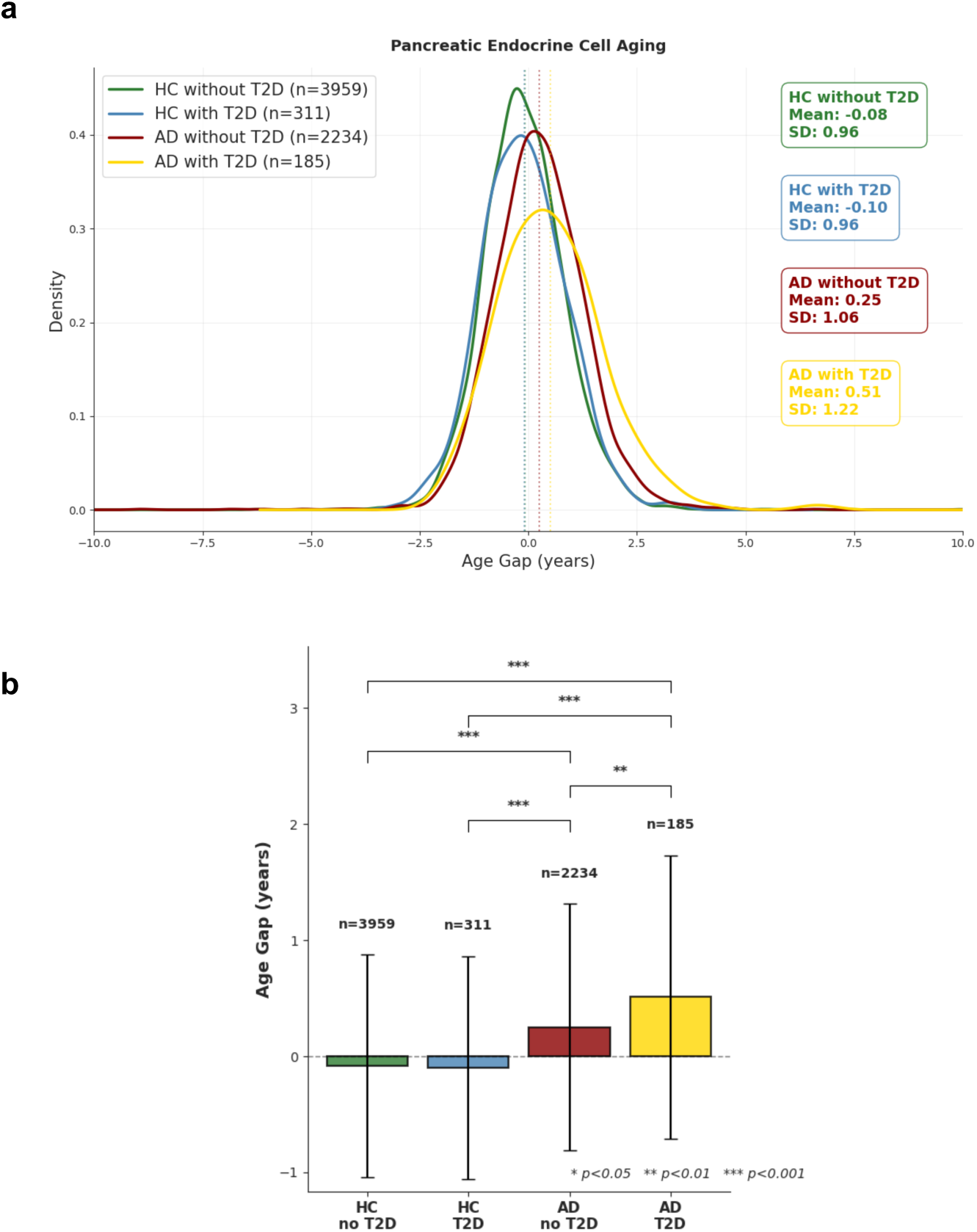
Pancreatic endocrine cell aging in Alzheimer’s disease (AD) with and without type 2 diabetes comorbidity in the GNPC cohort. **a**, Density distributions of pancreatic endocrine cell age gaps across four groups: healthy controls (HC) without T2D (n=3959, mean=-0.08, SD=0.96), HC with T2D (n=311, mean=-0.10, SD=0.96), AD without T2D (n=2234, mean=0.25, SD=1.06), and AD with T2D (n=185, mean=0.51, SD=1.22). Vertical dashed lines indicate mean values for each group. **b**, Comparison of pancreatic endocrine cell age gaps across groups showing mean (horizontal line) and standard deviation (error bars). Sample sizes and p-values from pairwise comparisons are shown (* p<0.05, ** p<0.01, *** p<0.001). AD patients demonstrate accelerated pancreatic endocrine cell aging independent of T2D status, with the effect amplified when both conditions are present.

**Extended Data Figure 11:**
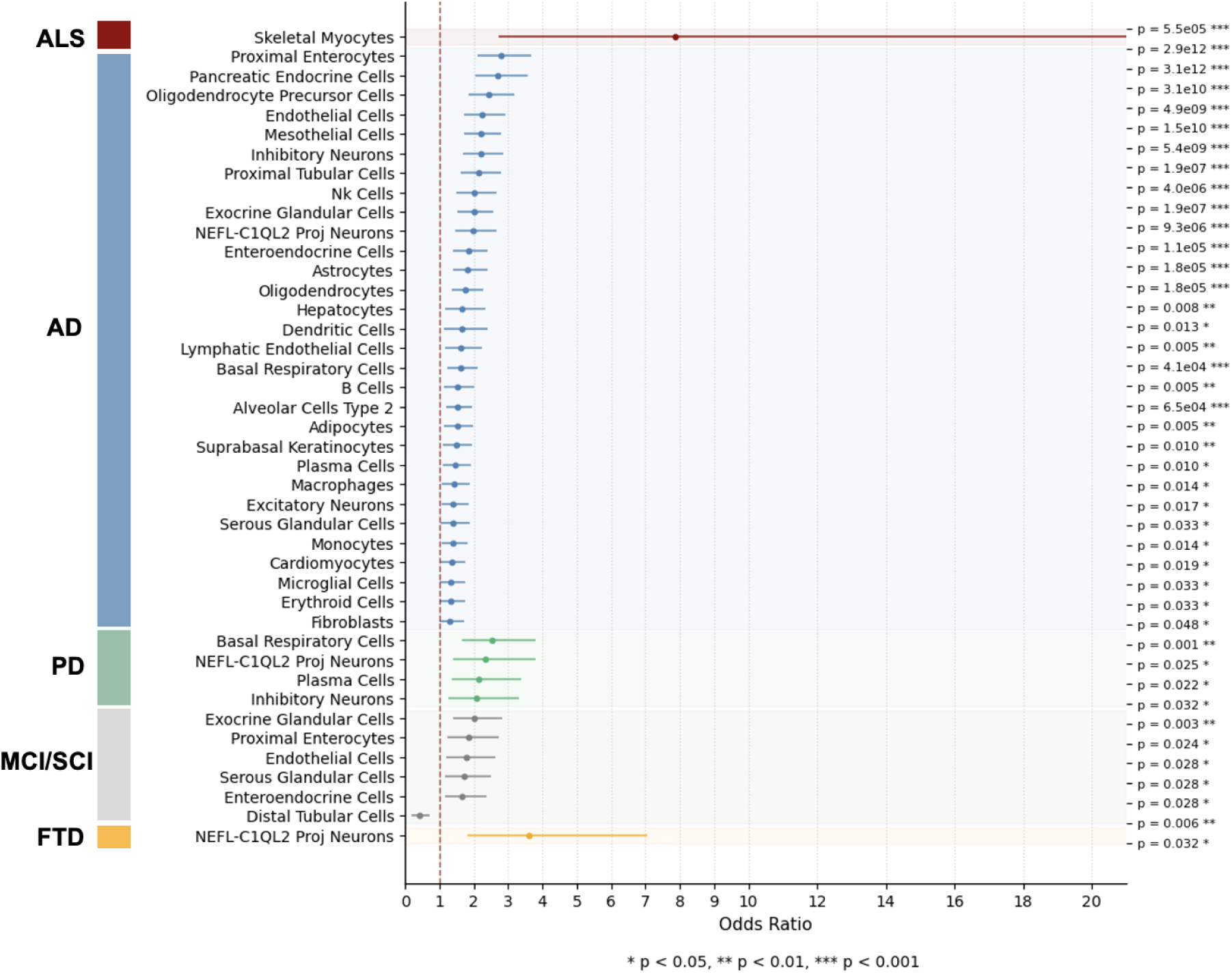
Odds ratio analysis of AD, ALS, PD, FTD, and MCI-SCI in the GNPC cohort. Forest plots showing odds ratios of cell type-specific extreme agers for Alzheimer‘s disease (AD, n=2761), amyotrophic lateral sclerosis (ALS, n=245), Parkinson’s disease (PD, n=476), frontotemporal dementia (FTD, n=199), and mild cognitive impairment/subjective cognitive impairment (MCI-SCI, n=1992). Odds ratios and 95% confidence intervals for significantly associated cell types are shown. All p-values have been adjusted using the Benjamini–Hochberg procedure, with a significance threshold of 0.05. Only significant associations are shown.

**Extended Data Figure 12:**
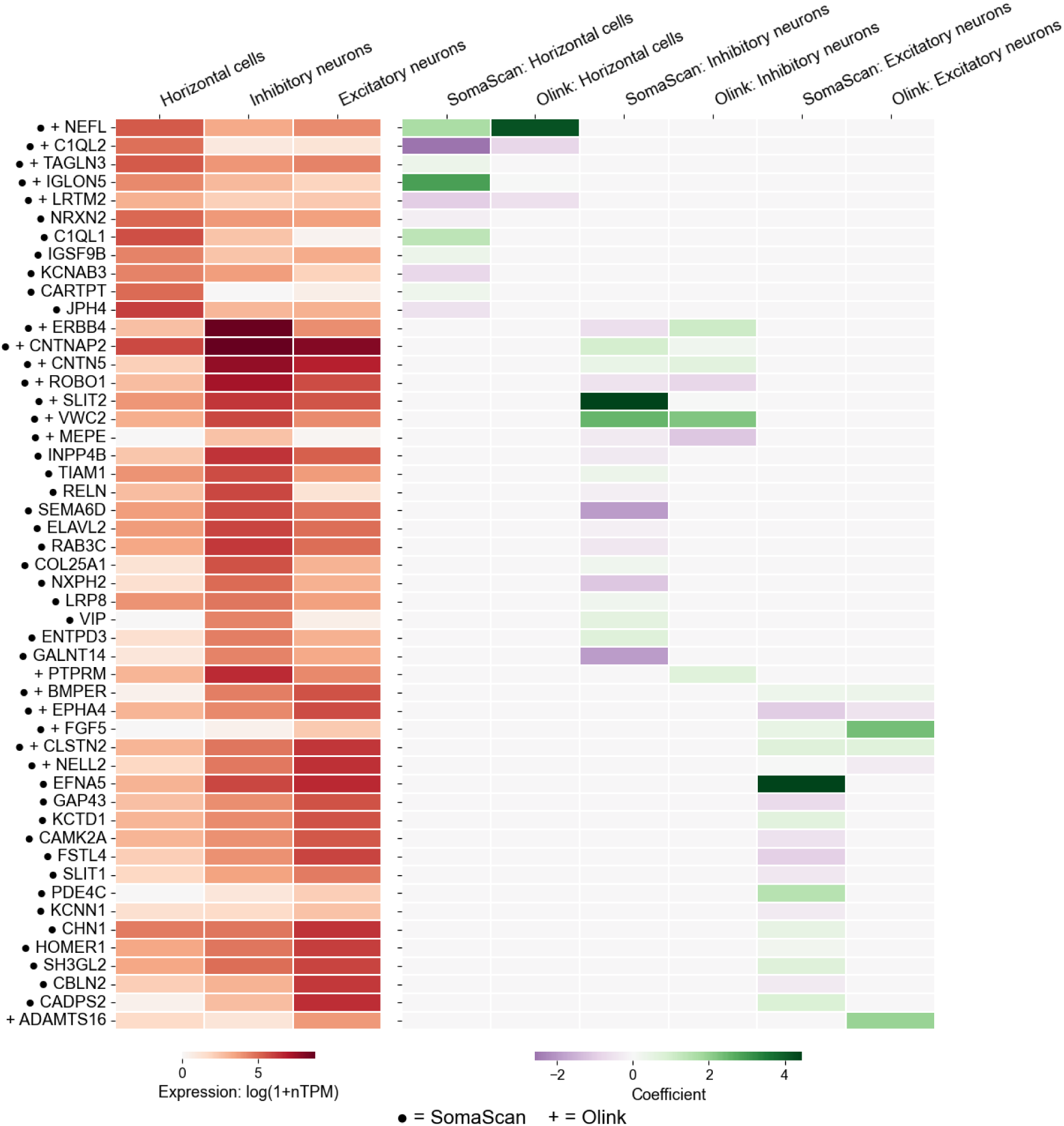
Expression profiles of horizontal cell and neuronal cell signatures and aging model coefficients. Left: Heatmap displays gene expression levels from the Human Protein Atlas single-cell transcriptomic dataset (version 24.1) for signatures of horizontal cells, inhibitory neurons, and excitatory neurons. Color intensity represents expression level, with darker red indicating higher expression. Right: Model coefficients from cellular aging clocks for horizontal cells, inhibitory neurons, and excitatory neurons on SomaScan and Olink platforms. Color represents coefficient magnitude (purple: negative; green: positive); only signatures with an absolute coefficient greater than 0.2 are shown. Circles indicate proteins measured with SomaScan; plus signs indicate Olink coverage. The horizontal cell signature is dominated by NEFL (neurofilament light chain) and C1QL2 (complement C1q-like protein 2), both showing high expression in horizontal cells and strong absolute coefficients in aging models. NEFL is a widely recognized biomarker of axonal injury frequently elevated in FTD, while C1QL2 is a synaptic organizer known to be prominent in temporo-limbic structures vulnerable to fronto-temporal lobar degeneration. The distinct expression and coefficient patterns support renaming this signature to NEFL-C1QL2 projection neuron aging to better reflect its molecular architecture and neurological relevance.

**Extended Data Figure 13:**
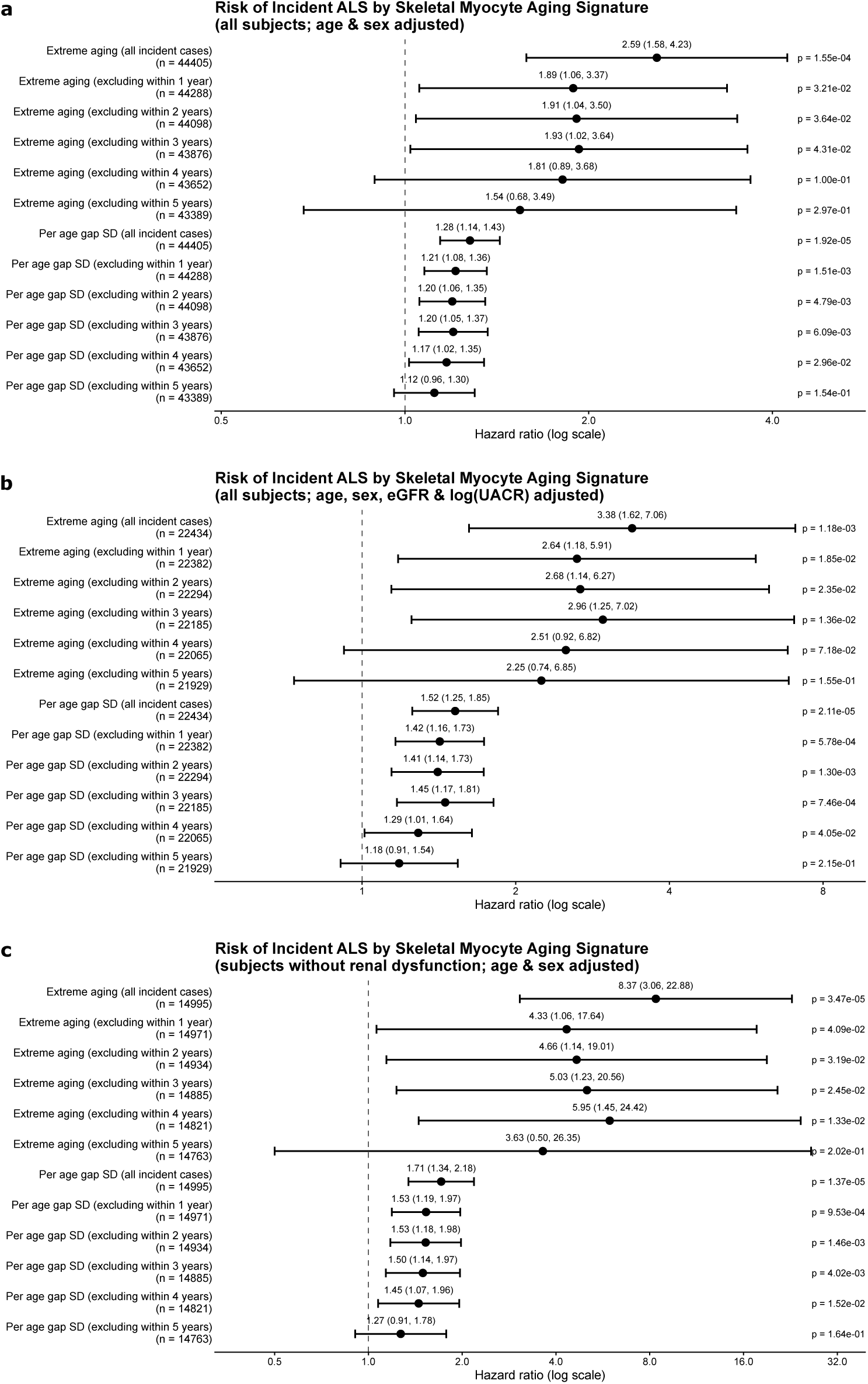
Skeletal myocyte aging is prognostic of incident ALS in age- and sex-adjusted Cox proportional hazards models in UKB, with predictive power retained after excluding cases diagnosed within multiple years of plasma collection. a, Forest plot showing hazard ratios for the skeletal myocyte aging signature, evaluated as both extreme aging status and continuous age gap (z-scored). Models include all incident ALS cases or exclude cases diagnosed within 1, 2, 3, 4, or 5 years after plasma collection to assess predictive power across lag periods. b, Hazard ratios with additional adjustment for estimated glomerular filtration rate (eGFR) and log-transformed urine albumin-to-creatinine ratio (UACR). c, Hazard ratios in a subpopulation restricted to individuals without evidence of kidney dysfunction with estimated glomerular filtration rate *≥* 90mL/min/1.73m^2^ and normal (A1) urine albumin-creatinine ratio (UACR); no adjustment for renal function variables.

**Extended Data Figure 14:**
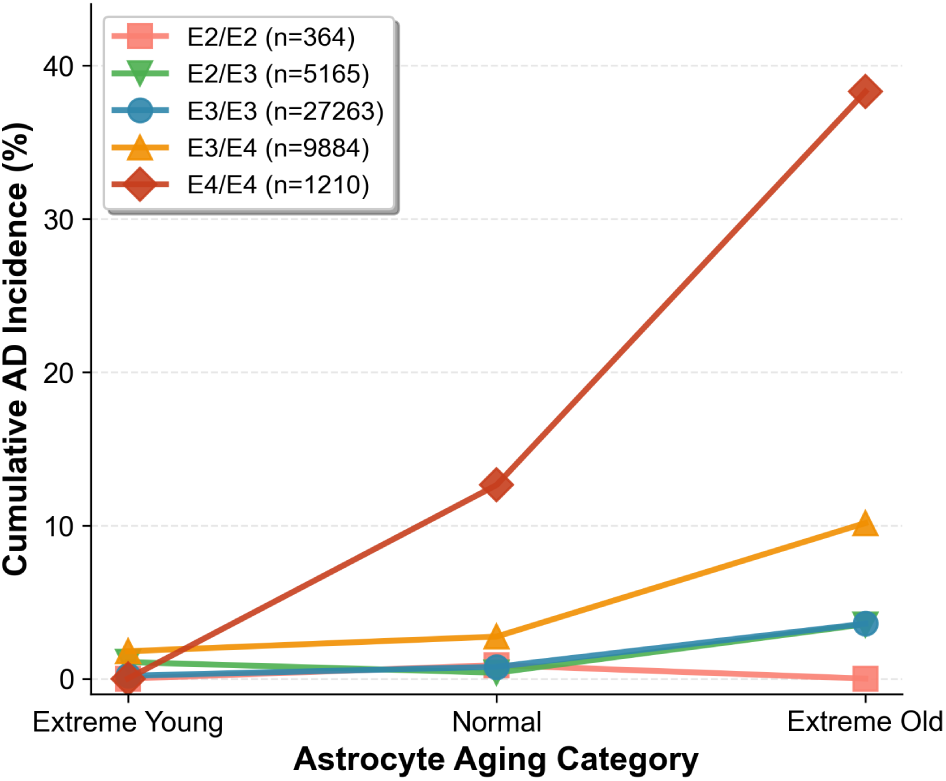
Cumulative AD incidence over 15 years of follow-up in UKB by astrocyte aging status (extreme young, normal, extreme old) and APOE genotype. Lines represent E2/E2 (n=364), E2/E3 (n=5,165), E3/E3 (n=27,263), E3/E4 (n=9,884), and E4/E4 (n=1,210).

**Extended Data Figure 15:**
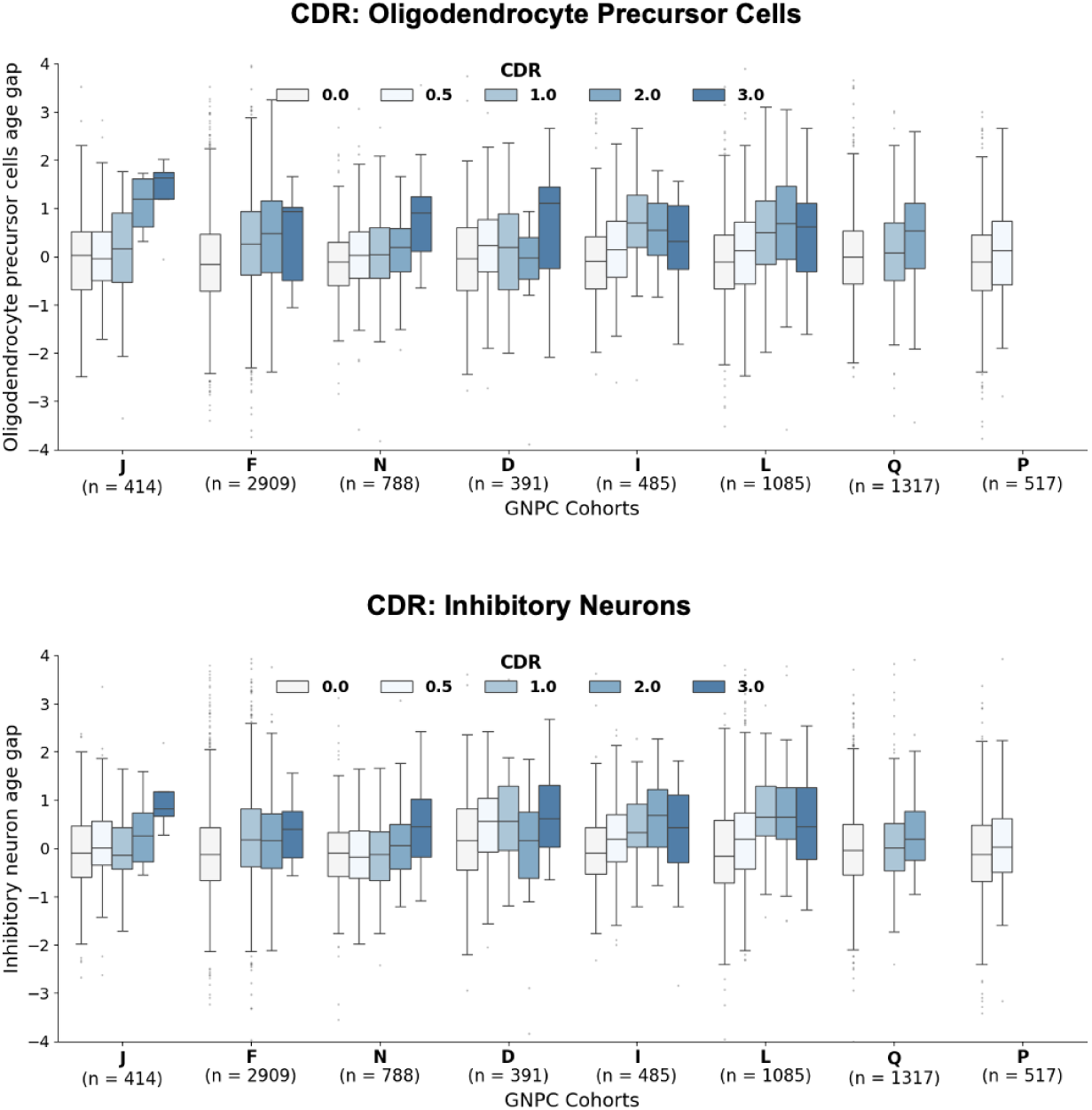
Box plot visualization of age gaps in oligodendrocyte precursor cells and inhibitory neurons stratified by Clinical Dementia Rating (CDR) scores across GNPC cohorts. Oligodendrocyte precursor cells and inhibitory neurons demonstrate the strongest correlation with CDR, with their age gaps showing an overall stepwise increase with worsening cognitive impairment, particularly in cohorts J, F, and N.

**Extended Data Figure 16:**
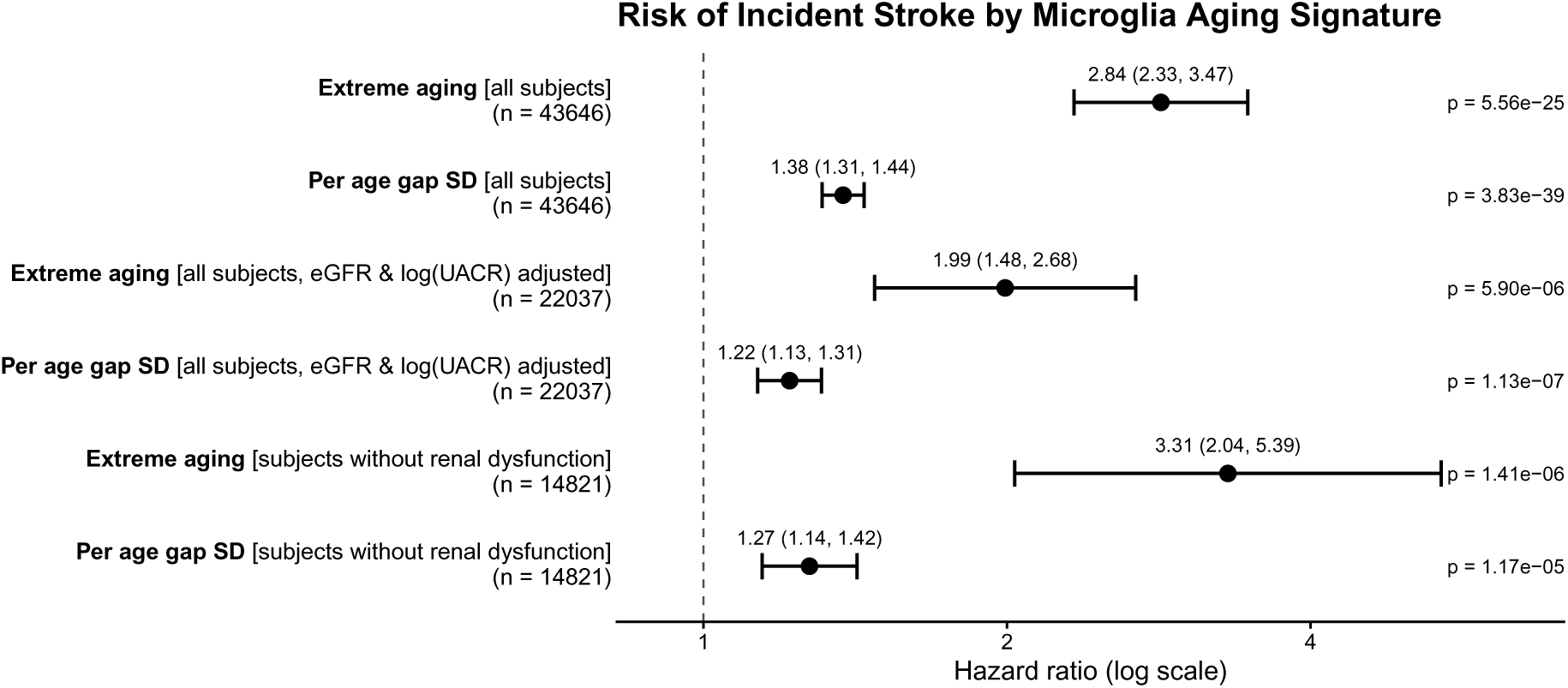
Cox proportional hazards models demonstrate the relationship between stroke risk and the microglial cell aging signature. All models include adjustment for sex and age. Hazard ratios showing the effects of additional adjustment for renal function variables of estimated glomerular filtration rate (eGFR) and log-transformed urine albumin-to-creatine ratio [log(UACR)]—or restriction to individuals without evidence of renal dysfunction (eGFR *≥* 90mL/min/1.73m^2^ and normal A1 albuminuria with UACR <30mg/g). Interestingly, the protein with the largest absolute coefficient in the microglial cell aging model is HAVCR1. HAVCR1 is produced in microglia and, to a lesser extent in renal tubule cells especially in response to injury. In our analysis, after further adjustment for renal function including eGFR and log(UACR) or with restriction of the population to subjects without any signs of kidney impairment, microglial extreme aging remains the second-highest prognostic cell type signature after NEFL-C1QL2 projection neurons for stroke. This finding suggests that the originating cell for the observed relationship between plasma HAVCR1 and stroke risk could be aging microglia. The relationship between plasma HAVCR1 and increased stroke incidence has been previously observed in the Malmö Diet and Cancer cohort (n=4,591, 19.5 years of follow-up), which also reported minimal change after adjustment for kidney function [3].

**Extended Data Figure 17:**
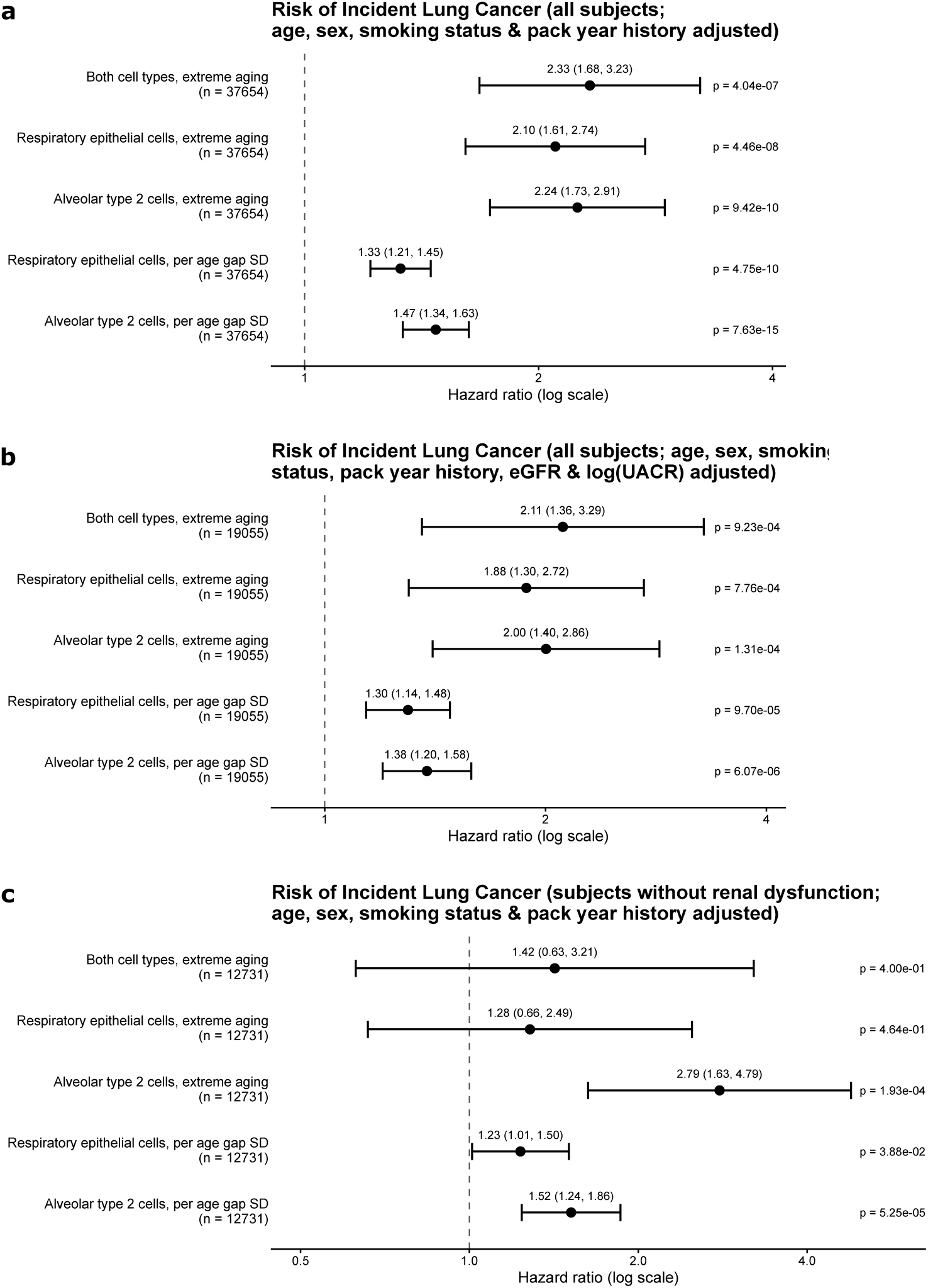
Cox proportional hazards models demonstrate that alveolar type 2 cell and respiratory epithelial cell aging signatures are prognostic of incident lung cancer, after adjustment for age, sex, smoking status, pack-year history, and renal function. **a**, Forest plot showing hazard ratios for the alveolar type 2 cell and respiratory epithelial cell aging signatures, evaluated as both extreme aging status and continuous age gap (z-scored). **b**, Hazard ratios with additional adjustment for estimated glomerular filtration rate (eGFR) and log-transformed urine albumin-to-creatinine ratio (UACR). **c**, Hazard ratios in a subpopulation restricted to individuals without evidence of kidney dysfunction with estimated glomerular filtration rate eGFR *≥* 90mL/min/1.73m^2^ and normal urine albumin-creatinine ratio (A1); no adjustment for renal function variables. Statistical power may be limited by sample size: only 289 subjects had extreme respiratory epithelial cell aging and no evidence of renal dysfunction; only 124 subjects had extreme aging in both cell types and no evidence of renal dysfunction.

**Extended Data Figure 18:**
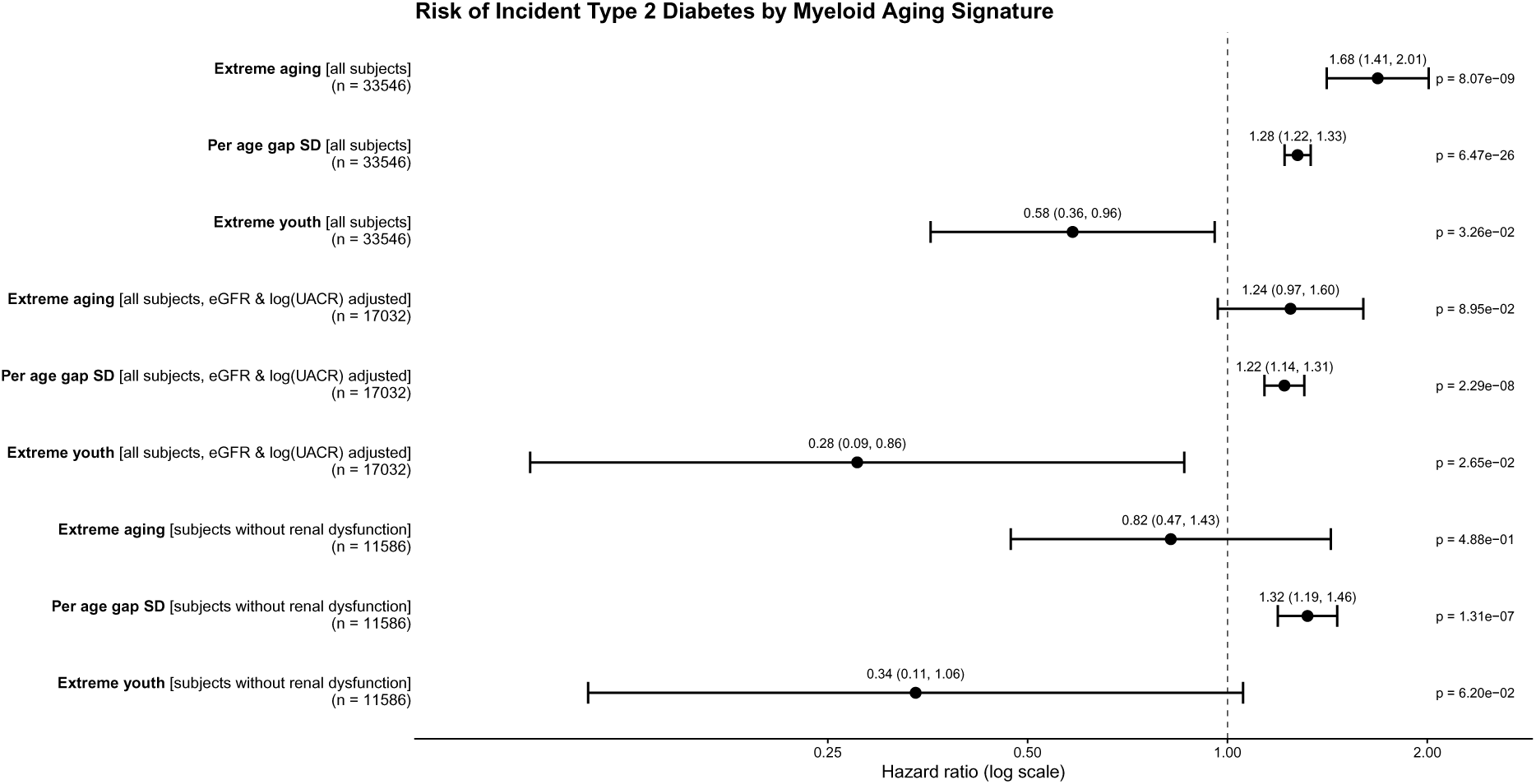
Myeloid lineage aging shows prognostic value for incident type 2 diabetes after adjustment for known risk factors (hemoglobin A1c, body mass index, smoking status, pack year history, sex, and age). Forest plots show hazard ratios per z-scored age gap, for extreme aging, and for extreme youth, in models either further adjusted for renal function based on eGFR and log(UACR) or within the population without evidence of renal dysfunction (eGFR *≥* 90mL/min/1.73m^2^ and normal (A1) urine albumin-creatinine ratio). Power may be limited by small sample sizes (only 118 subjects with both extreme myeloid aging and no evidence of renal dysfunction; only 247 subjects with both extreme myeloid youth and no evidence of renal dysfunction).

**Extended Data Figure 19:**
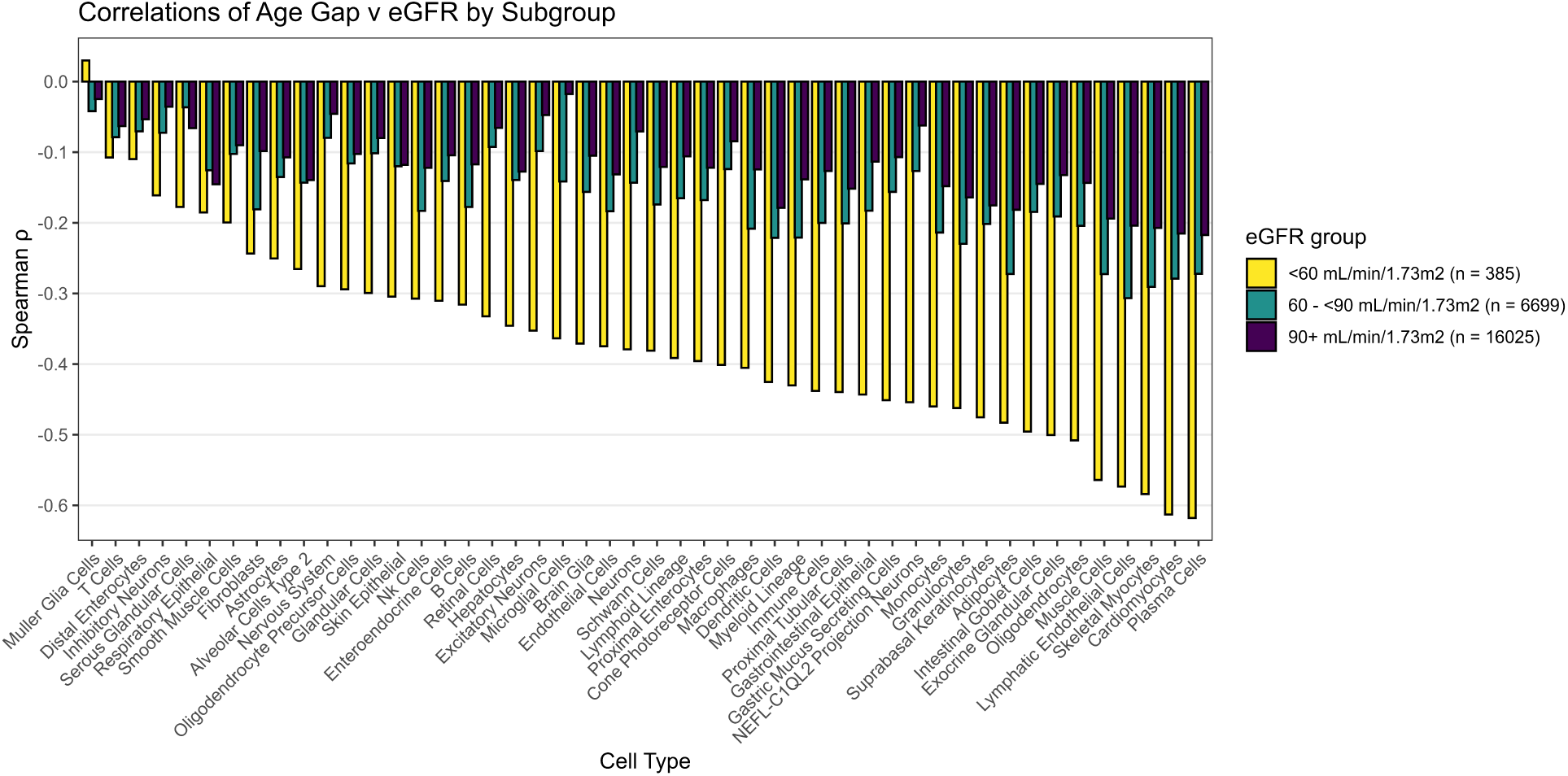
Spearman correlation between age gaps and estimated glomerular filtration rate (eGFR) for each cellular aging clock model, stratified by eGFR ranges. At normal or slightly decreased eGFR (90+ mL/min/1.73m^2^ or 60-90 mL/min/1.73m^2^, which does not meet diagnostic criteria for chronic kidney disease (CKD) unless additional dysfunction is present), the correlation between age gap and eGFR is modest. For subjects with CKD (eGFR < 60 mL/min/1.73 m^2^), there is a strong correlation between the majority of age gaps and renal function, with worsened renal function associated with a more aged plasma signature.

**Extended Data Figure 20:**
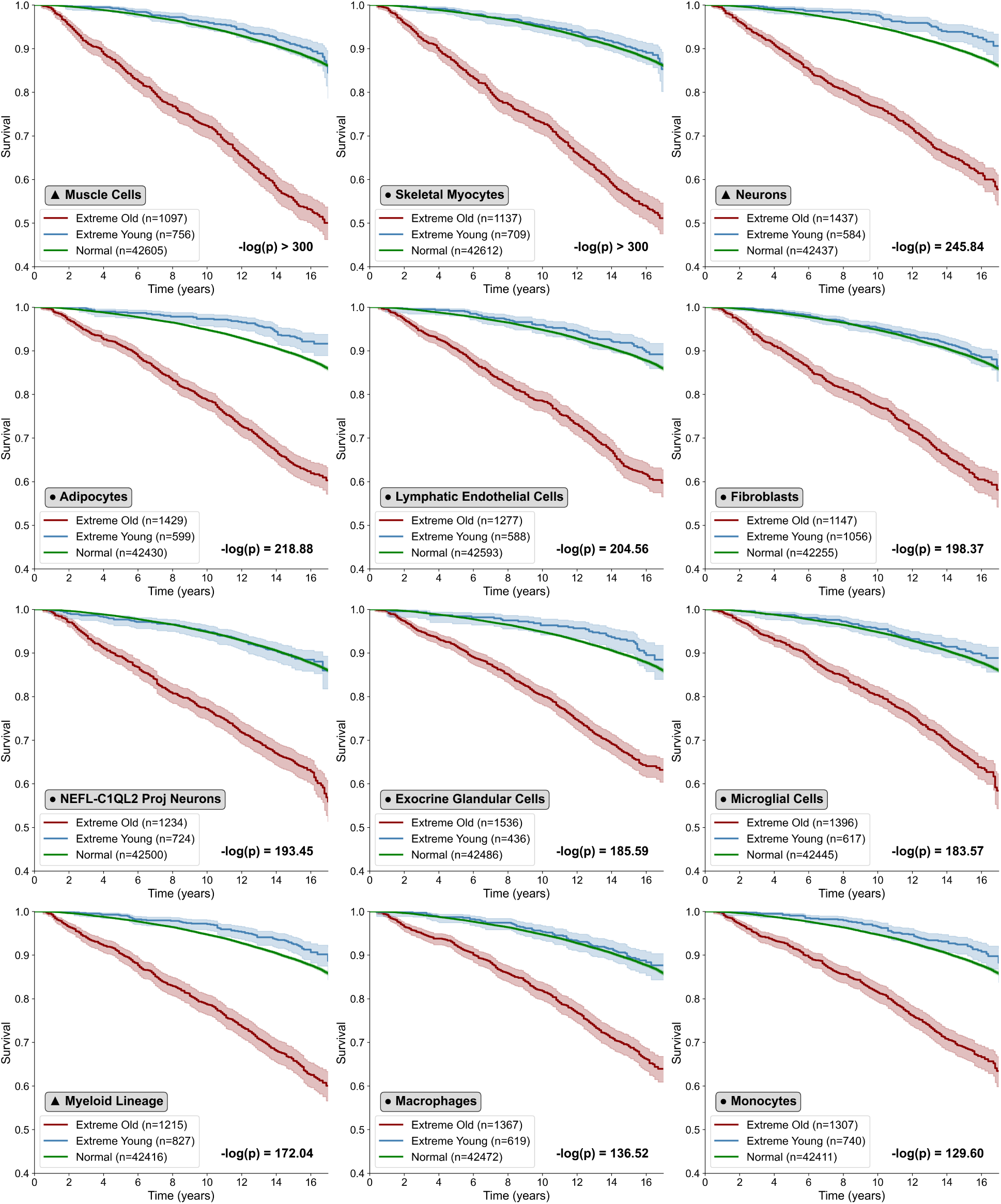
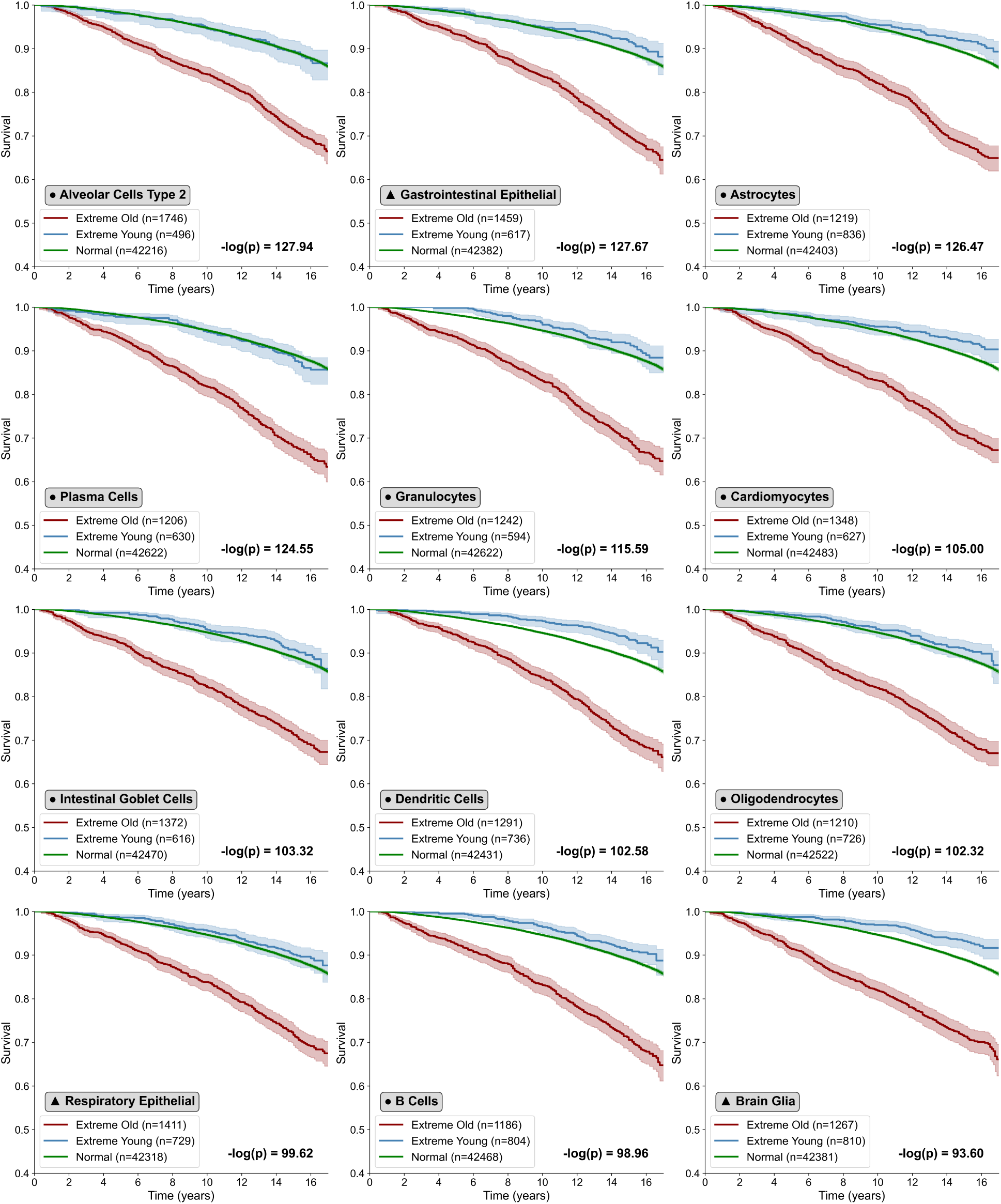
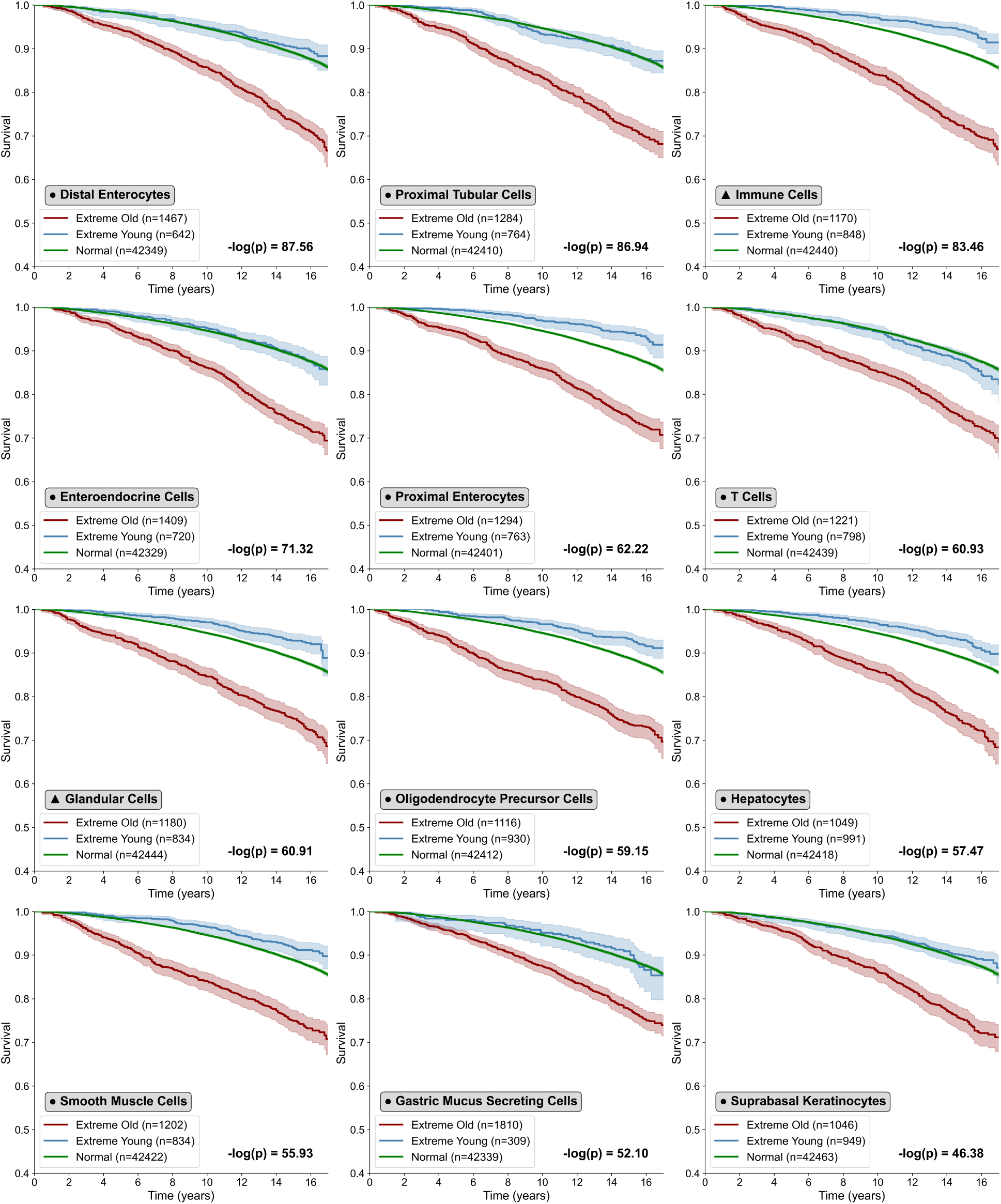
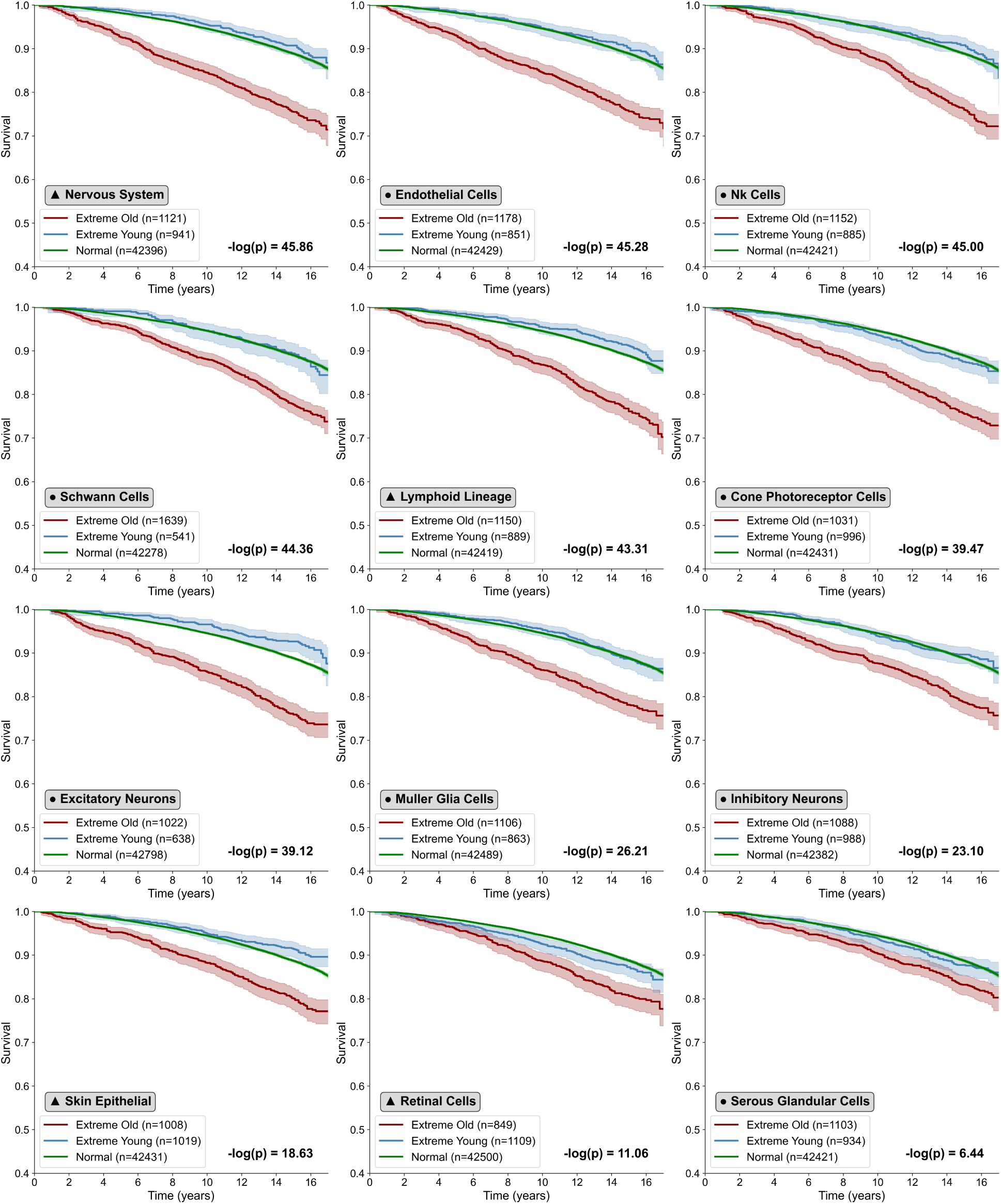
Stratified survival analysis reveals differential mortality risk associated with cellular aging. Kaplan-Meier survival curves showing all-cause mortality stratified by cellular aging status across cell types in the UK Biobank (n = 44,458). Participants are categorized into three groups: extreme old (red), extreme young (blue), and normal aging (green), with sample sizes annotated for each group. Each panel represents a distinct cell type, with circles indicating individual cell types and triangles indicating lineage cell types. Adjusted p-values are displayed as -log(p), where statistical significance was assessed using the log-rank test with Benjamini-Hochberg correction for multiple testing.

**Extended Data Figure 21:**
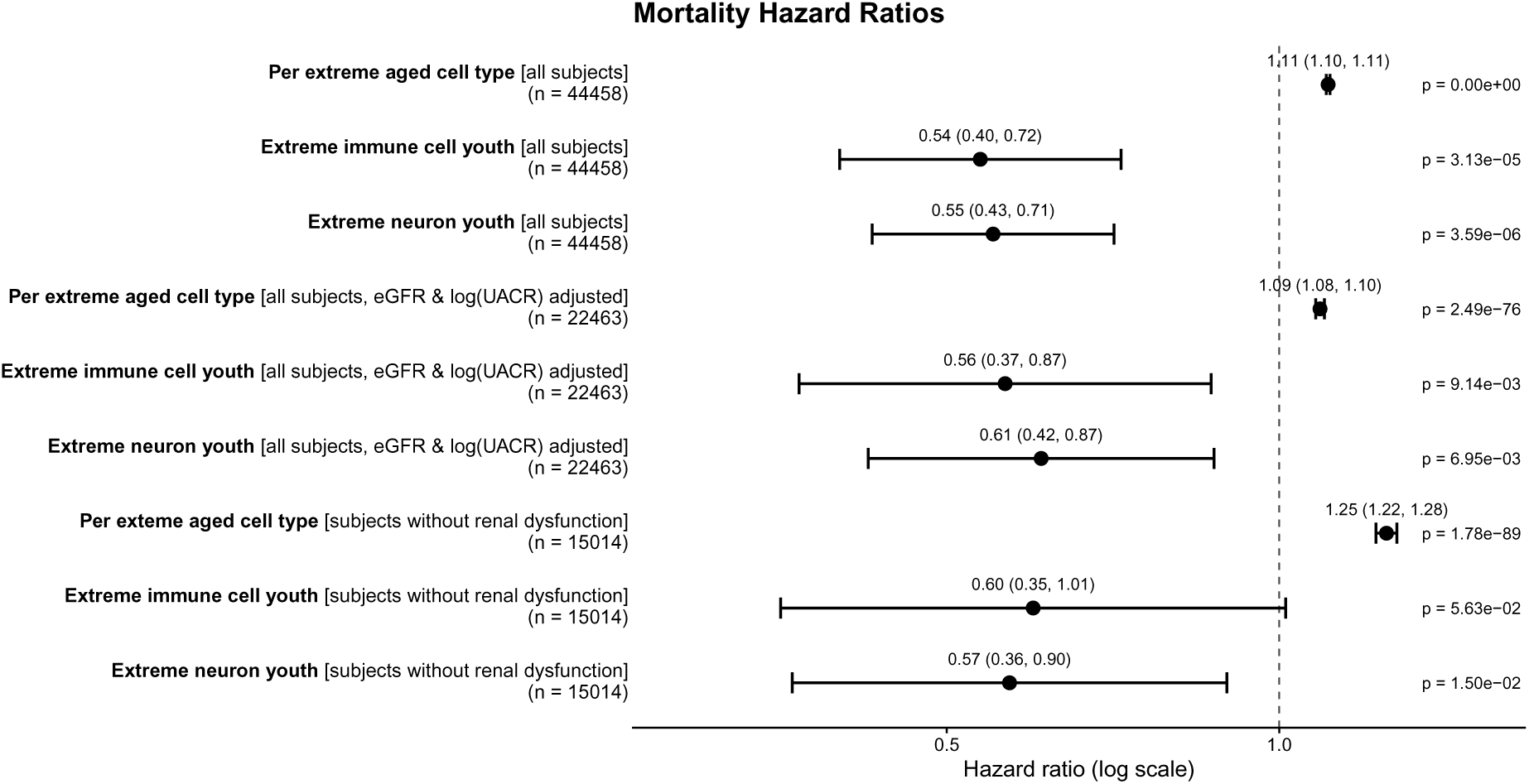
Cox proportional hazards models demonstrate the impact of bearing an additional extreme aged cell type on mortality risk and evaluate the protective effect associated with extreme youth of immune cells and neurons. All models include adjustment for sex and age. Hazard ratios are shown with additional adjustment for renal function variables of estimated glomerular filtration rate (eGFR) and log-transformed urine albumin-to-creatine ratio) [log(UACR)] or with restriction to individuals without evidence of renal dysfunction (eGFR *≥* 90mL/min/1.73m^2^ and normal A1 albuminuria with UACR <30mg/g).

